# Consensus methods for trees of blobs and level-1 phylogenetic networks

**DOI:** 10.64898/2026.10.31.763369

**Authors:** Kristina Wicke, Hector Baños, Niels Holtgrefe, Elizabeth S. Allman, Cécile Ané, Olivier Gascuel

## Abstract

Phylogenetic networks are increasingly used to represent evolutionary histories involving hybridization, introgression, and other reticulate processes that cannot be captured by a single tree. Recent advances have produced numerous methods for inferring networks from genomic data. These methods may produce multiple plausible networks due to optimality ties, bootstrap analyses, Bayesian posterior samples, or differences in tools, datasets, and parameters. While consensus methods are well established for trees and some approaches exist for rooted networks, no consensus framework currently exists for semi-directed (unrooted) networks. We address this gap by introducing a new consensus framework for phylogenetic networks, rooted or semi-directed. We first develop a method for constructing a consensus tree of blobs, taking as input networks of any complexity. Building on this approach, we construct a consensus for level-1 networks (i.e., networks with non-overlapping cycles). This method, called ‘L1N consensus’, takes a collection of level-1 networks as input and returns a level-1 network. The different components of this consensus network (e.g., blobs, hybrid clades, and edges) have support values that correspond to the frequency of the given component among the input networks. These methods are implemented in PhyloSummaries, building on the foundation of PhyloNetworks. We evaluate the performance of L1N consensus using large-scale simulated data. Our results demonstrate that L1N consensus outperforms existing methods in the rooted setting in terms of informativity and accuracy. Similar results are obtained in the semi-directed setting. Applying L1N consensus to Leopardus and HIV datasets confirms these findings and demonstrates the usefulness of the method.

## 1 Introduction

Reconstructing the evolutionary history of species is a central problem in biology. Although phylogenetic trees have traditionally been used for this purpose, advances in genome sequencing have made it increasingly clear that evolution is often not strictly tree-like. Processes such as recombination, hybridization, and lateral gene transfer can produce reticulate patterns of ancestry, and phylogenetic networks have therefore become essential tools for representing and inferring such complex evolutionary scenarios.

Phylogenetic networks come in several forms, but two representations are especially relevant for empirical analyses. See Figure 1 for an illustration, and the Supplementary Material for formal definitions. A *rooted phylogenetic network* resembles a rooted tree: edges are oriented forward in time, and reticulation events are represented by nodes with two incoming edges, indicating contributions from two ancestral lineages. This representation is biologically intuitive but often difficult to infer reliably in practice, in part because the root position is often not identifiable under common evolutionary models. Consequently, many inference tools instead produce a *semi-directed phylogenetic network*. Mathematically, such a network is obtained from the rooted version by suppressing the root and removing directions on all tree-like edges while preserving the orientation of hybrid edges. The resulting structure is unrooted, analogous to an unrooted phylogenetic tree, yet still encodes the direction of gene flow or parental contributions through its directed hybrid edges. In addition, all edges below hybrid nodes can be oriented from past to present. A semi-directed network can be rooted to recover a rooted phylogenetic network when an outgroup is available, provided that the outgroup is not below any hybrid node.

**Figure 1:**
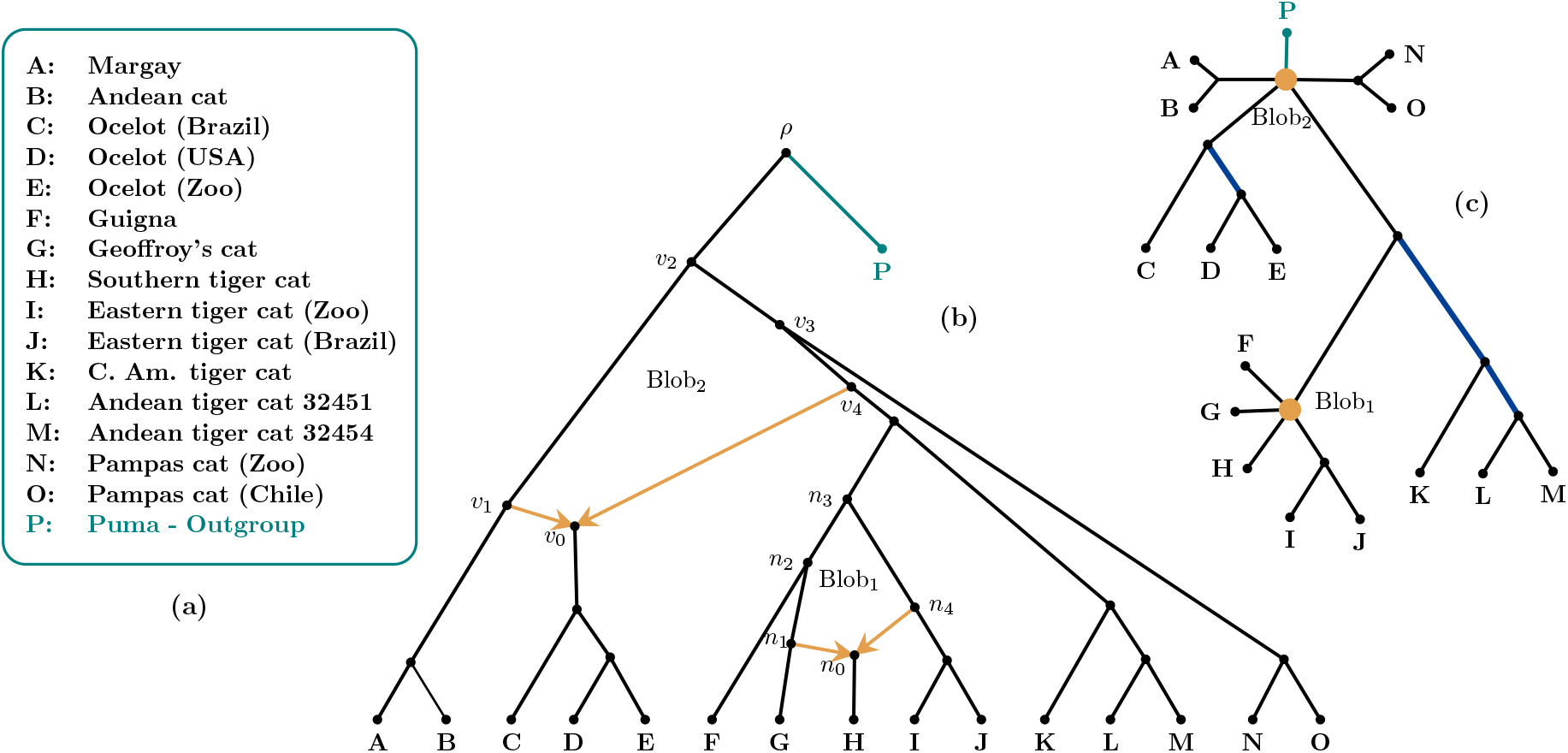
Level-1 phylogenetic network of Neotropical small felids in the genus *Leopardus*. (Lescroart et al., 2023). **(a)** Species and/or individuals; Puma is the outgroup. **(b)** Phylogenetic network with two hybridization events, each associated with two hybrid (or reticulation) edges, in orange. These events give rise to two hybrid clades, {*C, D, E*} and {*H*}, one of which, {*H*}, contains a single taxon. Other network nodes and edges describe speciation and divergence events and strictly vertical inheritance, with interpretations similar to those in phylogenetic trees. Each hybridization event defines a cycle with circular order: (*v*_0_, *v*_1_, *v*_2_, *v*_3_, *v*_4_) and (*n*_0_, *n*_1_, *n*_2_, *n*_3_, *n*_4_). In this network, the two cycles do not share any nodes or edges, making this an example of a level-1 network. Each cycle reveals the two origins of its hybrid clade, here ancestors *v*_1_, *v*_4_ for {*C, D, E*}, and *n*_1_, *n*_4_ for {*H*}. Without an outgroup specified, software programs typically infer semi-directed networks (Baños, 2019; Solís-Lemus and Ané, 2016), which means that only the hybrid edges are directed. In fact, edges below hybrid nodes are implicitly oriented from past to present. Here, Puma (*P*, in teal) is an outgroup, and all edges are directed from past to present away from the root.**(c)** Tree of blobs obtained from (b) by suppressing the root and contracting cycles. Blobs in the network correspond to orange blob nodes in the ToB, and edges inducing non-redundant splits are shown in blue. Note that these orange blob nodes have a different interpretation from polytomies in phylogenetic trees. They signify that one (or more) hybridization(s) occurred in the underlying network, but convey no information about the underlying blob. Specifically, for a blob node in a network’s ToB, there is no information about which taxon group is of hybrid origin, nor about the circular ordering of subgraphs around the blob node. Since current methods (Allman et al., 2024b; Dai et al., 2026; Rhodes et al., 2026a) typically infer undirected trees of blobs, only the inclusion of an outgroup makes it possible to direct edges from past to present.

Over the past decades, phylogenetic network inference has seen substantial theoretical and methodological development, resulting in a growing suite of tools for estimating networks from genomic data. Existing approaches span a broad methodological spectrum. Combinatorial methods, such as FHyNCH (Bernardini et al., 2024) and PhyloFusion (Zhang et al., 2025), reconstruct networks by amalgamating collections of gene trees. Likelihood-based approaches rely on explicit evolutionary models, accounting for incomplete lineage sorting (ILS) with the multispecies coalescent (MSC), but tend to be computationally prohibitive. For example, such methods are available in PhyloNet (Wen et al., 2018). Others include Legofit (Rogers, 2019), which evaluates the likelihood of a single candidate network at a time, and NetRAX (Lutteropp et al., 2022), which ignores ILS, limiting its biological realism. Bayesian methods also tend to be computationally expensive. Implementations include PhyloNet (Wen and Nakhleh, 2017; Wen et al., 2016; Zhu et al., 2018), BEAST2 (Rabier et al., 2021; Zhang et al., 2017), AdmixtureBayes (Nielsen et al., 2023), and BPP (Flouri et al., 2019) which focuses on a candidate network. Pseudo-likelihood methods such as SNaQ (Solís-Lemus and Ané, 2016), PhyNEST (Kong et al., 2024), TreeMix (Pickrell and Pritchard, 2012), and find graphs (in ADMIXTOOLS 2, Maier et al., 2023) also account for ILS while remaining computationally scalable compared to full-likelihood approaches (frequentist or Bayesian). NANUQ (Allman et al., 2019) combines statistical and combinatorial principles for coalescent-based inference. Conceptually different, TINNiK (Allman et al., 2024b), TOB-QMC (Dai et al., 2026), and ECToBlob (Rhodes et al., 2026a) do not directly infer a full phylogenetic network. Instead, they estimate the associated *tree of blobs* (ToB), collapsing reticulate structures into single nodes while preserving the underlying tree-like branching (Fig. 1). NANUQ^+^ (Allman et al., 2025) and NetCS (Dai and Molloy, 2026) build on this idea, first inferring a tree of blobs and then resolving it into a network under a coalescent framework. Relatedly, Squirrel (Holtgrefe et al., 2025b), though not coalescent-based, constructs a sequence of tree-of-blobs-like structures and infers networks from complete sets of quarnets, or 4-leaf subnetworks.

Despite differences among methods and assumptions, phylogenetic network analyses frequently yield multiple plausible solutions. Such variability may arise from ties among equally well-supported networks, bootstrap resampling, posterior distributions from Bayesian analyses, discrepancies across inference methods or datasets, or even differences in parameter settings within a single software package. As for phylogenetic tree inference, it is therefore desirable to summarize the shared structural features of a set of networks into a single representative consensus network.

Consensus procedures for phylogenetic trees are well established (Bryant, 2003). For rooted phylogenetic networks, a consensus method was recently introduced by Huber et al. (2023). This consensus method, which we refer to as the HMS (Huber-Moulton-Spillner) consensus, is based on an encoding of so-called 1-nested networks as collections of ordered pairs of subsets of the taxon set. The HMS consensus only applies to rooted phylogenetic networks and is not applicable to the semi-directed setting. Furthermore, the authors did not provide a practical implementation. We therefore developed a basic Perl implementation to facilitate comparison with our approach. As we demonstrate, the HMS consensus often yields consensus networks that are less resolved and less informative than those produced by our method.

Another consensus method was implemented in AdmixtureBayes (Nielsen et al., 2023) to summarize a Bayesian sample of rooted phylogenetic networks. We refer to this method as HWC (hardwired cluster) consensus, as it is based on the *hardwired clusters* of the input networks, defined as the set of descendant leaves of their nodes (Huson et al., 2010). As with HMS, our results indicate that HWC often produces less informative consensus networks than our approach. Moreover, the current implementation of HWC in AdmixtureBayes limits its practical applicability (see below for details). To address these limitations and facilitate direct comparison with our method, we also provide a basic Perl implementation of the HWC consensus.

Importantly, no consensus method has yet been developed for semi-directed phylogenetic networks, which are commonly inferred by most methods. For instance, while SNaQ (Solís-Lemus and Ané, 2016; Solís-Lemus et al., 2017) implements bootstrap analyses, it does not natively produce a bootstrap consensus network. Instead, PhyloNetworks can map support values onto a reference network such as the inferred network. Similarly, the SummarizeNetworks command in PhyloNet can identify shared components across networks but does not construct an explicit consensus network.

The goal of this paper is to address the current lack of consensus approaches for phylogenetic networks. Phylogenetic networks can be classified by their level, which captures the complexity of their reticulate structure. Level-1 networks are those in which the cycles corresponding to reticulation events do not overlap (Figure 1). This class is particularly relevant in practice, as several widely used network-inference methods, including SNaQ (Solís-Lemus and Ané, 2016; Solís-Lemus et al., 2017), PhyNEST (Kong et al., 2024), NANUQ (Allman et al., 2019), NANUQ^+^ (Allman et al., 2025), and Squirrel (Holtgrefe et al., 2025b) infer level-1 networks. The popularity of these software tools and the level-1 assumption is based on theoretical results demonstrating identifiability properties (Allman et al., 2024a; Baños, 2019; Gross et al., 2021; Solís-Lemus and Ané, 2016; Warnow et al., 2025), while there are few theoretical guarantees for more complex network classes (Allman et al., 2023; Englander et al., 2025; Holtgrefe et al., 2025a; Rhodes et al., 2025; Xu and Ané, 2023).

Our approach consists of two steps. First, we present a consensus framework for trees of blobs that applies to networks of any complexity. The algorithmic scheme bears similarity to the greedy construction of consensus trees (Dong et al., 2010), although there are differences between the two frameworks, as will be seen. Then, we present a consensus method for rooted and semi-directed level-1 networks. Our method, called ‘L1N consensus’, takes as input level-1 networks and returns a consensus level-1 network. We use support values for various structural network features, which we refer to as *components* (e.g., blobs, edges, and hybrid clades), measured by the proportion of input networks that share these components. Our step-by-step approach aims to maximize the support of the consensus network components using either greedy or exact algorithms, where possible. Consensus ToBs and level-1 networks, as well as any reference level-1 network, can be annotated with these support values. These provide bootstrap support when input networks come from a bootstrap analysis, or posterior probabilities when input networks come from a Bayesian posterior sample. To facilitate their application, we provide an implementation in Common Lisp as well as in the Julia package PhyloSummaries (Ané and Sharma, 2026), and illustrate their usage on both simulated and empirical datasets. Together, these approaches extend the well-established principles of tree consensus to the network setting, offering practical tools for summarizing and comparing sets of inferred trees of blobs and rooted and semi-directed level-1 networks.

## 2 Methods, Benchmarks and Data

### 2.1 Overview of the consensus methods

Our goal is to summarize a collection of input networks (Fig. 1) via a consensus tree of blobs or consensus network, together with support values for their components. The input networks are obtained using standard software programs, and thus are typically binary with all internal nodes of degree three. In rooted binary networks, *tree nodes* have one incoming edge and two outgoing edges, while *hybrid nodes* have two incoming hybrid edges and one outgoing edge that roots a hybrid clade or taxon. By way of contrast, in semi-directed networks, only the reticulation edges are directed, though edges in hybrid clades can be oriented from past to present using this knowledge. When cycles do not share any edges, the network is of *level-*1.

Phylogenetic networks are complex objects, even those of level-1. They have tree and hybrid nodes, tree and hybrid edges, blobs that induce partitions of the taxa into subsets, called *taxon blocks* in this paper, and hybrid clades. In level-1 networks, the blob-induced partitions are circular (see Fig. 1, or formal definitions in the supplement). The first step in our approach is parsing each input network and extracting a set of *components* that are used to construct a consensus. These components comprise the blobs, their hybrid clades, their circular orders of taxon blocks in the binary level-1 case, and the splits that are non-redundant with any of the blobs’ taxon blocks. Throughout, components are associated with sets of taxa; however, when no ambiguity arises, we abbreviate a set like {*A, B*} simply by *AB*. For instance, the network in Figure 1 contains two blobs: Blob_1_ corresponds to circular order *H* ↔ *G* ↔ *F* ↔ *ABCDEKLMNOP* ↔ *IJ* with hybrid clade *H*; and Blob_2_ to circular order *CDE* ↔ *AB* ↔ *P* ↔ *ON* ↔ *FGHIJKLM* with hybrid clade *CDE*. Although most standard software programs infer cycles of four or more edges, our method and software apply more generally, allowing cycles with three or more edges.

To obtain the tree of blobs of a network, the blobs of the network are contracted into single nodes. In the tree of blobs, these nodes and standard polytomies must be distinguished from each other. A blob node indicates one or more hybridization events in the underlying network, while a polytomy may result from an absence of signal or a radiation event. A network’s tree of blobs is thus labeled, in the sense that each node is labeled either as a *blob node* if it represents a blob in the network (shown as large colored dots in figures), or otherwise as a *divergence node*. In practice, blob nodes typically have degree at least four. If a network is binary, then its divergence nodes all have degree three.

In a tree, every edge induces a *split*, or bipartition, of the taxa. For example, in Figure 1, the edge rooting the hybrid clade *CDE* induces the split *CDE*|(*X* − *CDE*), where *X* denotes the set of all taxa and (*X* − *CDE*) is the set of all taxa but *C, D*, and *E*. We are interested in *non-redundant splits*, those induced by edges that are not connected (or adjacent) to blobs. For example, *IJ*|(*X* − *IJ*) is redundant since it is connected to Blob_1_. Equivalently, one block of the split, *IJ*, corresponds to a block induced by Blob_1_. The network in Figure 1 contains three non-redundant splits: *DE*|(*X* − *DE*), *KLM*|(*X* − *KLM*), and *LM*|(*X* − *LM*). In the Supplementary Material, we demonstrate that any level-1 network can be reconstructed from its non-redundant splits and blobs’ taxon blocks with associated circular orders and hybrid clades.

Once all input networks have been parsed and transformed into sets of components, the consensus network is built. The first step is to construct a consensus tree of blobs from the input networks’ trees of blobs, with their nodes labeled as blob or divergence nodes. In this consensus tree of blobs, internal nodes are labeled as blob or divergence nodes as well. Some divergence nodes may have a high degree in cases of low signal or when the same radiation is found in many input networks. Although the tree-of-blobs consensus method is applied here to level-1 networks in simulations and real data, it can be applied to collections of networks of any level, and does not require that the networks are binary. After this construction, each blob node in the consensus tree is associated with a circular order of taxon blocks and a hybrid clade to obtain a consensus level-1 network.

Overall, our approach is similar to those used for trees, where strict, majority-rule, and greedy consensuses based on the frequency of the splits in input trees are long-established (Bryant, 2003; Meacham, 1983). The strict consensus tree contains the splits found in all input trees. It is generally poorly resolved and rarely used in practice. The majority-rule consensus tree contains all splits that appear in a strict majority of the input trees. It has the advantage of being the median tree of the input trees with respect to the Robinson-Foulds (RF) distance (Barthélemy and McMorris, 1986; Robinson and Foulds, 1981). To obtain a more resolved ‘greedy’ consensus tree, splits are greedily added in order of decreasing frequency, as long as they are compatible with the splits that have already been selected (Bryant, 2003). Our approach for the consensus ToB is similar and has analogous properties, in particular regarding the median. In the following, we first describe the inference of the consensus tree of blobs and then how this is turned into a consensus level-1 network.

### 2.2 Computing the consensus of a collection of trees of blobs

In this section, we make a few assumptions on the class of input networks for our methods. They can be of any level and non-binary. These networks are transformed into trees of blobs (ToBs), and each ToB is converted into a set of *partition* components: blob partitions (or blobs for short) and non-redundant splits. A split defines a bipartition of the taxa, and a blob defines a partition of the taxa. For instance, Blob_1_ in Figure 1 divides the taxa into five subsets: {*F*}, {*G*}, {*H*}, {*I, J*}, and *X* − {*F, G, H, I, J*}. For simplicity, such a blob is denoted by *F*|*G*|*H*|*IJ* |(*X* − *FGHIJ*). Note that taxon blocks in blobs are not ordered, but checking for equivalence can be achieved in time *O*(*n*), where *n* is the number of taxa (see the Supplementary Material).

The condition for the tree-compatibility of two splits is well known (Semple and Steel, 2003). Two splits *A*|*A*′ and *B*|*B*′ are *tree-compatible* if *A* or *A*′ is included in *B* or *B*′. For example (Fig. 1), the splits *LM*|(*X* − *LM*) and *KLM*|(*X* − *KLM*) are tree-compatible, since *LM* is included in *KLM*. Note that (*X* − *KLM*) is also included in (*X* − *LM*), and that this condition is symmetric.

The notion of *blob-compatibility* (*compatibility* for short) generalizes this idea. A collection of non-redundant splits and blob partitions are compatible if they can coexist in a single ToB. Clearly, tree-compatible splits are compatible in this sense. For two blobs, if we let *A* = *A*_1_|*A*_2_|…|*A*_*p*_ and *B* = *B*_1_|*B*_2_|…|*B*_*q*_ be their blob partitions, then they are compatible if all *A*_*i*_’s but one are contained in one of the *B*_*j*_’s. For example, Blob_1_ and Blob_2_ in Figure 1 are compatible since *F*, *G, H*, and *IJ* are included in *FGHIJKLM*. This condition is again symmetric since *AB, CDE, NO*, and *P* are included in (*X* − *FGHIJ*). Note that in this example, the inclusions are strict (*e*.*g*., *FGHIJ* ⊊ *FGHIJKLM*), but equality can and does occur when the two blobs in a network are connected by a single edge.

The compatibility of a split and a blob partition follows the same line. A split *S* = *A*|*A*′ is compatible with a blob partition *B* = *B*_1_|*B*_2_|…|*B*_*q*_ if *A* or *A*′ is included in one of the *B*_*i*_’s, and it is easily seen that this definition is symmetric. For example, *AB*|(*X* − *AB*) is compatible with Blob_1_ since *AB* is included in (*X* − *FGHIJ*) and (*X* − *AB*) contains *F*, *G, H* and *IJ*. However, for a split to be non-redundant with a blob partition, this inclusion must be strict. For example, *AB*|(*X* − *AB*) is compatible with Blob_2_, but it is redundant with this blob, since *AB* is equal to one of its taxon blocks.

A naïve implementation makes it possible to compute the compatibility of two partition components (blobs or splits) in *O*(*npq*) time, where *n* is the number of taxa and *p* and *q* are the number of taxon blocks in the two components (2 for splits, and ≥ 3 for blobs). In the Supplementary Material, a faster algorithm in *O*(*n* + *pq*) time is described. Also there, we show that if a collection of blobs and non-redundant splits is pairwise compatible, then this collection is globally compatible and defines a unique tree of blobs. This fundamental property makes the construction of the majority-rule consensus tree of blobs straightforward:

1. Collect all partition components (from blob nodes and non-redundant splits) from the input ToBs.
2. Calculate the frequency of these components among the input networks, and collect those present in a strict majority.
3. This collection uniquely defines the *majority-rule consensus tree of blobs*, with each node labeled as a blob or divergence node.

Importantly, in step (2), the frequency of splits only counts the networks in which they are non-redundant. When a split is induced by a blob partition, for example, the bipartition is *AB*|(*X* −*AB*) induced by Blob_2_ in Figure 1, it does not contribute to the frequency computation. This prevents the overemphasis of components that occur both as non-redundant splits and as redundant splits induced by blobs, which otherwise might result in the exclusion of some blobs in the final consensus. Note that any two components from step (2) are compatible because they belong to at least one input ToB.

Full details for step (3) are given in the Supplementary Material. There we demonstrate that the majority-rule consensus ToB is included in the standard majority-rule consensus tree in terms of splits, but it can differ from the latter despite the similarity of the construction algorithms (see also Fig. 3). This is because, in ToBs, blobs are of the utmost importance since they correspond to hybridization events in the underlying networks and are thus preserved and not resolved, unlike polytomies in consensus trees. We also prove that the majority-rule consensus ToB is the median (or centroid) of the input ToBs. To demonstrate this, we use a natural distance between two ToBs, which is the size of the symmetric difference between their sets of partition components. The majority-rule consensus tree of blobs minimizes the sum of the aforementioned distances to the input ToBs.

The number of partition components collected in step (1) is *O*(*kn*), where *k* is the number of input ToBs and *n* is the number of taxa. Since checking the equality of two partition components takes *O*(*n*) time, and we must compare *O*(*k*^2^*n*^2^) pairs of such components in step (2), a naïve implementation of this algorithm has *O*(*k*^2^*n*^3^) time complexity. However, this is a worst-case scenario. The algorithm can be accelerated considerably by using hash-coding techniques and possibly some of the techniques used for the standard majority-rule consensus tree, achieving optimal time complexity of *O*(*kn*). In practice, our simple implementations are very fast. In our simulation experiments involving 21 taxa and 100 input networks, computing the majority-rule consensus ToB takes less than 0.01 seconds in terms of both real and CPU time on a standard laptop.

While the construction of the majority-rule consensus tree of blobs is straightforward and has the important median property, the resulting ToB can be under- or even unresolved. Indeed, in satisfying the majority criterion, it can contain divergence nodes of degree greater than three that represent uncertainty, rather than a radiation shared by more than 50% of the input networks. Importantly, these divergence nodes might be further resolved by adding blobs or splits present in fewer than 50% of the input ToBs, yielding more summary information about the input. To address these concerns, we implemented a greedy consensus tree-of-blobs algorithm, which we now outline. Full details are provided in the Supplementary Material.

1. Compute the frequencies of all blob and non-redundant split taxon partitions (components) from the set of input ToBs, and construct the majority-rule consensus tree of blobs. If this ToB has no divergence nodes of degree greater than 3 and if the input ToBs have no degree-3 blob nodes, then return it. If not, order the remaining components by decreasing frequency, prioritizing blobs over non-redundant splits when breaking ties.
2. Select the most frequent of these remaining components, calling it *C*. If *C* is compatible with all components in the current ToB, then *C* is added to these components. Move to the next component in the ranked list, call it *C*, and repeat this process. These iterations stop when all divergence nodes in the tree of blobs in construction have degree 3, and the list has no more degree-3 blob partitions, or the list of available components is empty. The final iterate is returned as the greedy (or extended) consensus tree of blobs.

Note that step (2) can add a blob partition that causes a previously-added split to become redundant (see example in Figure 3).

Though there is no formal guarantee, the greedy consensus ToBs did not contain divergence nodes of degree greater than 3 for all the datasets we analyzed. In some cases, the greedy consensus contained more blob nodes than the majority-rule version. However, often there were only additional splits. Furthermore, due to ties, the greedy consensus ToB is not guaranteed to be unique. In terms of computing time, it is difficult to perform a time complexity analysis of such a greedy heuristic. For the datasets we analyzed, our simple implementations only required a negligible amount of extra time compared to the majority-rule consensus. Most of the time is actually spent parsing the input networks in extended Newick format to obtain the blob partitions and non-redundant splits. Ordering these components based on their frequency and checking their compatibility is then very quick.

As a detailed example, consider the five rooted networks shown in Figure 2, where taxon *A* is an outgroup, and their corresponding unrooted trees of blobs. These 9-taxon networks put forth strong evidence that *D*_1_ and *D*_2_ are sister taxa and that hybridization occurred among the group {*M*_1_, *M*_2_, *M*_3_}, a hunch confirmed more rigorously by our consensus trees of blobs and networks (Figs. 3, 4). Also shown in Figure 2 is a table with the frequency of shared components used in our algorithms. Specifically, all blob partitions are listed, but only non-redundant split components of frequency greater than 1/5. Figure 3 shows the majority-rule and greedy consensus tree of blobs for this input, and the standard majority-rule consensus tree.

**Figure 2:**
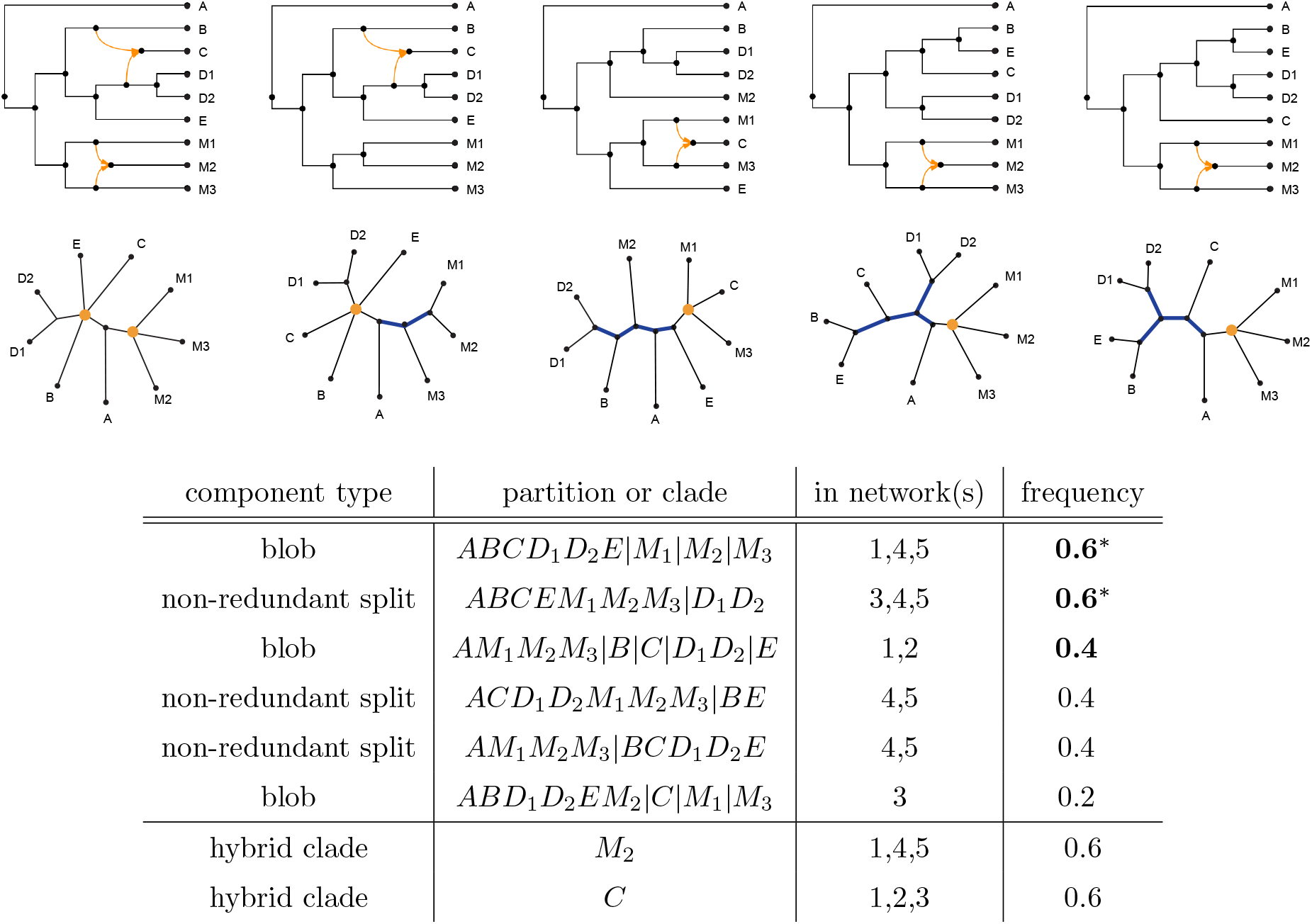
A sample of five rooted level-1 networks on 9 taxa (top), and their unrooted trees of blobs (middle). Blob nodes are shown in orange, and non-redundant splits in blue. For simplicity, from left to right, the networks, respectively, trees of blobs, are numbered *k* = 1, …, 5. The table (bottom) shows the most frequent components (partition components then hybrid clades). The most frequent partition components are listed by decreasing frequency, with blob partitions ranked higher than non-redundant split partitions when ties occur. Partitions in the majority-rule consensus tree of blobs have frequencies displayed in bold with an asterisk, and those in the greedy consensus ToB in bold. In building the greedy consensus, the blob partition with frequency 0.4 is chosen over the tied non-redundant splits. This greedy consensus ToB is identical to the ToB of network 1.

**Figure 3:**
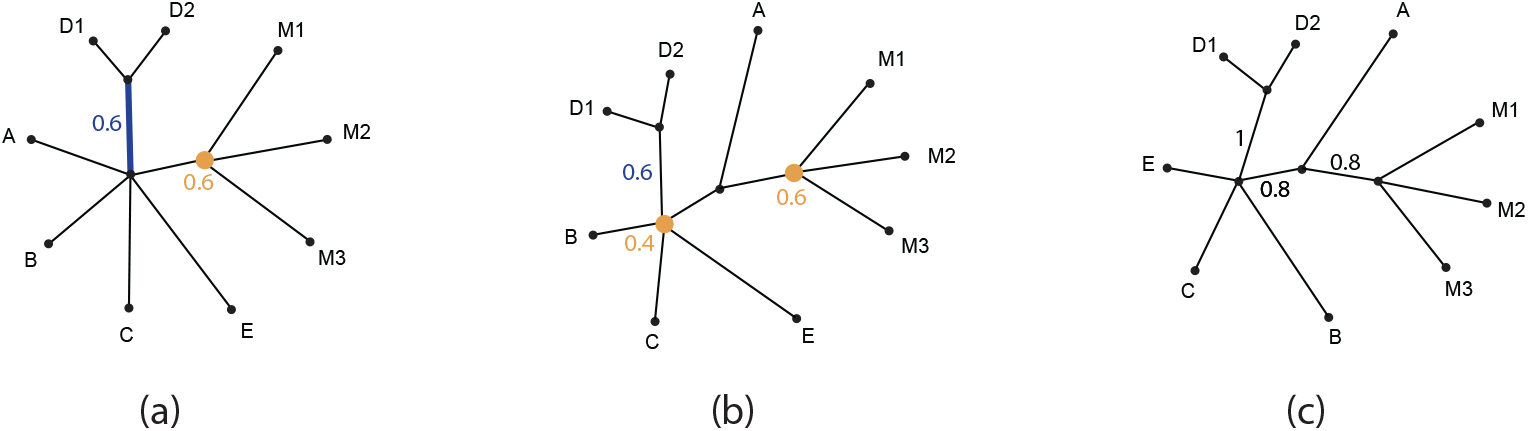
(a) Majority-rule ToB for the networks in Figure 2. Both the blob node (orange dot) and the non-redundant split (blue edge) occur in 3/5 of the networks, and have support 0.6. Although the split *D*_1_*D*_2_|(*X* − *D*_1_*D*_2_) occurs in all five input networks, twice it appears as a redundant split, which does not contribute to its support value. (b) Greedy consensus ToB with an additional orange blob node labeled with its support 2*/*5 = 0.4. The edge leading to {*D*_1_, *D*_2_} remains labeled with its non-redundant split support 0.6, as it occurs in the majority-rule tree of blobs, but this edge is adjacent to an orange blob node and becomes redundant in this greedy consensus ToB. (c) For contrast, the traditional majority-rule consensus tree of the 5 networks’ ToBs is shown with support values in black. This differs from the majority-rule ToB by having an extra edge, whose split *AM*_1_*M*_2_*M*_3_ | (*X* − *AM*_1_*M*_2_*M*_3_) appears in 4/5 of the ToBs (redundant in 2/5), illustrating that only support for non-redundant splits should be included when considering also blob nodes. In this small example, the greedy consensus ToB (b), the majority-rule consensus tree (c), and the first input ToB (Fig. 2) have the same tree topology. However, this is not a general rule. See the Supplementary Material for another example with different outcomes.

**Figure 4:**
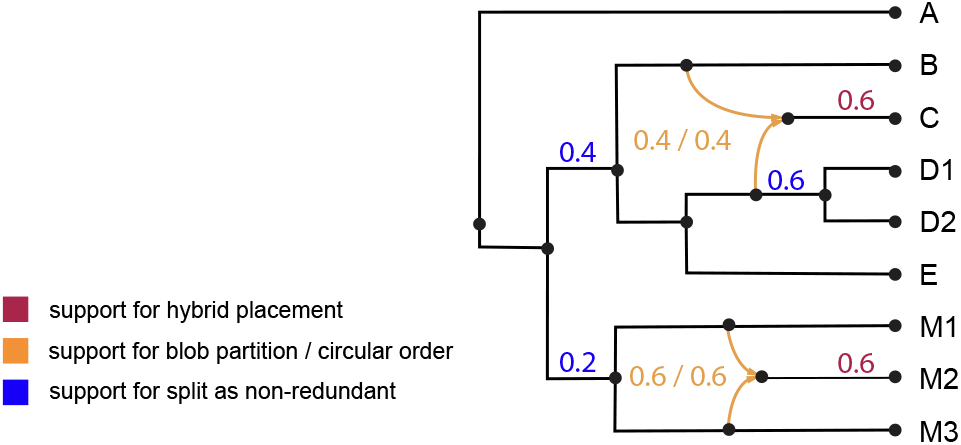
Greedy consensus level-1 network for the five level-1 networks from Figure 2. This consensus network is rooted using the outgroup *A*, and is identical to input network 1. Note that all splits are redundant in this consensus. Support values, computed as frequencies in the input networks, for blob partitions, circular orders, non-redundant splits, and hybrid clades are shown. In this example, the support for a circular order equals that of the blob partition (orange) because each cycle appears with a unique circular order. Note that the edge above the cherry {*D*_1_, *D*_2_}, though redundant in this consensus network, has a non-redundant support value of 0.6 because it occurs in 3/5 of the networks as non-redundant. This contrasts with its traditional split support value of 1, computed from its occurrence in all five input networks.

In this small toy example, the traditional majority-rule consensus tree, the greedy consensus ToB, and one of the input ToBs share the same topology. However, this is generally not the case. Moreover, there is no guarantee that a greedy consensus ToB is unique since, similar to the traditional greedy consensus tree algorithm, these constructions typically work from a list of conflicting (non-compatible) components and thus depend on the order in which tied components are added to the tree. The Supplementary Material provides a modified version of this example that illustrates the possibility of several greedy consensus ToBs that do not match the traditional majority-rule consensus tree or any of the input ToBs.

### 2.3 Expanding a consensus tree of blobs into a consensus level-1 network

In this section, we assume that all input networks are binary and of level at most 1 (trees or networks with non-overlapping cycles). The first step is to compute their greedy consensus ToB as described above. The second step is to assign a circular order together with a hybrid clade to each blob node of this consensus ToB. To do so, when parsing the input networks we track each cycle and its hybrid clade in the networks (see above examples and Fig. 1), and thus for each blob node in the consensus ToB we have a collection of associated circular orders belonging to (some of) the input networks. The support of this blob node is then the (possibly weighted) proportion of these, as a fraction of the input size. To select the particular circular order resolving that given blob node, we simply choose the most frequent one. For making this choice, we must determine when two circular orders are equal. For example, the circular order *AB* ↔ *C* ↔ *D* ↔ (*X* − *ABCD*) is equivalent to the circular orders *D* ↔ (*X* −*ABCD*) ↔ *AB* ↔ *C* (rotation) and *AB* ↔ (*X* −*ABCD*) ↔ *D* ↔ *C* (reversal). All of these representations resolve the blob node in the same manner. We achieve this computation by rotating and reversing the circular orders and comparing the corresponding ordered partitions. This computation takes *O*(*np*) time, where *p* is the number of taxon blocks in the blob.

Assigning the hybrid clade to a given blob node in the consensus ToB follows a similar approach. If no outgroup is available and the consensus ToB is unrooted, then all taxon blocks associated to a given blob node are potential hybrid clades. When the consensus ToB is rooted using an outgroup, the taxon block containing the outgroup is excluded as a potential hybrid clade. In both scenarios, the goal is to select a hybrid clade with the highest frequency. However, since a hybrid clade may arise from distinct blobs (*e*.*g*., from blobs nodes of differing degrees), its frequency may be greater than that of any single blob partition in which it is hybrid. Another complication is that hybrid clades must be mutually compatible in the sense that there is a level-1 network that induces all of them. Concretely, let *C* and *H* be two potential hybrid clades. If their intersection is empty or equal to one of *C* or *H*, then *C* and *H* are compatible. For example, in Figure 1, after removing the outgroup *P*, the block *X* − *ABCDENO* in Blob_2_ is compatible with *IJ* in Blob_1_, but not with *X* − *FHGIJ*. Two useful observations follow. First, when one or both hybrid clades are reduced to a single taxon, they are necessarily compatible. Second, if two hybrid clades both have frequency greater than 50%, they must also be compatible, since they co-occur in at least one input network. In practice, for the datasets we have analyzed, incompatibilities were rare.

Determining hybrid clades is thus formulated as an optimization problem. When input networks are rooted using an outgroup, the consensus ToB inherits this rooting and all potential hybrid clades are mutually compatible. In this case, we simply select the most frequent candidate hybrid clade(s).

When no root or outgroup is available, we instead consider all possible rootings induced by each taxon *x*. For fixed *x*, we discard all potential hybrid clades that contain *x*. The remaining potential hybrid clades are then mutually compatible, and we again select, independently for each blob node, the most frequent potential hybrid clade. This produces a candidate solution, which we score by the sum of the selected hybrid clades’ frequencies. Repeating this procedure over all taxa and retaining the best-scoring solution yields an exact and efficient algorithm. This approach relies on structural assumptions (see Supplementary Material), namely that blob nodes have degree at least three and that they arise from level-1 networks. Under these assumptions, a set of potential hybrid clades (one per blob node) is compatible if and only if there exists a taxon *x* that is excluded from all of them, which justifies our rooting-based exact algorithm.

Once the circular orders and hybrid clades attached to the consensus ToB have been selected, we reconstruct and output the corresponding level-1 network. The frequency (support) is provided for each of the components (blob partitions, circular orders, hybrid clades, and non-redundant splits) of this network. As an example, an annotated greedy consensus network from our earlier example is shown in Figure 4. Computing such a level-1 consensus network is fast. In our simulation experiments involving 21 taxa and 100 input networks, the computation took approximately 0.05 seconds of CPU time and 0.1 seconds of real time on a standard laptop.

### 2.4 Implementation of tree of blobs and level-1 consensus in PhyloSummaries

The consensus methods, first developed in Lisp, were implemented in the Julia package PhyloSummaries (Ané and Sharma, 2026). It depends on the core package `PhyloNetworks` (Solís-Lemus et al., 2017) for network utilities, such as parsing input networks and saving output networks using the extended Newick format (Cardona et al., 2008). The key functions, `consensus_treeofblobs` and `consensus_level1network`, take a list of networks as input. They output a greedy consensus by default, with data tables to list the support values of components.

For the level-1 consensus, users can specify one taxon as the outgroup, which affects the choice of hybrid clades in the consensus. Users can specify a value *p*, to exclude components appearing in a proportion *p* or fewer input networks, with *p* = 0.5 corresponding to the majority rule ToB consensus. A list of network weights can also be provided. By default, all input networks are weighted equally. Another optional argument is the minimum degree *k* ≥ 3 for blobs to be considered. Specifically, if an input network has a blob of degree *d < k*, this blob is contracted into a tree node in the input network (and suppressed if *d* = 2) before calculating the consensus. By default *k* = 4 since blobs of degree 3 are often unreliable, and it may be preferable to ignore them when building the consensus network. The package has extensive documentation, including a tutorial with example data, code, output, and interpretation (see Fig. 5).

**Figure 5:**
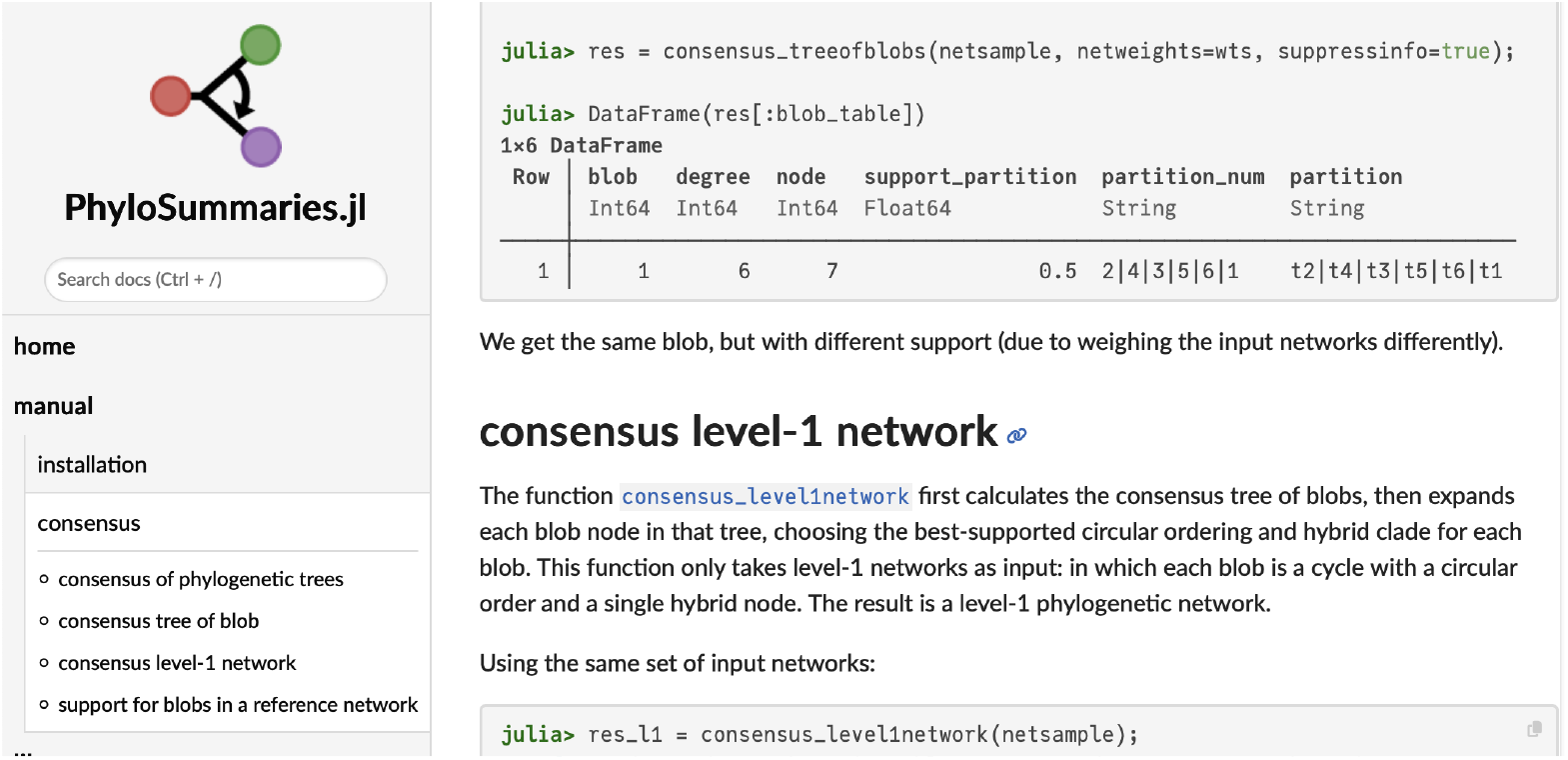
Example section from the online documentation of the `Julia` package `PhyloSummaries`.

#### Support values

Our implementation further returns tables listing the support for various components in the consensus ToB and level-1 network, with support measured by the proportion of input networks that have the component. If network weights are provided, support is measured as the sum of the weights of the networks that have the component of interest (normalized by the total sum of weights). For each blob (node or cycle) in the consensus, we provide the support for its partition into taxon blocks, the support for each circular ordering of its blocks if the input networks are of level 1, and for each one of its taxon blocks, the support for this block as a hybrid clade (in this or any other blob). These support values provide extra information that complements the consensus level-1 network, by listing alternative circular orders of blocks and alternative hybrid clades that are not in the consensus level-1 network but are still supported by some input networks.

For each split in the consensus, we provide the support for this split as non-redundant with respect to blobs in input networks. Note that some splits that are redundant with a consensus blob partition may receive positive support as non-redundant in input networks and vice versa: a non-redundant split in the consensus network may receive some support as redundant with some blobs in input networks. Furthermore, the redundant support of a redundant split can be higher than the support of the corresponding blob partition, but not lower. The sum of the redundant and non-redundant supports is equal to the standard split support, as used with phylogenetic trees, for example.

#### Support for blob properties in a reference network

As a further enhancement, we implemented the function `blobpartitions_support` which takes the same input as the consensus functions, and additionally a reference network *N*_0_. For each blob in *N*_0_, it calculates the support for the same blob properties listed above. Specifically, support values are reported for splits in *N*_0_ as non-redundant, and for each blob *B* of *N*_0_, support for *B*’s partition into taxon blocks, for *B*’s circular ordering of taxon blocks when *N*_0_ and input networks are of level-1, and for each of *B*’s blocks as candidate hybrid clade. Additional supports available in PhyloNetworks are described in the Supplementary Material.

### 2.5 Simulation settings

We performed simulations to compare these novel consensus methods with other approaches. Complete details and software options are provided in the Supplementary Material, while the main steps of the simulation pipeline are summarized below.

We generated 20 rooted level-1 phylogenetic networks with 20 taxa using `SiPhyNetwork` (Justison et al., 2023). These networks were required to have either at least three cycles of size at least 4 or one cycle of size at least 6. Some of the retained networks contain cycles of lengths 2 and 3. While this is realistic, such short cycles cannot be recovered using current inference methods such as `Squirrel` (Holtgrefe et al., 2025b), which was used in this simulation study. Similarly, hybrid clades resulting from cycles of length 4 cannot be identified using some quartet-based methods like `Squirrel`. On average, there were 3.35 cycles of size greater than 3 per network (67 in total for the 20 networks), of which ~50% (34 in total) were of size 4. Two such networks are shown in Figure 6; the others are provided in the Supplementary Material. To ensure that ILS was moderately high, but not too high, we resampled all branch lengths shorter than 0.4 coalescent units (cu) to be at least 0.4 cu. Additionally, to ensure a sufficiently strong hybridization signal, hybridization parameters smaller than 0.3 were set to 0.3, with the complementary hybridization parameter set to 0.7.

**Figure 6:**
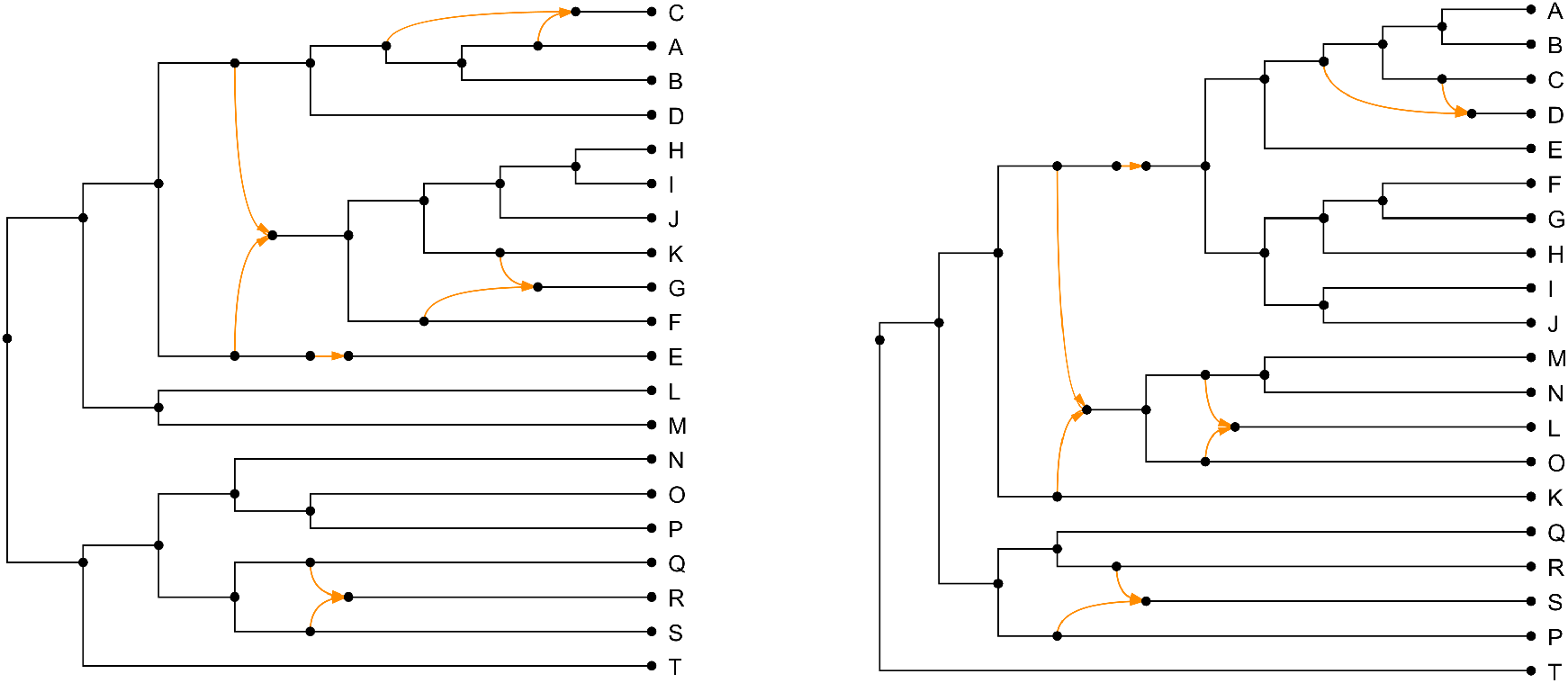
Two of the 20 level-1 phylogenetic networks with 20 taxa used in the simulation study. Both networks have a 2-cycle, depicted with single orange arrows.

For each network, we simulated 400 gene trees under the Network Multispecies Coalescent model using the `PhyloCoalSimulations` package (Fogg et al., 2023) with default parameters. We rescaled gene branch lengths (in number of substitutions per site) of the gene trees so that they matched the average length observed in real gene tree data used in a similar phylogenetic network context (Kong et al., 2024). To obtain realistic gene trees, we introduced a deviation from the strict molecular clock (ultrametricity) and globally rescaled the gene trees to reflect the fact that some genes evolve faster than others. Once again, parameters of these gene tree perturbations were selected using real data. Finally, an outgroup taxon was grafted onto the root of each gene tree, resulting in 21-taxon trees (and model networks).

We next simulated DNA sequences of varying lengths along gene trees using `AliSim` (Ly-Trong et al., 2022). The minimum sequence length was ~300 bp, with a mean of ~600 bp. We used a GTR+G4 model with realistic parameter values, and inserted gaps using the mimic option of `AliSim` and real data (Vanderpool et al., 2020) as a template.

Networks were inferred using `Squirrel` (Holtgrefe et al., 2025b), with its *λ* parameter (which controls sensitivity to tree-like versus hybrid signals in the data) set to 0.15. We selected this value so that the inferred networks with simulated data had an average number of cycles of size at least 4, similar to the real one (~3.1 versus 3.35 for the 20 model networks). Unlike other methods, `Squirrel` does not use gene trees, but instead uses the full multiple sequence alignment (MSA) obtained by concatenating all gene MSAs. Analyses were performed under different settings. In the ‘Third’ condition (using only a third of the sites per gene), the signal was substantially lower than in the ‘Full’ condition (using all the sites per gene). We also analyzed the data with and without knowledge of the outgroup taxon. When outgroup knowledge is given, the problem is easier since, by design, `Squirrel` will not infer a hybrid clade containing the outgroup. The probability of error is thus lower. Furthermore, in this ‘Rooted’ condition, `Squirrel` produces rooted networks (using the outgroup) that can be analyzed using the HMS (Huber et al., 2023) and HWC (Nielsen et al., 2023) consensus methods. In the ‘Unrooted’ condition, these methods are not applicable, and our greedy level-1 network (L1N) consensus method is the only option.

There were four settings in total: two for the outgroup signal (one in which outgroup information was provided to Squirrel and one in which it was not), and two for the length of the data (one using the full MSA and one using only one third of each gene). The one with the highest signal was ‘Rooted & Full’, while ‘Unrooted & Third’ was expected to have the lowest signal, resulting in greater difficulty in recovering the true network and greater uncertainty measured by support values.

To obtain a collection of networks for analysis by consensus methods, we employed a block bootstrap approach (Felsenstein, 1985), based on the gene-based partition of the concatenated MSA. In this method, we resample genes (or blocks) with replacement, while retaining the original site configuration within each selected gene. This approach mimics the random sampling of genes from a large genome. We applied the same procedure for both the ‘Full’ and ‘Third’ settings; in the latter, only the first third of the sites of each selected gene were considered. For each of the four settings, we conducted 100 block bootstrap replicates, yielding 100 level-1 networks of 21 taxa, either rooted under the ‘Rooted’ condition or left unrooted. To account for the variability introduced by the resampling of genes, we repeated the block bootstrap approach 10 times. In total, we thus obtained 20 (model networks) *×* 4 (settings) *×* 10 (replicates) sets of 100 input bootstrap networks, which were given to the various consensus methods for comparison purposes.

We measured the performance of these consensus methods under two different correlated angles: how well they represent the input bootstrap networks and how accurately they recover the true network used to simulate the data. As explained in previous sections, we considered several types of network components: the blob partitions (or blobs for short); the circular orders of blob blocks (or cycles); the hybrid clades (or hybrids); and the non-redundant splits. We showed that a level-1 network is uniquely defined by its components (see Supplementary Material for a formal proof). Two networks are identical if and only if they have the same set of components. When they differ, we can measure the extent of the difference between their component sets. To illustrate the behavior, strengths, and weaknesses of the various consensus methods, we examined each type of component separately. To compare the sets of components of two networks, we used false positive (FP) and false negative (FN) errors, as well as the sum of these errors (i.e., the symmetric difference of the two sets of components), and various normalizations of these measurements.

First, we describe how we measure the dissimilarity between the input bootstrap networks and a given consensus network. The primary goal of consensus methods is to minimize this dissimilarity. Let *N*_*T*_ be one of the 20 true model networks generated as described above, *N*_*I*_ be an input bootstrap network inferred from the data generated under *N*_*T*_, and *N*_*C*_ be a consensus network inferred using method *C* from the bootstrap networks associated with *N*_*T*_. We describe the evaluation measures for a single model network *N*_*T*_; for each setting, the measures are subsequently averaged over the 20 model networks and 10 replicates. Consider a specific component type; for example, consider the blob partitions (or blobs for short). 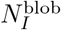 and 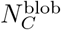 are the set of blobs induced by *N*_*I*_ and *N*_*C*_, respectively. The FP error of 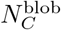 with respect to 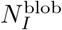 is equal to the number of blobs belonging to the consensus *N*_*C*_ but not the input *N*_*I*_. Similarly, the FN error of 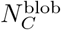 with respect to 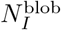 is equal to the number of blobs belonging to the input *N*_*I*_ but not the consensus *N*_*C*_. The total error of 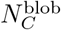 with respect to 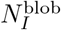 is the sum of the FP and FN errors, which is equal to the size of the symmetric difference between 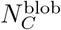 and 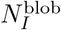. We call this sum the symmetric distance between 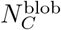 And 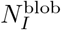 it is a metric distance since it satisfies the three properties of a metric (reflexivity, symmetry, and the triangle inequality) and lends itself to the standard Robinson-Foulds (RF) distance in the case of splits. To normalize the symmetric distance and make it easier to interpret, we divide the symmetric difference between the blob sets by twice the number of blobs in the corresponding model network, 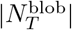. To account for the fact that the various consensus methods infer component sets with substantially different sizes, we also used the Jaccard distance between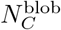 and 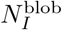. Furthermore, we averaged these measurements across the 100 input bootstrap networks used for each model network, replicate, and setting. More formally:

False positives:

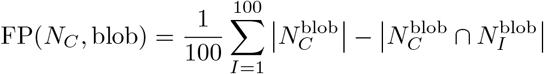

False negatives:

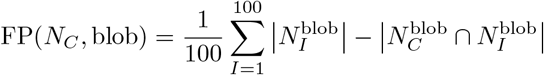

Symmetric distance (normalized):

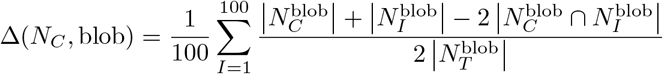

Jaccard distance:

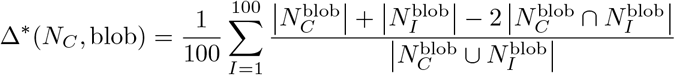

The latter formula can be problematic when both the consensus and the inferred network do not contain any blob. In this case, the Jaccard distance is consistently set to zero.

The smaller the symmetric (Δ) and Jaccard (Δ*) distances, the better *N*_*C*_ represents the input networks with respect to blobs. As explained above and in the Supplementary Material, the majority rule is optimal with respect to the symmetric distance Δ (e.g., the RF distance in the case of splits). Here, this means that the set of blobs belonging to the majority of input networks minimizes the symmetric distance Δ. However, this property does not hold for the Jaccard distance Δ*.

For each setting (e.g., ‘Full & Rooted’), we averaged these metrics across the 20 model networks and 10 replicates to obtain Δ(*C*, blob) and Δ*(*C*, blob). These represent the ability of the consensus method *C* to capture the input networks in terms of blobs in the given setting. Similarly, we defined Δ(*C, X*) and Δ*(*C, X*), where *X* ∈ {blob, circular order, hybrid, split}, to measure the performance of *C* with respect to the four network component types. To ease interpretation of the results, all splits (both redundant and non-redundant) were considered. For blobs and circular orders, we considered the blobs of size at least 4. Furthermore, only hybrid clades occurring in blobs of size greater than 4 were considered. This was done to avoid blurring the comparisons, and because hybrid clades in blobs of size 4 are not identifiable by Squirrel. Finally, to evaluate the global dissimilarity in the given setting between the consensus network defined by method *C* and the input networks, we averaged the results across the four component types:

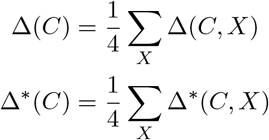

It should be noted that by following these definitions and calculating the average of the distances between sets of components, both the symmetric difference and the Jaccard distance yield proper metric distances between binary semi-directed level-1 networks without 2-cycles, provided that blobs of size at least 3 and their hybrid clades are considered. These results hold because (1) each network is uniquely defined by its components, and (2) each of the four component-based distances is a proper metric on the component sets. To the best of our knowledge, these easily tractable distances have not been previously described in the context of level-1 networks (see Supplementary Material for additional details).

We used similar criteria to evaluate the performance of the various consensus methods in recovering the model networks used to simulate the data. Although this is not the primary objective of consensus methods, we would expect to see a correlation between the accuracy results obtained using these criteria and those obtained using the above criteria to measure the closeness of the consensus to the inputs. Such results have been observed in several studies on consensus trees in relation to RF distance (Berling et al., 2025; Berry and Gascuel, 1996; Holder et al., 2008; Takazawa et al., 2026). Furthermore, unless there is a strong bias in the inference method, we would expect the accuracy of the consensus to be higher than the average accuracy of the inputs. This is similar to the reason why averaging various numerical estimates generally improves the estimation by reducing the variance component of the error.

To measure the accuracy of recovering the true model network *N*_*T*_, we used the normalized symmetric difference and the Jaccard index, as well as precision, recall, and the F1 score. The symmetric difference is defined as shown above and normalized in a similar way. The Jaccard index is equal to 1 minus the Jaccard distance. The precision of a given consensus network, *N*_*C*_, for blobs, is equal to the proportion of blobs in *N*_*C*_ that belong to the model network *N*_*T*_. A blob precision of 1 indicates that all blobs in the consensus network are correct. The recall of *N*_*C*_ is equal to the proportion of blobs in *N*_*T*_ that are retrieved in *N*_*C*_. A blob recall value of 1 indicates that all true blobs are recovered in the consensus network. The F1 score is a standard statistical combination of precision and recall; it is equal to the harmonic mean of these two accuracy measures. The symmetric difference must be minimized (the lower the better), while all other criteria must be maximized and are normalized between 0 and 1. As before, let 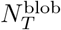 and 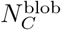 be the sets of blobs in the model network and the consensus network inferred by method *C*, respectively. These criteria are defined as follows:

Symmetric distance (normalized):

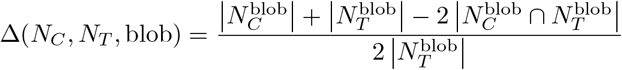

Jaccard index:

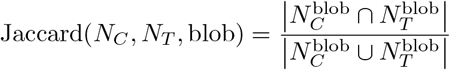

Precision:

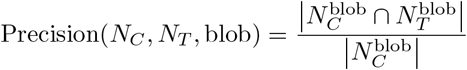

Recall

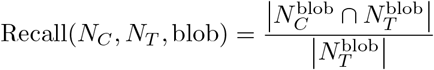

F1 score:

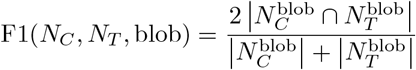

The formula for precision is problematic when 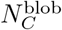 is empty. In this case, we consistently set the precision value to zero (in our simulated data, 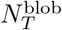 was never empty and the same holds true for the other network components).

As before, for each setting, these measures were averaged over the 20 model networks and 10 replicates, and generalized to the four network components. By computing the mean over the four components, we obtained the following global criteria measuring the accuracy of *C* in recovering the true model network in the given setting: Δ(*C*), Jaccard(*C*), Precision(*C*), Recall(*C*), and F1(*C*).

The consensus methods compared are:

- HMS (Huber et al., 2023) consensus, which applies only to rooted networks. The method is based on an encoding of so-called 1-nested networks (rooted networks that are slightly more general than level-1 networks and coincide with them in the binary case) as collections of ordered pairs of subsets of the taxon set. Intuitively, each such pair corresponds to a node *v* and records (1) the taxa that can only be reached from the root via *v*, and (2) the taxa that descend from *v* but can also be reached from the root along a path that avoids *v*. For a collection of input networks encoded by these *set pairs*, the HMS consensus constructs a majority-rule-type consensus network by selecting the most frequently occurring set pairs. See the Supplementary Material for further details on the HMS method and our Perl implementation.
- HWC (hardwired cluster) consensus in AdmixtureBayes (Nielsen et al., 2023), which applies only to rooted networks. This method is based on the *hardwired clusters* (Huson et al., 2010) of the input networks. For each node *v* in a network, its hardwired cluster is the set of taxa that descend from *v*. Given a threshold *α* (we used *α* = 0.5), the HWC consensus network is obtained by retaining only those clusters that appear in strictly more than an *α* proportion of the input networks, and constructing the Hasse diagram of the resulting set of clusters (see the Supplementary Material for further details). Note that the resulting consensus network is not necessarily level-1, even when all input networks are level-1. Also, the current implementation of HWC in AdmixtureBayes outputs only a graphical representation (plot in PDF format) of the consensus network. In addition, the input format for networks is not based on the standard extended Newick format, requiring potential users to reformat their input networks prior to analysis. To address these limitations, we used our Perl implementation of the HWC consensus (see Supplementary Material for details).
- Our L1N greedy consensus for level-1 networks, implemented in PhyloSummaries using the `consensus_level1network` function.
- L1N-majority: for each component (blobs, circular orders, hybrid clades, and splits) in the L1N consensus, we recorded whether is was supported by a strict majority of the input networks, to classify it as either ‘recovered’ or not, and the sets of recovered components were used in the above formulas in place of 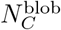. With regard to splits, we used the standard support, as we considered both redundant and non-redundant splits. We used L1N-majority to make a fair comparison with HMS, which also uses a form of majority rule, and with HWC, which we ran under a majority rule setting (using *α* = 0.5). Furthermore, L1N-majority provided us with a measurement of the usefulness and informativity of the supports. Note, however, that L1N-majority does not always define a level-1 network. For example, a given blob partition and its hybrid clade may have support higher than 50%, while all the corresponding circular orders may have support lower than 50%. This illustrates the difficulty of defining a majority-rule level-1 network and shows the advantage of using support values combined with a well-resolved network, as provided by L1N consensus.

To obtain further points of comparison, we applied the above criteria to the bootstrap networks provided as input to the consensus method. For instance, we calculated the average symmetric and Jaccard distances (refereed as ‘TRUE’ bars in figures) between the true model network used to generate the data and the input bootstrap networks. Typically, we would expect the average distance between the true network and the input networks to be greater than the average distance between the input networks and any consensus network, since consensus methods tend to minimize this distance. Similarly, regarding accuracy measurements (e.g., precision and recall), we would expect any consensus network to be more accurate, on average, than the input networks (average accuracy measurements of the input networks are referred as ‘Input Nets’ in figures).

### 2.6 Leopardus data

Full-genome data for *Leopardus* has been studied to understand contentious phylogenetic relationships between these Neotropical felids (Lescroart et al., 2023), which occupy a broad range of habitats, spanning from Texas to Patagonia, and from high-altitude Andean to lowland rainforest, with a new species recently discovered in Bolivia (Lescroart et al., 2026). Among other conclusions, Lescroart et al. (2023) describe extensive local tree discordance between *g* = 16, 338 maximum likelihood local trees constructed from filtered 100-kb sliding windows along aligned genomes from 16 taxa, with Puma (*Puma concolor*) as outgroup. To explore these data, we carried out a bootstrap analysis using 100 replicates. For each replicate, we (1) sampled *g* trees with replacement from the original dataset; (2) used TINNiK (Allman et al., 2024b) with T3-test level *α* = 5 · 10^−29^ and *β* = 0.95 to construct a tree of blobs; and (3) used NANUQ^+^ (Allman et al., 2025) to resolve nodes of degree larger than 3 into cycles. Test levels were chosen to be consistent with the analysis in Allman et al. (2024b) and Allman et al. (2025). Based on the Network Multispecies Coalescent, which models ILS, both TINNiK and NANUQ^+^ do not consider information about the outgroup or rooting position, producing unrooted trees of blobs and fully resolved, semi-directed level-1 networks, respectively. When NANUQ^+^ returned *k* tied multiple optima, they were weighted uniformly, each by 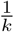. This produced 755 bootstrap networks, all of which shared a 5-cycle with the same circular order. However, each taxon block in this 5-cycle was equally likely to be the hybrid clade. Accordingly, the number of NANUQ^+^ solutions, *k*, was a multiple of 5 for all replicates. A consensus network was constructed from these 755 input networks and their weights using L1N, ignoring knowledge of the outgroup, and support for consensus network components was computed. For comparison with other methods, the HMS and HWC consensus networks were computed using the subset of 476 input networks that permitted rooting along the external edge leading to the Puma outgroup, including weights. An L1N consensus was also constructed using this subset of input networks. Analyses were performed using the R package MSCquartets (Rhodes et al., 2021, 2026b).

### 2.7 HIV data

HIV-1 is a natural benchmark for the consensus method studied here. Its pandemic group M lineages have been diversifying for over 100 years, resulting in nine distinct subtypes (A, B, C, D, F, G, H, J, and K), which have in turn split into sub-subtypes (e.g., F1 and F2) (Hemelaar, 2012). Furthermore, the group M sequences have undergone recombination (Robertson et al., 1995, 2000), producing circulating recombinant forms (CRFs) of the virus. The parental subtypes of CRFs can be identified using sequence-based methods (e.g., Hidden Markov Models; Schultz et al. (2012)), the results of which are manually verified and stored in regularly updated databases (Los Alamos National Laboratory, 2023). However, the deep branching of several subtypes is still debated (Buonaguro et al., 2007; Tongo et al., 2016), probably due to complex histories involving recombination and other biological mechanisms. This is precisely the type of situation in which network inference and consensus methods can be informative. Furthermore, some of their conclusions can be compared with the latest knowledge available in this field.

Following Holtgrefe et al. (2025b), we studied CRF03 A6B, also known as KAL153. This is a recombinant of sub-subtype A6 and subtype B, which was first identified among injecting drug users in Kaliningrad in the 1990s. It was one of the first CRFs to be identified and studied (Robertson et al., 2000). It is now prevalent in countries of the former Soviet Union (Lebedev et al., 2020). We used the 2023 RIP reference alignment from the LANL HIV Sequence Database (Los Alamos National Laboratory, 2023), which provides, for each group, subtype, sub-sub-type and CRF, up to four representative reference sequences. As input we used the reference sequences of the 14 pure group M (sub-)subtypes: A (sub-subtypes A1, A2, A3, A4, and A6), B, C, D, F (sub-subtypes F1 and F2), G, H, J and K, plus the outgroup N. This represents 15 taxa and 53 reference sequences (two to four per taxon), together with a single CRF03 A6B reference sequence (03 A6B.RU.97.KAL153 2.AF193276)

Because Squirrel (Holtgrefe et al., 2025b) takes a single aligned sequence per taxon, while most taxa have several references, we ran a resampling analysis of 1,000 replicates to assess the robustness of the reconstruction using a consensus approach. For each replicate, we (1) drew one sequence uniformly at random from each of the (sub-)subtype’s references, including the N outgroup, while always retaining the same CRF03 A6B reference sequence; and (2) ran Squirrel with N as outgroup to infer a rooted level-1 network from the resulting alignment, taking *λ* = 0.3 for consistency with the HIV analysis in Holtgrefe et al. (2025b). As replicates draw from overlapping subsets, the 1,000 networks are not independent but together capture the sampling variability across reference sequences. From these, we built level-1 outgroup-rooted networks using the L1N, HMS and HWC consensus methods.

We also ran Squirrel in its unrooted setting, without designating N as outgroup, which returns semi-directed level-1 networks directly, and applied L1N in its unrooted setting to the resulting 1,000 networks. The consensus is reported in the Supplementary Material.

As a reference phylogeny for the non-recombinant component of the data, we inferred a maximum likeli-hood tree with RAxML-NG (Kozlov et al., 2019) under the GTR+Γ substitution model using the reference-sequence alignment (53 sequences), excluding the CRF03 A6B sequence. We performed bootstrapping to assess the support of branches in this tree.

Finally, we repeated the entire procedure (the rooted, unrooted and tree analyses), with subtypes H and J excluded, leaving 13 taxa and all other settings unchanged. These are among the rarest and most sparsely sampled subtypes. There is no agreed placement for them in the literature, and they tend to distort the signal (see below). The results are reported in the Supplementary Material.

## 3 Results

### 3.1 Simulation results

As described above, we used 20 randomly generated level-1 networks, each with 21 taxa. Each network was used to generate 400 gene trees, which were then used to simulate the evolution of DNA sequences with an average gene length of 600 bp. These datasets were analyzed by Squirrel (Holtgrefe et al., 2025a) under four different settings. In the ‘Rooted’ setting, Squirrel was given knowledge of the outgroup taxon, while in the ‘Unrooted’ setting, this information was hidden, allowing for errors where the outgroup taxon could be included in some hybrid clade. In the ‘Full’ setting, Squirrel was given the full gene sequences, while in the ‘Third’ setting, it used only a third of each gene sequence, thus relying on less information. We concatenated the (full or third) gene MSAs to apply Squirrel. We then used a gene-based block bootstrap to obtain 100 input bootstrap networks, which we gave to the various consensus methods. For each of the 20 model networks and for each setting (e.g., ‘Full & Rooted’), the experiment was replicated 10 times, thus creating 10 samples of 100 input bootstrap networks each. We assessed the performance of the various methods from two angles: (1) average dissimilarity to the input networks, which shows how informative the consensus is with respect to these inputs, and (2) accuracy in reconstructing the model network used to simulate the data. As previously mentioned, these two viewpoints are correlated, provided that the inference method (here, Squirrel) is not excessively biased. The performance metrics and dissimilarities were measured with respect to blob partitions, circular orders, hybrid clades in blobs of size greater than 4, and all (both redundant and non-redundant) splits, as well as a global mean of these four measures. We compared our L1N consensus approach with the HMS (Huber et al., 2023) and HWC (Nielsen et al., 2023) consensuses, which are only applicable in the ‘Rooted’ setting. We also compared the results of the various consensus methods to the average dissimilarity between the input bootstrap networks and the corresponding true model network (‘TRUE’ measurements), and to the average accuracy metrics of the input networks (‘Input Nets’ measurements). All these metrics were averaged for each setting, across the 20 model networks and 10 replicates.

The results for dissimilarity to input networks are shown in Figure 7 (Jaccard distance) and Supplementary Figure S13 (symmetric distance). The main observations from Figure 7 are as follows:

**Figure 7:**
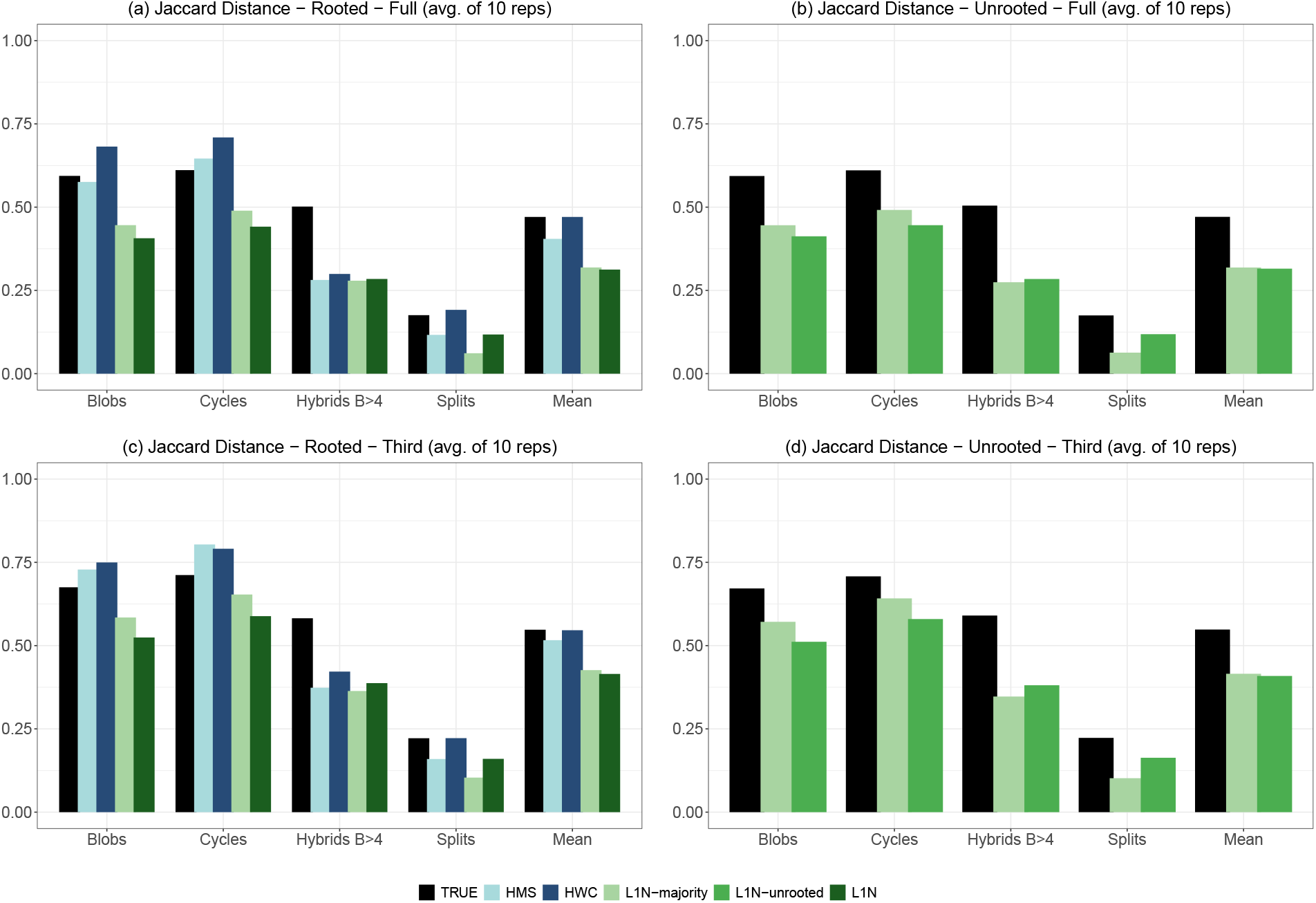
Average Jaccard distance to the input networks. For each method and setting, we provide the average Jaccard distance to the input networks with respect to blob partitions (Blobs), circular orders (Cycles), hybrid clades in blobs of size greater than 4 (Hybrids B*>*4), and all splits (both redundant and non-redundant), as well as the mean of these four measurements. (a) ‘Full’ genes and ‘Rooted’ setting; (b) ‘Full’ genes and ‘Unrooted’ setting; (c) ‘Third’ of each gene and ‘Rooted’ setting; (d) ‘Third’ of each gene and ‘Unrooted’ setting. For the ‘Rooted’ setting, we compare the HMS, HWC, and L1N consensuses. Moreover, we provide the results of L1N-majority, in which only components (e.g. blobs) with over 50% support are retained (standard support is applied to splits). In the ‘Unrooted’ setting, HMS and HWC consensuses are not applicable; we provide the results of L1N-unrooted, in which neither Squirrel nor L1N is given knowledge of the outgroup, and L1N-majority, which is also non-informed about the outgroup in that case. For all four settings, we provide the average Jaccard distance between the input networks and the ‘TRUE’ model network that was used to generate the data. This provides a basis for comparison, particularly in the ‘Unrooted’ setting, since we would expect any consensus method to be closer to the input networks than the ‘TRUE’ model is. See text for further details and explanations.

- As expected, all distances are larger in the ‘Third’ setting than in the ‘Full’ setting. Using only one-third of the gene sequences increases the variability of the networks inferred by Squirrel from the bootstrapped MSAs.
- Conversely, we observe little difference between the ‘Rooted’ and ‘Unrooted’ settings for L1N variants. Knowledge of the outgroup appears to have a limited impact on the bootstrap networks inferred by Squirrel.
- For all methods, distances with respect to blob partitions and circular orders are greater than those with respect to hybrid clades and, especially, splits. Blob partitions and circular orders are complex components that are difficult to infer without error or misplaced taxa; hence, their greater variability. Conversely, hybrid clades and splits, in particular, are simpler components that can be more easily inferred with lower variability when using the large amounts of sequence data employed in this study.
- In the ‘Rooted’ setting, there is a clear gap between the HMS and HWC consensuses, and the L1N variants, which perform better. The HMS consensus tends to be slightly closer to the input networks than the HWC consensus is; both show, on average, distances to the inputs that are similar to the distances between the true model network and the inputs (as shown by the ‘TRUE’ black bars in Fig. 7). L1N and L1N-majority are closer to the inputs than the true network is, for all measurements. This is somewhat expected, as their fundamental principle is to maximize support for the four component types, which results in low component-based distances to the inputs.
- In the ‘Unrooted’ setting, we observe the same: both L1N-unrooted and L1N-majority are clearly closer to the inputs than the true network is, for all measurements.
- On average, there is relatively little difference between L1N and L1N-majority with respect to Jaccard distance (Fig. 7). Globally, they are similar in their distance to the inputs. However, we observe differences regarding splits, where L1N-majority tends to perform better than L1N. Conversely, we observe the opposite with blob partitions and circular orders. This is likely due to the uniform use of a 50% threshold in L1N-majority, which is better suited to splits (e.g., think of the majority-rule consensus tree) than to complex components, such as blob partitions and circular orders.
- Considering the symmetric distance (Supplementary Fig. S13), the results are globally similar to those observed with the Jaccard distance. As expected, due to the optimality property (see above), the L1N-majority method is the best for all measurements. HMS and HWC consensuses, which also follow the majority rule, have better measurements than using the Jaccard distance; on average, they clearly improve over the ‘TRUE’ measurements, which correspond to the symmetric distances between the true network and the inputs. On average, L1N consensus, which reconstructs fully resolved networks, is the second best method.

The accuracy results in reconstructing the true model network used to generate the data are shown in Figure 8 (precision, recall, and Jaccard index) and Supplementary Figure S14 (symmetric distance and F1 score). The main observations from Figure 8 are as follows:

**Figure 8:**
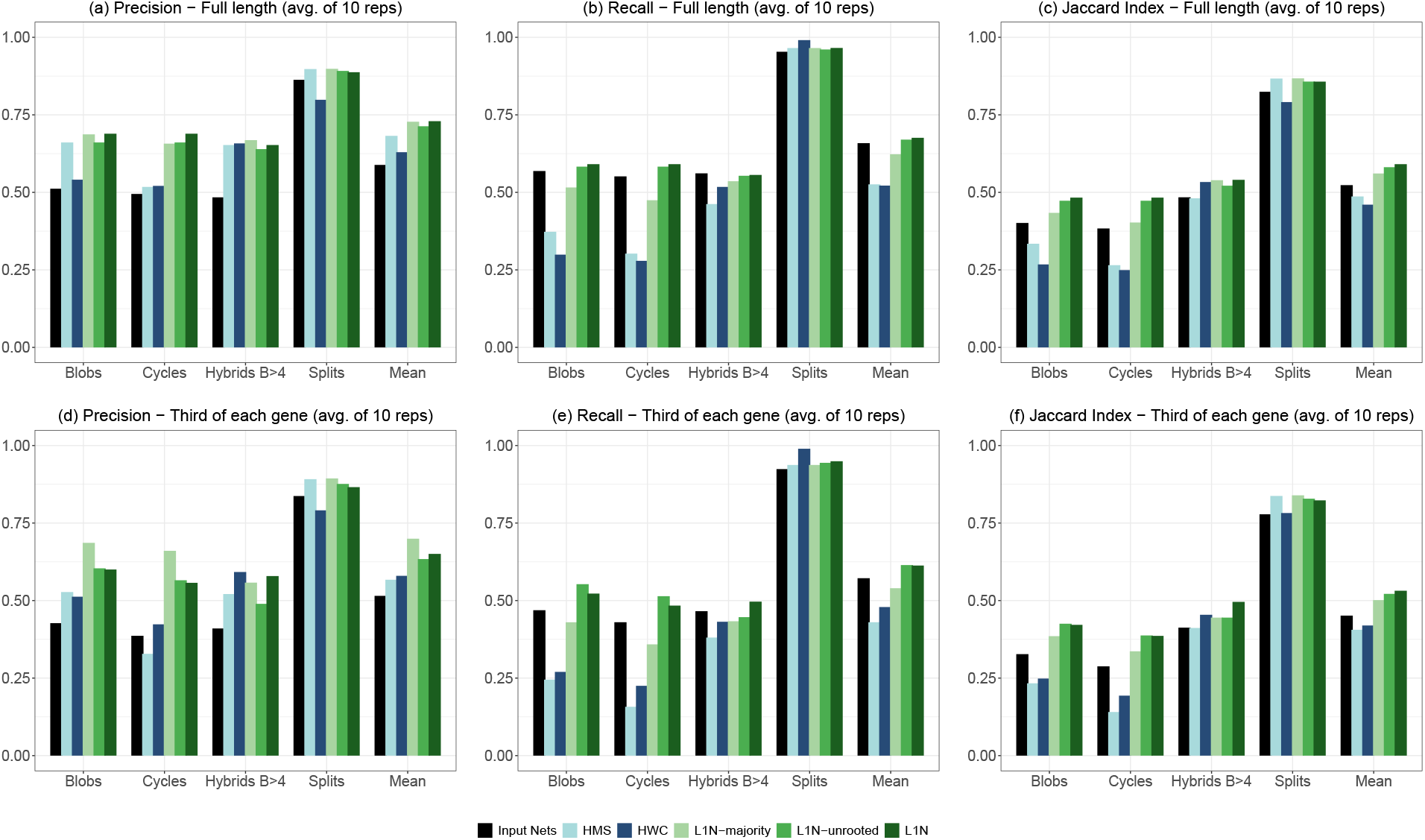
Accuracy in reconstructing the true model network. For each method, we provide the average precision (a,d), recall (b,e), and Jaccard index (c,f) to the true model network with respect to blob partitions (Blobs), circular orders (Cycles), hybrid clades in blobs of size greater than 4 (Hybrids B*>*4), and all splits (both redundant and non-redundant), as well as the mean of these four measurements. (a,b,c): ‘Full’ genes setting. (d,e,f): ‘Third’ of each gene setting. All results, except those of L1N-unrooted, are obtained in the ‘Rooted’ setting. In this setting, we compare the HMS, HWC, and L1N consensuses, as well as the L1N-majority variant, in which only components with over 50% support are retained. We also present the results of L1N-unrooted in the ‘Unrooted’ setting, in which neither Squirrel nor L1N is informed about the outgroup. Finally, we present the average accuracy measurements of the 100 input networks (Input Nets, in black), as we would expect a consensus network to be more accurate than the input networks on average. See the text and Supplementary Material for details.

- Similar to the distance-to-inputs results (Figure 7), accuracy decreases substantially in the ‘Third’ setting compared to the ‘Full’ setting for all methods and measurements except for splits where the difference is negligible. This is simply a consequence of Squirrel’s higher accuracy with full gene sequences. However, the splits are relatively easy to infer, even in the ‘Third’ setting, due to the amount of data used in this study.
- Accuracy is consistently much better for splits than for other components, across all methods and criteria. As seen in Figure 7, the accuracy (Figure 8) for hybrid clades falls between that of splits and that of blob partitions and circular orders.
- Consensus methods based on a majority rule, particularly HMS and HWC, and to a lesser extent L1N-majority, tend to have relatively high precision but lower recall. The latter is typically lower than the average recall of the input bootstrap networks (corresponding to the black ‘Input Nets’ bars in Fig. 8) except for splits that are inferred with high accuracy by all methods. This outcome is expected, as the majority rule discards poorly supported components. On average, the true model networks and the input bootstrap networks contain 3.35 and ~ 3.1 blobs, respectively, while HMS, HWC and L1N-majority contain ~ 1.5, ~ 1.4 and ~ 2.4 blobs, respectively (see Supplementary Table S1 for details). This lack of resolution decreases recall - particularly in the ‘Third’ setting, where the signal is lower and more components are poorly supported but it increases precision for the blob partitions and circular orders.
- The greedy methods (L1N and L1N-unrooted) do not perform this type of selection. They infer fully resolved networks with ~3.1 blobs on average, and produce more balanced results. They have similar to or higher precision and recall than do input bootstrap networks on average, for all measurements.
- Based on the Jaccard index, HMS and HWC perform similarly. They exhibit high error rates with respect to blob partitions and circular orders due to their low recall and resolution. However, they are fairly accurate with respect to splits, with hybrid clade recovery being intermediate. Overall, their ‘Mean’ Jaccard indices are lower than those of the input networks.
- The three L1N variants show similar performance with respect to the Jaccard index for all components. On average, L1N-majority lags slightly behind the other two, while L1N is slightly better than L1N-unrooted, as expected.
- The F1 score results (Supplementary Figure S14) are nearly identical to the Jaccard index results. There is a clear gap between HMS and HWC, and L1N variants. The former have a lower ‘Mean’ accuracy compared to input networks on average, while the L1N variants improves on input networks, particularly the fully resolved L1N consensus.
- The results regarding the symmetric distance (Supplementary Figure S14) are more balanced, particularly in the ‘Third’ setting, where the signal is low. All methods improve upon input bootstrap networks. Thanks to majority-rule selection, the HMS and HWC consensuses are close to the fully resolved greedy L1N and L1N-unrooted consensuses. As expected, the L1N-majority method performs best by discarding poorly supported components.

In summary, our simulations show that the L1N variants outperform the existing HMS and HWC consensuses in the ‘Rooted’ setting for nearly all criteria. This is true in terms of both representing the input networks and accurately reconstructing the true model network. In a few cases, such as with regard to splits, all methods demonstrate similar good performance. In the ‘Unrooted’ setting, L1N consensus is the only available method, to the best of our knowledge. The results are nearly identical to those in the ‘Rooted’ setting, which demonstrates the strength and applicability of the L1N approach.

### 3.2 Leopardus data

Figure 9 shows the L1N, HWC, and HMS consensus networks for the Leopardus data. L1N analyses were performed using all 755 bootstrap networks, and those 476 bootstrap networks consistent with the Puma outgroup as non-hybrid (Fig. 9 (a) and (b)). The 476 networks were rooted along the external edge leading to Puma and used as input to HMS and HWC, which returned the same rooted consensus network (Fig. 9(c)).

**Figure 9:**
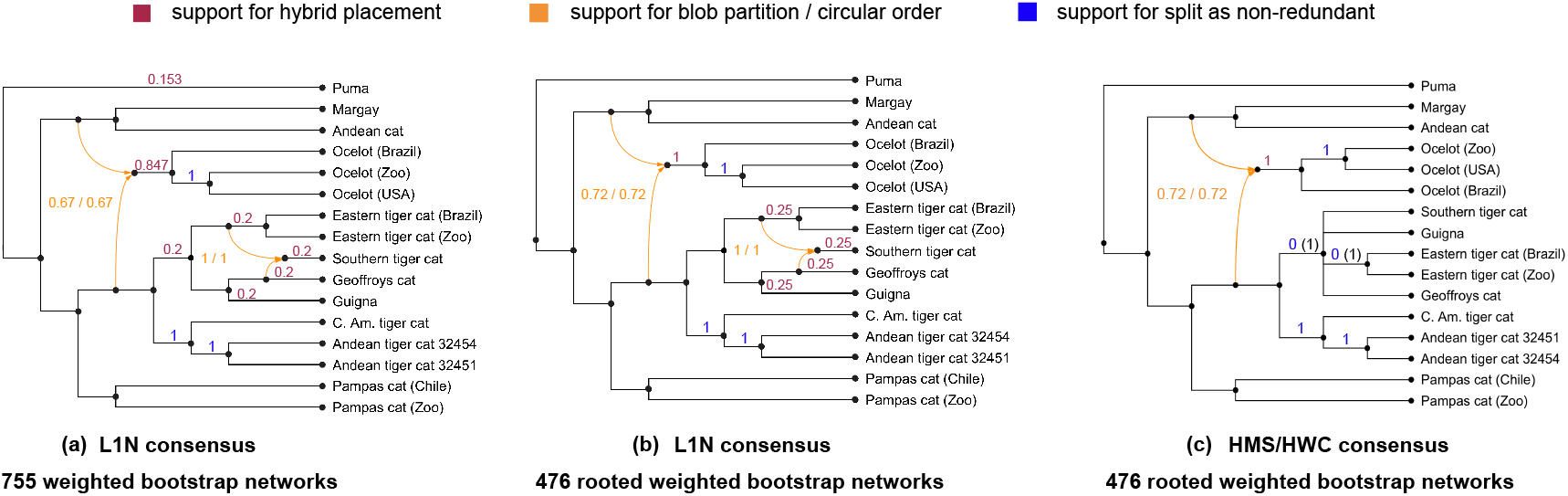
Consensus networks for the Leopardus genus. **(a)** The L1N consensus unrooted (semi-directed) network based on 755 unrooted, weighted bootstrap networks inferred using TINNiK and NANUQ^+^. The L1N consensus rooted network based on 476 rooted, weighted networks. These were obtained from the previous 755 networks by retaining only those that permitted rooting along the external edge leading to the Puma outgroup. **(c)** The less-resolved HMS and HWC consensus network(s), which are identical and were obtained using the same 476 rooted, weighted input networks. Magenta: Support for the hybrid clades (0 values are not indicated). Orange: Support for the blob partitions and their associated circular order. Blue: Support for non-redundant splits computed from the input networks. In (c), two splits have non-redundant split support value 0, since they always appear as redundant splits in the input (see (b)); however, their standard support (black, in parentheses) is 1. Networks were plotted with PhyloSketch and PhyloPlots (Ané, 2025; Huson, 2025). See text and Supplementary Material for details.

Both L1N consensus networks are identical to the one published by Lescroart et al. (2023), who used SNaQ (Solís-Lemus and Ané, 2016) which returns level-1 semi-directed networks. This network was also found using NANUQ^+^ (Allman et al., 2025) using a TINNiK starting ToB (as done here) from the original, non-resampled set of 16,338 local trees. This network contains two well-supported 5-cycles, each associated with a unique circular order. Since the circular orders are unique, the blob partition support and circular order support values are equal (top-left cycle: 0.67 in (a) and 0.72 in (b); bottom-right cycle: 1.0). In the first cycle (top left), support for the Ocelots as a hybrid clade is quite high, which is consistent with previous analyses of these data (Allman et al., 2025; Lescroart et al., 2023). In the unrooted L1N consensus (a) using all bootstrap networks, Puma (the outgroup) is the only block apart from the Ocelots with non-zero support as a hybrid in this cycle. This surprising outcome might be explained by model violation, or uncertainty of signal in the data, and merits further investigations. Once bootstrap networks inconsistent with rooting at Puma are removed from the analysis, as was done in (b), the Ocelots have full support as the hybrid clade.

In regard to the second cycle in L1N consensuses (bottom right, Fig. 9(a) and (b)), there is no signal for the determination of the hybrid clade, with all taxon blocks having equal hybrid clade support. This gives additional evidence to the hypothesis, stated in Allman et al. (2025), that the level-1 assumption might be inadequate to explain gene flow between these taxa.

The two L1N consensus networks (Fig. 9(a) and (b)) and their associated support values capture and quantify features of the input networks. In particular, they point out agreement among the bootstrap networks (hybrid origin of the Ocelots, reticulate history within the clade Guigna, Geoffroy’s cat, and Southern and Eastern tiger cats) and uncertainty (no identification of the hybrid(s) among these species).

The HMS and HWC consensus network(s) (Fig. 9(c)) recover the same top-left blob partition, circular order, and Ocelot hybrid clade as the L1N consensus (b), and the support values computed using PhyloSummaries (Ané and Sharma, 2026) are identical. However, this HMS/HWC consensus does not have a second 5-cycle, but rather a degree-5 node. This marks a clear loss of information when compared to the input networks, all of which contain the same cycle and circular order, indicating the presence of reticulation. Because the input networks vary greatly with respect to the hybrid clade in this cycle, the HMS/HWC consensus lacks resolution. Indeed, these two methods are based on hardwired clusters (HWC) and set pairs (HMS), rather than structural components of the input networks. Consequently, neither HWC nor HMS explicitly accounts for common blob partitions or reconstructs the corresponding cycles by associating hybrid clades and circular orders with blob partitions, as L1N does. Using the component-based Jaccard distance, the HMS/HWC consensus has a mean distance of 0.63 to the input networks, which is nearly twice that of the L1N consensus (0.33), illustrating the loss of resolution.

Overall, both the simulation and *Leopardus* analyses demonstrate some advantages of the component-based approach implemented in the L1N consensus, and the importance of quantifying variability using component support values. More features of the input networks are preserved in the L1N consensus, with some measure of uncertainty, than in the HMS/HWC networks.

### 3.3 HIV data

Figure 10 shows the L1N, HWC and HMS consensus networks for the HIV data, each based on the same 1,000 outgroup-rooted networks, together with the RAxML-NG tree inferred from the 53 reference sequences with the A6B recombinant removed.

**Figure 10:**
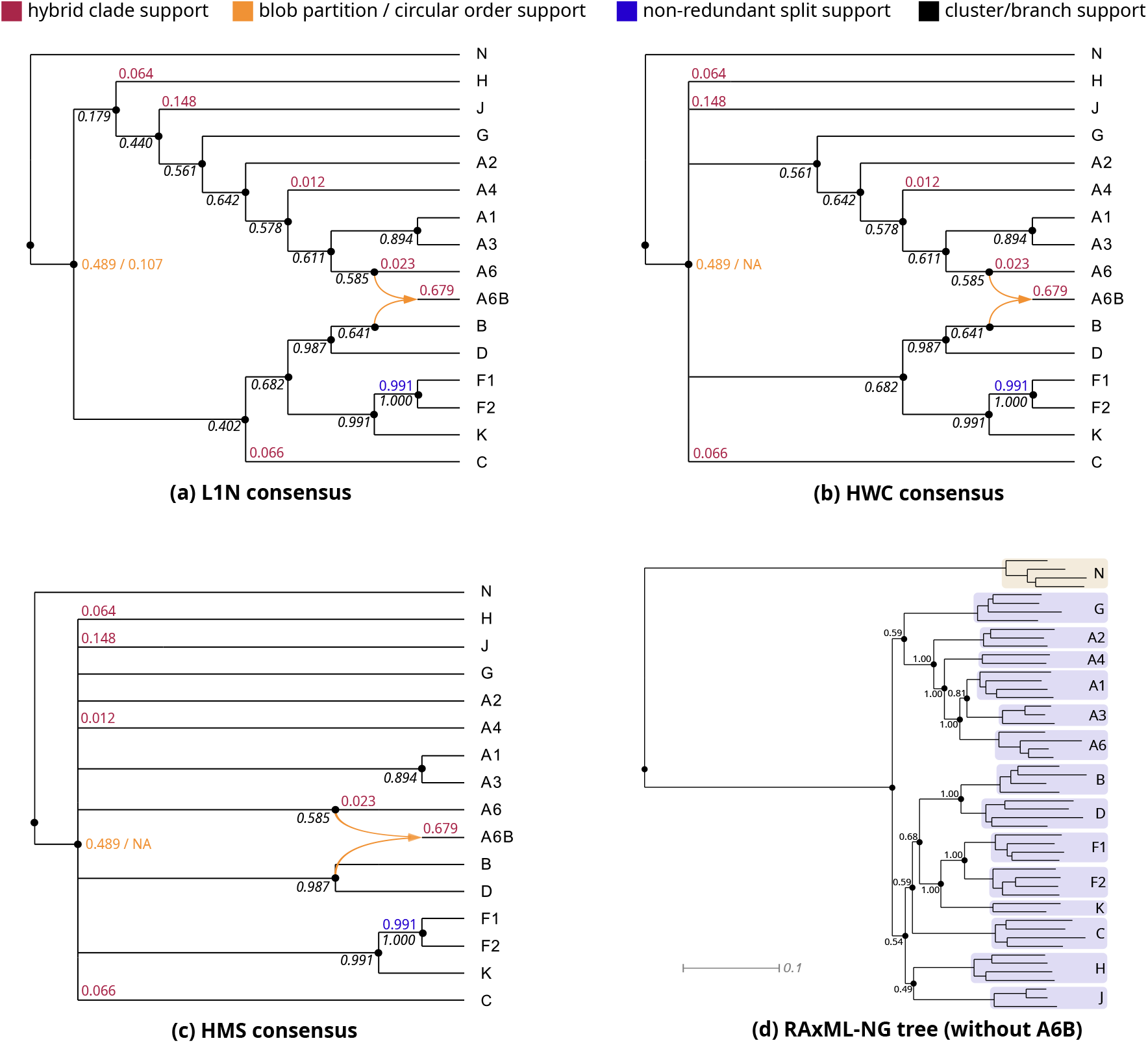
Consensus networks for the HIV-1 group M subtypes. The L1N consensus network (a) on 1,000 rooted networks inferred with Squirrel (Holtgrefe et al., 2025b), the less-resolved HWC (b) and HMS (c) consensus networks, together with the RAxML-NG (Kozlov et al., 2019) tree inferred from the 53 reference sequences with the A6B recombinant removed (d). In addition to the hybrid clade, blob partition, and non-redundant split support values, we report the cluster support for the internal branches. In (a), (b) and this is the proportion of the 1,000 input networks in which the corresponding hardwired cluster occurs. In (d), we use the standard Felsenstein bootstrap branch support (Felsenstein, 1985) from the RAxML-NG analysis. We omit hybrid clade support below 0.01 in (a), (b) and (c), and in (d) the branch support on the branches leading to each (sub-)subtype and those between their representative sequences.

Several relationships are recovered consistently and identically by all three consensus methods: the (A1,A3) grouping within the A clade, the (B,D) clade, the (F1,F2) clade, and the (F1,F2,K) clade. These match the established subtype phylogeny of (Hemelaar, 2012) and are likewise recovered with very high bootstrap support in our own RAxML-NG analysis of the sequence alignment with the CRF removed. All three consensus networks recover the A6B recombinant with strong support (0.679), and its placement is consistent with its documented recombinant ancestry (Robertson et al., 2000), although HMS is less resolved here: its second A6B parent is a high-degree node, the ancestor of B and D, rather than B itself.

Support is noticeably weaker for the deep structure of the network. All three consensus methods return the same blob partition for the unique cycle in the respective network, supported at 0.489, but differ in the extent to which they resolve the cycle: L1N returns a circular ordering, though supported at only 0.107, while HWC resolves less, attaching H, J and C at a high-degree ancestral node, and HMS leaves almost the entire cycle unresolved, so the latter two methods do not return a unique circular ordering at all. While the L1N and HWC consensuses recover most of the tree structure, the HMS consensus network is poorly resolved. It does not recover the (B, D, F, K) clade (Hemelaar, 2012) and the subtree corresponding to the A subtypes. Neither the HMS nor the HWC consensus recovers the (B, C, D, F, K) clade, which is seen in both the L1N consensus network and the RAxML-NG tree.

Despite the disagreement in input networks, the L1N consensus is the single most frequent input network, although it occurs in only about 10% of replicates. Thus, the L1N greedy approach returns a network that Squirrel actually inferred rather than a composite supported by none of the replicates. The HMS and HWC consensus network are not among the input networks, as their high-degree nodes make them non-binary while every input network is binary. The difference in component-based Jaccard distance to the input networks is small among the three consensuses (HMS/HWC: 0.58; L1N: 0.55), unlike the *Leopardus* dataset. This can be explained by the identical blob partition, hybrid clade, and splits among the three consensuses, as well as the low support (10%) of the L1N circular order. See the Supplementary Material for details on the distance calculation with non-binary networks.

For each branch, we calculated the support for its cluster of descendant taxa (cluster support). We attribute the weak support values in the deep part of the cycle (Figure 10) to conflicting phylogenetic signal involving subtypes J and H. The group M subtypes first radiated in a complex fashion with recombination playing a role (Buonaguro et al., 2007; Tongo et al., 2016), so deep branching order within group M is intrinsically hard to resolve, as is apparent from the rather low bootstrap supports for the deep branches in our RAxML-NG tree. This is compounded for J and H, which are among the rarest subtypes (each *<* 1% of global infections) and hence sparsely sampled (Indihar et al., 2025), with no agreed-upon placement for them in the literature. Our RAxML-NG analysis confirms this, placing J and H differently from Hemelaar (2012). As this tree was inferred without the CRF, the uncertainty is really a property of the subtypes themselves and not of the addition of the recombinant sequence.

To confirm that J and H are one of the drivers of this conflict, we repeated the entire analysis with the two subtypes removed (see the Supplementary Material). L1N, HWC, and HMS all return the same consensus network, so the differences between the methods disappear once the conflicting taxa are removed. Blob partition support rises to 0.567 and circular-ordering support to 0.533, and the A6B hybrid is recovered at 0.999 against 0.679 on the full data; the consensus network now coincides with 53.3% of the input networks. Apart from a slight decrease for the (A1,A3) clade, all cluster supports increase significantly as well. Removing J and H also improves the positioning of the outgroup N, which is further evidence that the two subtypes are a source of the conflict: the bipartition of the taxa induced by the root’s child (i.e., the two hardwired clusters), takes 64 distinct forms across the full-data replicates, the most frequent accounting for only 18.6%, whereas a single bipartition occurs in 94.6% of the replicates without J and H. The placement of J and H therefore appears to inject substantial conflict into the data, in line with them being rare subtypes of uncertain phylogenetic placement.

We also repeated the analysis with Squirrel run in its unrooted setting, which returns semi-directed networks directly and does not assume N to be the outgroup. The consensus network obtained this way is topologically identical to the L1N network of Figure 10, with N receiving a hybrid support of 16.4%, similar to the Puma in Figure 9(a), which again merits further investigation, and the other support values differing by less than 0.04 throughout (see the Supplementary Material).

Taken together, this HIV analysis illustrates the advantages of L1N: it returns a fully resolved level-1 network that Squirrel itself inferred with interpretable support values. HWC and especially HMS lose resolution and, for the second, precision about the parents of A6B recombinant. The support values indicate the structural features for which this resolution is warranted. Thus, the weakly determined deep ordering of taxa around the cycle is reported as such rather than being silently collapsed into a high-degree node.

## 4 Discussion

In this paper, we introduced a framework for constructing consensus summaries of phylogenetic networks, applicable to both rooted and semi-directed networks. Our approach first constructs a consensus tree of blobs (ToB), taking as input any networks of arbitrary complexity. Building on this approach, we developed L1N consensus, which takes a collection of level-1 networks (rooted or semi-directed) as input and returns a level-1 consensus network, with support values for its components, including blob partitions, circular orders, hybrid clades, and splits. The methods are implemented in PhyloSummaries (Ané and Sharma, 2026), building on PhyloNetworks (Solís-Lemus et al., 2017). Our simulation results demonstrate that the L1N variants generally outperform the existing HMS and HWC consensus methods (Huber et al., 2023; Nielsen et al., 2023) in the rooted setting, in terms of both representing the input networks and recovering the true model network. Although the methods perform similarly well for some criteria, such as split recovery, the overall results support the effectiveness of the L1N approach. In the semi-directed setting, L1N is, to the best of our knowledge, the only available consensus method, and its performance is nearly identical to that in the rooted setting. Analyses of the Leopardus and HIV datasets further illustrate the practical utility of the approach for summarizing collections of inferred networks.

An important contribution of this work is the generality of the ToB consensus framework. Although a tree of blobs captures only part of a network’s structure, it provides a meaningful summary of its evolutionary history. Indeed, recent work has shown that accurate ToB inference is key to accurate inference of level-1 networks (Dai and Molloy, 2026), and ToBs are identifiable from a broad range of data types, including for networks of arbitrary level (Allman et al., 2022; Xu and Ané, 2023). Thus, a ToB may be well supported by the data even when the detailed reticulation structure within individual blobs is uncertain.

Our framework can also be applied to ToBs inferred directly from data, without reconstructing the full networks. Several methods have recently been proposed for this purpose (Allman et al., 2024b; Dai et al., 2026; Rhodes et al., 2026a). When the inferred nodes are labeled as blob or divergence nodes, as in ECToBlob (Rhodes et al., 2026a), our consensus ToB method can summarize these estimates directly.

The majority-rule consensus ToB has a median property, providing a formal justification for its use as a summary of input ToBs. Extending this property to network consensus is more challenging because blob partitions, circular orders, hybrid clades, and splits must satisfy compatibility constraints. Although our L1N-majority method minimizes a component-based symmetric difference, its output need not correspond to a valid phylogenetic network. More generally, defining a majority-rule consensus network is not straightforward. For example, a blob partition may occur in more than half of the input networks even if no associated circular order has majority support. Similarly, a hybrid clade may have majority support, yet be absent from every blob partition that has majority support itself. Thus, the majority-supported components need not combine into a valid level-1 network.

We therefore report support values for all components, providing a more informative summary when support is distributed across competing network structures. Although our empirical results demonstrate the effectiveness of L1N, they do not establish that it minimizes a particular network distance or satisfies a general set of axiomatic properties. Establishing such formal properties for L1N consensus remains an important direction for future work.

Another direction for future work is to explore alternative consensus approaches. For example, distance-based consensus methods could seek a network that minimizes the aggregate distance to the input networks, using network dissimilarity measures beyond the component-based distances considered here (see, e.g. Huson et al. (2010); Marchand et al. (2025); Maxfield et al. (2024)). Alternatively, quartet- or quarnet-based methods could construct a consensus network from the frequencies of induced 4-taxon relationships across the input networks. However, since this information does not generally uniquely determine a phylogenetic network (see, e.g., Huber et al. (2014)), such approaches may not fully capture the reticulation structure of the input networks, unless certain structural constraints (such as bounded level) are imposed.

Relatedly, extending consensus methods to networks of higher level remains an important challenge. The generality of our ToB framework means that it can summarize networks of arbitrary complexity, but constructing a consensus network that retains detailed reticulation structure is a different problem. In higher-level networks, overlapping and more complex interactions among reticulation cycles introduce additional compatibility constraints and make it more difficult to identify a representative network. Notice, however, that accurate and scalable inference of higher-level networks itself is a challenging problem that is currently actively investigated. One possible direction for consensus methods could be to extend the component-based approach of L1N-majority, allowing support to be assessed for increasingly complex networks and key biological components, such as hybrid clades, without requiring all structural features to be assembled into a single fully resolved network.

## 5 Data and resource availability

All materials required to reproduce the analyses presented in this study are available at https://github.com/HectorBanos/NetworkConsensus. The repository includes Perl implementations of the HMS and HWC consensus methods and a Common Lisp implementation of the L1N consensus method; a Julia implementation of L1N is also available through PhyloSummaries (Ané and Sharma, 2026). The repository further contains the 20 model networks used in the simulation study, the simulated data and network inference and consensus results, and all files required to reproduce the empirical analyses of the HIV and *Leopardus* datasets.

## 6 Author contributions statement

This work started as a joint project at the NSF-funded institute ICERM. OG laid out the direction of the group project and oversaw all aspects of the research, including a first implementation of consensus methods in Lisp. CA implemented our novel algorithms in Julia, and KW implemented the HMS and HWC methods in Perl. HB oversaw the simulation studies, NH the HIV analysis, and EA the applications to Leopardus data. OG and KW wrote the first draft of the manuscript and supplementary material. All authors were involved in discussions and project development, and in editing the final manuscript.

## 7 Acknowledgements

This research started at the Institute for Computational and Experimental Research in Mathematics in Providence, RI, during the “Theory, Methods, and Applications of Quantitative Phylogenomics” program in Fall 2024, supported by grant National Science Foundation DMS-1929284. The NSF also funded this work through grants DMS-2051760, DMS-2554614 to ESA, DMS-2023239 to CA, DMS-2331660 to HB, and DMS-2554615 to KW. NH was supported by grant OCENW.M.21.306 of the Dutch Research Council (NWO). We thank Ayush Sharma for help with implementation and John A. Rhodes for initial discussions.

## Supplementary Material

### Consensus methods for trees of blobs and level-1 phylogenetic networks

We provide additional formal definitions and rigorous proofs of all mathematical statements underlying the proposed consensus methods. Furthermore, we present the algorithms employed by these methods, describe the simulation procedures, and include supplementary figures.

## 1 Definitions

Throughout, *X* denotes a non-empty finite set of *taxa*, which are typically *species*, but could also be populations or individuals more generally.

### 1.1 Phylogenetic networks and trees of blobs

#### Definition 1.1

(Rooted phylogenetic network). A *rooted phylogenetic network N* ^+^ on *X* is a rooted, acyclic, directed graph without loops (edges from a vertex to itself) that satisfies the following conditions:

i. The (unique) *root* has in-degree zero and out-degree at least two.
ii. The set of vertices with out-degree zero is in bijective correspondence with *X*; these vertices are called *leaves*.
iii. All other vertices are of two types:
  - *Tree vertices* (in-degree one, out-degree at least two).
  - *Reticulations* or *hybrid* vertices (in-degree at least two, out-degree one).

If “at least” is replaced by “precisely” in the degree conditions, *N* ^+^ is called *binary* or *fully resolved*.

Due to the bijective correspondence between the leaves of *N* ^+^ and the taxa in *X*, the set is *X* sometimes referred to as the *leaf set* of *N* ^+^, depending on the context. An edge directed into a reticulation is a *reticulation edge*; all other edges are *tree edges*.

#### Definition 1.2

(Semi-directed phylogenetic network). A *(binary) semi-directed phylogenetic network N* on *X* is a partially directed graph obtained from a (binary) rooted phylogenetic network *N* ^+^ by undirecting all tree edges and suppressing the former root.

Reticulations and reticulation edges are defined in *N* analogously to *N* ^+^. A *(binary) unrooted phylogenetic X-tree T* is a (binary) semi-directed network on *X* with no reticulations.

#### Definition 1.3

(Blob; tree of blobs). A *blob* of a rooted or semi-directed phylogenetic network is a maximal connected subgraph that contains no cut edges (edges whose removal disconnects the graph). A blob that consists of a single vertex is *trivial*. A blob is said to be a *degree-m* blob if it is incident to *m* cut edges of the network. *Contracting* a blob means replacing it with a single vertex. The *tree of blobs* (or ToB for short) *T* (*G*) of a phylogenetic network *G* on *X* is the phylogenetic tree with leaf set *X* obtained by contracting every blob of *G*, and then suppressing non-root vertices of degree two, if any. If *N* ^+^ is a rooted network, then *T* (*N* ^+^) is a rooted tree. If *N* is semi-directed, then *T* (*N*) is unrooted tree.

In the following, we focus on a semi-directed network *N*. Vertices of *T* (*N*) arising from contracted cycles are called *blob vertices* or *blob nodes*. All other non-leaf vertices are called *divergence vertices* or *divergence nodes*. With these node labels, *T* (*N*) is called a *labeled tree of blobs*.

For our network consensus method, we consider binary semi-directed networks *level-*1 networks, that is, networks in which each reticulation belongs to exactly one undirected cycle (ignoring the directions of reticulation edges). In particular, blobs correspond to simple underlying cycles for these networks (see Fig. S1(b)).

**Figure S1:**
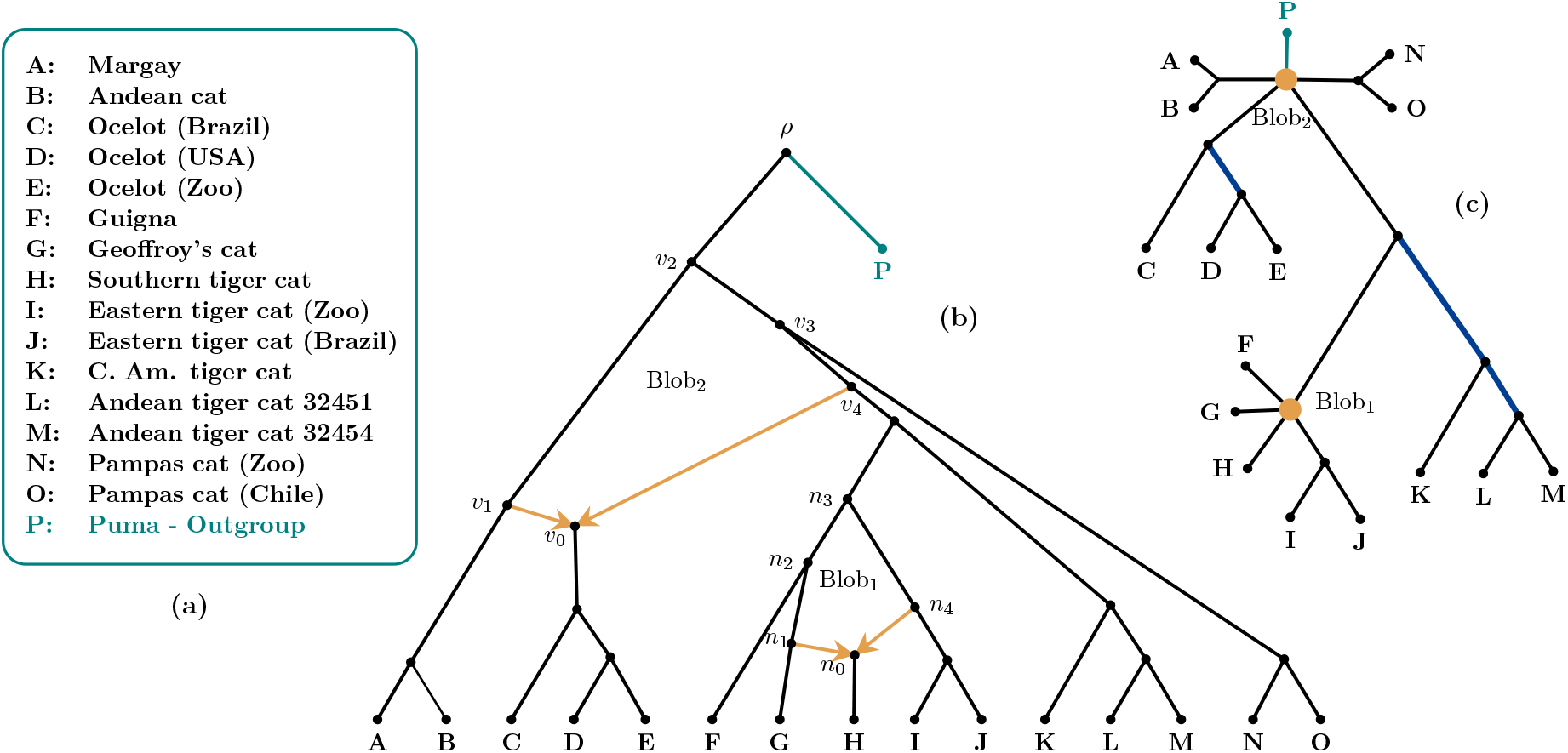
Level-1 phylogenetic network of Neotropical small felids (species names given in (a)) comprised in the genus *Leopardus* [14] (b) together with its induced tree of blobs (c). See main body of our manuscript for details.

### 1.2 Splits

An *X*-split is a bipartition of *X* into two non-empty subsets *A* and *B* (called *blocks*). We denote the *X*-split whose blocks are *A* and *B* by *A* | *B* or *B* | *A*. An *X*-split is *trivial* if |*A*| = 1 or |*B*| = 1.

Two splits *A* | *B* and *C* | *D* of *X* are said to be *tree-compatible* (or simply *compatible*) if at least one of the intersections *A* ∩ *C, A* ∩ *D, B* ∩ *C*, or *B* ∩ *D* is empty. Equivalently, *A* | *B* and *C* | *D* are compatible if *A* or *B* is included in *C* or *D*.

Recall that a cut edge in a graph is an edge whose removal disconnects the graph. In a semi-directed network, a cut edge is an edge that does not lie on any cycle. The cut edges of a semi-directed phylogenetic network *N* on *X* naturally induce a collection of splits of *X*.

#### Definition 1.4

(Splits induced by a network). A cut edge *e* of a semi-directed phylogenetic network *N* on *X induces* the split *σ*_*e*_ = *A* | *B* of if *X* its removal disconnects the leaves labeled by elements in *A* from those labeled by elements in *B*. We use Σ(*N*) to denote the set of all splits of *X* induced by the cut-edges of the semi-directed network *N*.

As an example, consider the tree edge incident to the vertex *v*_0_ in the semi-directed network *N* shown in Figure S1(b). This edge is a cut edge, since its removal disconnects the network, and it induces the split {*C, D, E*} | (*X* − {*C, D, E*}).

The splits Σ(*T*) induced by an unrooted phylogenetic *X*-tree *T* are defined analogously, and the following classical result describes when a collection of splits of *X* is induced by a phylogenetic *X*-tree *T*.

#### Theorem 1.5

(Splits-Equivalence Theorem [3]). *Let* Σ* *be a collection of X-splits. Then, there is an unrooted phylogenetic X-tree T such that* Σ* = Σ(*T*) *if and only if the splits in* Σ* *are pairwise tree-compatible. Moreover, if such an unrooted phylogenetic X-tree exists, then T is unique*.

### 1.3 Blob partitions and circular orders

Let *T* be a tree of blobs on taxon set *X*, induced by a (possibly unknown) semi-directed phylogenetic network *N* on *X*. While the edges of *T* induce *X*-splits, each blob vertex *b* of *T* induces a partition of *X* into *p* ≥ 3 non-empty subsets *A*_1_, …, *A*_*p*_ (as for splits, referred to as *blocks*), where each subset corresponds to the leaf set of one of the subtrees incident with *b*. We denote this resulting partition by

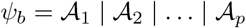

and call it a *blob partition*. Note that, as for splits, permuting the order of the sets *A*_*i*_ in the notation above does not change the blob partition.

As an example, consider the tree of blobs shown in Figure S1(c). The vertex labeled Blob_1_ induces the blob partition *ψ*_1_ = {*F*} | {*G*} | {*H*} | {*I, J*} | (*X* − {*F, G, H, I, J*}).

#### Remark 1.6.

If *N* is binary, then inner vertices of *T* with degree at least four correspond to reticulation structures in *N*. If *N* is non-binary, however, inner vertices of *T* of degree at least four may arise either from reticulation events or from genuine multifurcations in the underlying network. Moreover, in both cases (irrespective of whether *N* is binary), vertices of degree three in *T* may correspond either to tree vertices of *N* or to blobs (such as 3-cycles). This can be captured by assuming that each inner vertex of *T* is explicitly labeled as either a *blob vertex* or a *divergence vertex*.

Only those vertices labeled as blob vertices are considered to encode reticulate structure and thus induce meaningful partitions of *X*. Inner vertices that are not labeled as blob vertices are interpreted as genuine multifurcations. This labeling may be inferred from external information or specified by the user. In a consensus setting, this labeling is induced by the input networks, depending on whether a given vertex arises from the contraction of reticulate structure or not.

#### Definition 1.7

(Blob partitions in a tree of blobs). Let *T* be a tree of blobs on taxon set *X*, together with a designated set *B*(*T*) of inner vertices of *T* designated as blob vertices. Then

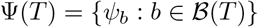

denotes the *set of all blob partitions* of *T*.

Recall that the blob vertices in a tree of blobs *T* induced by some binary semi-directed level-1 network *N* correspond to cycles in *N*. Let *b* be a blob vertex of *T*, and let *C*_*b*_ denote the corresponding cycle in *N*. While *b* induces a partition *ψ*_*b*_ = *A*_1_ |*A*_2_| …| *A*_*p*_, the cycle *C*_*b*_ carries additional information in the form of a circular ordering of these blocks.

#### Definition 1.8

(Circular partition and hybrid clade). Let *b* be a blob (cycle) of a binary semi-directed level-1 network *N*, and let {*A*_1_, …, *A*_*p*_} be the blocks attached to *b* via cut edges. The *circular partition* of *b* is a circular ordering *C*_*b*_ = (*A*_1_, …, *A*_*p*_) of these blocks, defined up to rotation and reversal. Moreover, the unique block *A*_*i*_ connected to the unique reticulation of *b* is called the *hybrid clade* of *b*.

For example, for the cycle labeled Blob_2_ in the network shown in Figure S1(b), the induced circular partition is ({*C, D, E*}, {*F, G, H, I, J, K, L, M*}, {*N, O*}, {*P*}, {*A, B*}), and {*C, D, E*} is the hybrid clade. Note that the circular ordering of the taxon blocks and the uniqueness of the hybrid clade of *b* requires that each node in the cycle has degree 3: adjacent to 2 edges in the cycle, and a single cut-edge outside the cycle that connects the node to a single taxon block. If *N* is binary, this condition is met. However, we can relax this assumption, to only require that nodes in cycles be of degree-3.

## 2 Compatibility of blob partitions and splits

While blob partitions capture the higher-order structure associated with reticulation, a tree of blobs also encodes information through its edges. Each edge induces a split of the taxon set, but some of these splits are already captured by the blob partitions and therefore do not contribute additional information. We now formalize the notion of those splits that are informative in this sense.

### Definition 2.1

(Non-redundant splits in a tree of blobs). Let *T* be a tree of blobs on taxon set *X*. An *X*-split A|B induced by an edge of *T* is called *non-redundant* if it is not implied by any blob partition in Ψ(*T*); that is, there exists no blob partition *ψ* ∈ Ψ(*T*) such that *A* (and hence *B*) can be expressed as a union of blocks of *ψ*. We define

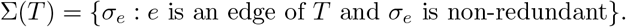

Observe that a split induced by an edge *e* of *T* is non-redundant if and only if *e* is not incident to any vertex designated as a blob vertex in *T*. Edges incident to such vertices induce splits that are already implied by the corresponding blob partitions and are therefore redundant. Since our goal is to introduce a consensus method for collections of trees of blobs, we note that a given split *σ* may be redundant in some input trees of blobs, but non-redundant in others. For instance, in Figure S6, the split {*A, B, C, D*} | {*E, F, G*} is redundant in *T*_1_, *T*_2_, *T*_3_, and *T*_5_, but non-redundant in *T*_4_.

Furthermore, note that a trivial split arises precisely from an edge incident to a leaf. Every tree of blobs, and more generally every phylogenetic network on *X*, displays all trivial-splits. As these carry no additional structural information, we henceforth restrict attention to non-trivial splits. For simplicity of notation and readability, we will refer to these simply as *X*-splits.

As an example, consider the tree of blobs depicted in Figure S1(c). In this case, there are three (non-trivial) non-redundant *X*-splits, namely {*D, E*} | (*X* − {*D, E*}), {*K, L, M*} | (*X* − {*K, L, M*}), and {*L, M*} | (*X* − {*L, M*}).

Given a collection of trees of blobs on the same taxon set *X*, together with their associated sets of blob partitions and non-redundant *X*-splits, we aim to construct a consensus tree of blobs that is consistent with these partitions and splits.

### Definition 2.2

(Compatibility of partitions and splits). Let Ψ* be a collection of partitions of *X*, each with at least three blocks (the blob partitions), and let Σ* be a collection of *X*-splits that are non-redundant with respect to Ψ*. We say that the pair (Ψ*, Σ*) is *blob-compatible*, or *compatible* for short, if there exists an unrooted tree *T* with leaf set *X* such that

- Ψ* is the set of blob partitions induced by the blob vertices of *T*, and
- Σ* is a subset of the set of edge-induced splits of *T* that are not implied by any partition in Ψ*.

*No restriction is imposed on additional partitions or splits induced by divergence vertices, respectively edges, of T*.

### 2.1 Pairwise compatibility conditions

Recall that two *X*-splits *σ*_1_ = *A* | *B* and *σ*_2_ = *C* | *D* can be induced by the edges of an unrooted phylogenetic *X*-tree *T* (and hence are blob-compatible) if at least one of the intersections *A*∩ *C, A*∩ *D, B* ∩ *C*, or *B* ∩ *D* is empty. Equivalently, *σ*_1_ = *A* | *B* and *σ*_2_ = *C* | *D* are tree-compatible if one of *A* or *B* is contained in *C* or *D*.

We next present a simple lemma that characterizes when a single blob partition of *X* and an *X*-split are compatible.

#### Lemma 2.3.

*Let ψ* = *A*_1_ | *A*_2_ | · · · | *A*_*p*_ *be a blob partition of X with p* ≥ 3, *and let σ* = *B* | *C be a non-trivial X-split that is non-redundant with respect to ψ. Then ψ and σ are compatible if and only if there exists a block A*_*i*_ *of ψ such that either*

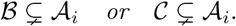

*In other words, one block of the split must be strictly contained within a single block of the blob partition*.

*Proof*. Suppose *ψ* = *A*_1_ | *A*_2_ | · · · | *A*_*p*_ and *σ* = *B* | *C* are compatible. By definition, there exists an unrooted tree *T* with leaf set *X* such that *ψ* is induced by an inner vertex *v* of *T* and *σ* by an edge *e* not incident to *v* (otherwise, *σ* would be redundant).

Let *T*_1_, …, *T*_*s*_ be the subtrees of *T* attached to *v*, with leaf sets *A*_1_, …, *A*_*p*_, respectively. Since *e* is not incident to *v*, it lies entirely within one of these subtrees, say *T*_*i*_. It follows immediately that all leaves of one part of the split, either *B* or *C*, are strictly contained in *A*_*i*_ (Fig. S2).

Conversely, suppose there exists a block *A*_*i*_ of *ψ* = *A*_1_ | · · · | *A*_*p*_ such that *A* ⊊ *B*_*i*_ (without loss of generality). We construct an unrooted tree *T* on leaf set *X* realizing both *ψ* and *σ* = *B* | *C* as follows (see Fig. S2 for an illustration): Let *v* be a vertex of degree *p* incident with *p* star subtrees on leaf sets *A*_1_, …, *A*_*p*_. By construction, *v* induces the partition *ψ*. To realize the split *σ*, insert an edge *e* inside the subtree corresponding to *A*_*i*_ so that removing *e* separates *B* from the remaining leaves of *A*_*i*_. Since *e* lies entirely inside *T*_*i*_, it is not incident to *v* and therefore does not interfere with the partition induced by *v*.

**Figure S2:**
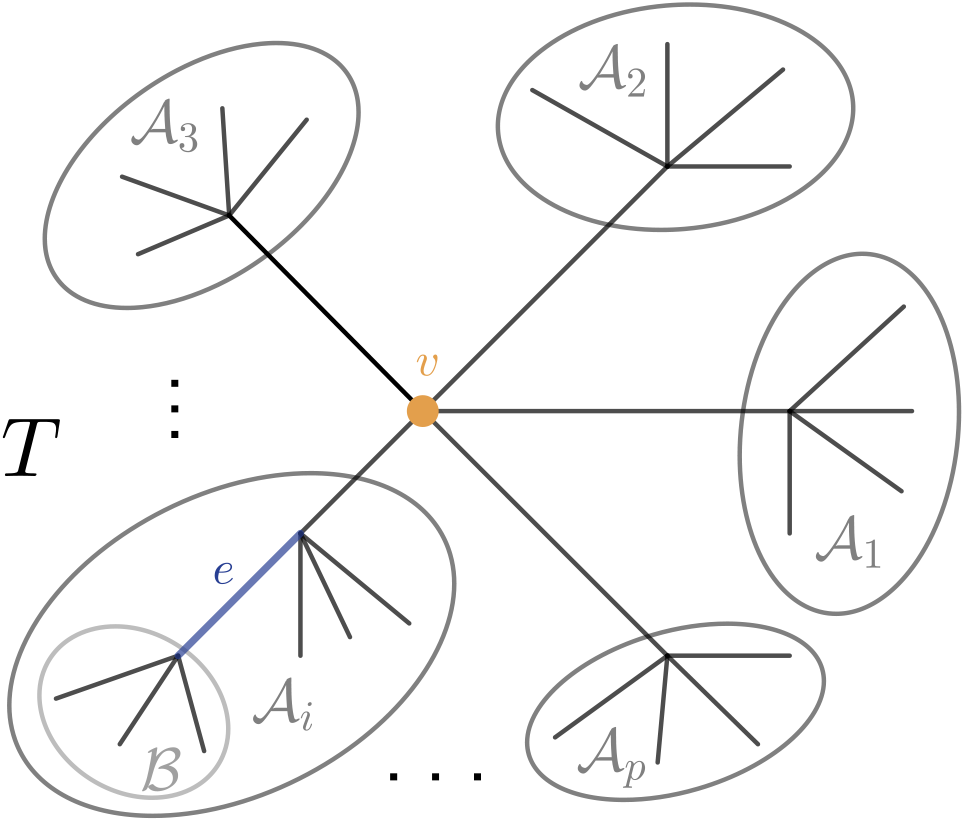
Unrooted tree *T* on leaf set *X* with *X* = *A*_1_ ∪ … ∪ *A*_*p*_ as constructed in the proof of Lemma 2.3. Vertex *v* realizes the blob partition *ψ* = *A*_1_ | · · · | *A*_*p*_ and edge *e* realizes the split *σ* = *B* | *C*, where *B* ⊊ *A*_*i*_.

The resulting tree simultaneously realizes both *ψ* and *σ*, showing that they are compatible.

The following lemma gives a criterion for when two blob partitions of *X* are compatible. In contrast to the previous lemma, the stated inclusions need not be strict. In particular, when two blob vertices are adjacent (i.e., connected by an edge), equality occurs.

#### Lemma 2.4.

*Let ψ*_1_ = *A*_1_ | *A*_2_ | … | *A*_*p*_ *and ψ*_2_ = *B*_1_ | *B*_2_ | … | *B*_*q*_ *be two distinct blob partitions of X with p, q* ≥ 3. *Then, ψ*_1_ *and ψ*_2_ *are compatible if and only if there exists a block A*_*i*_ *of ψ*_1_ *and a block B*_*j*_ *of ψ*_2_ *such that*

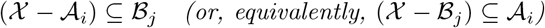

*Proof*. Let *ψ*_1_ = *A*_1_ | *A*_2_ | … | *A*_*p*_ and *ψ*_2_ = *B*_1_ | *B*_2_ | … | *B*_*q*_ be two distinct partitions of *X*, with *p, q* ≥ 3. Assume first that *ψ*_1_ and *ψ*_2_ are compatible. By definition, there exists an unrooted tree *T* with leaf set *X* and with distinct inner vertices *s* and *t* such that *s* induces *ψ*_1_ and *t* induces *ψ*_2_ (Fig. S3). Removing *s* from *T* disconnects the tree into components whose leaf sets are exactly *A*_1_, …, *A*_*p*_; similarly, removing *t* disconnects *T* into components with leaf sets *B*_1_, …, *B*_*q*_.

Let *A*_*i*_ be the block of *ψ*_1_ corresponding to the component of *T* − {*s*} that contains *t*, and let *B*_*j*_ be the block of *ψ*_2_ corresponding to the component of *T* − {*t*} that contains *s*.

Take any *x* ∈ *X* − *A*_*i*_. Since *x* lies in a component of *T* − {*s*} distinct from the one containing *t*, the unique path in *T* from *x* to *t* passes through *s*. Consequently, this path reaches *s* before *t*, so *x* lies in the component of *T* − {*t*} that contains *s*, whose leaf set is *B*_*j*_. Hence,

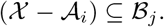

**Figure S3:**
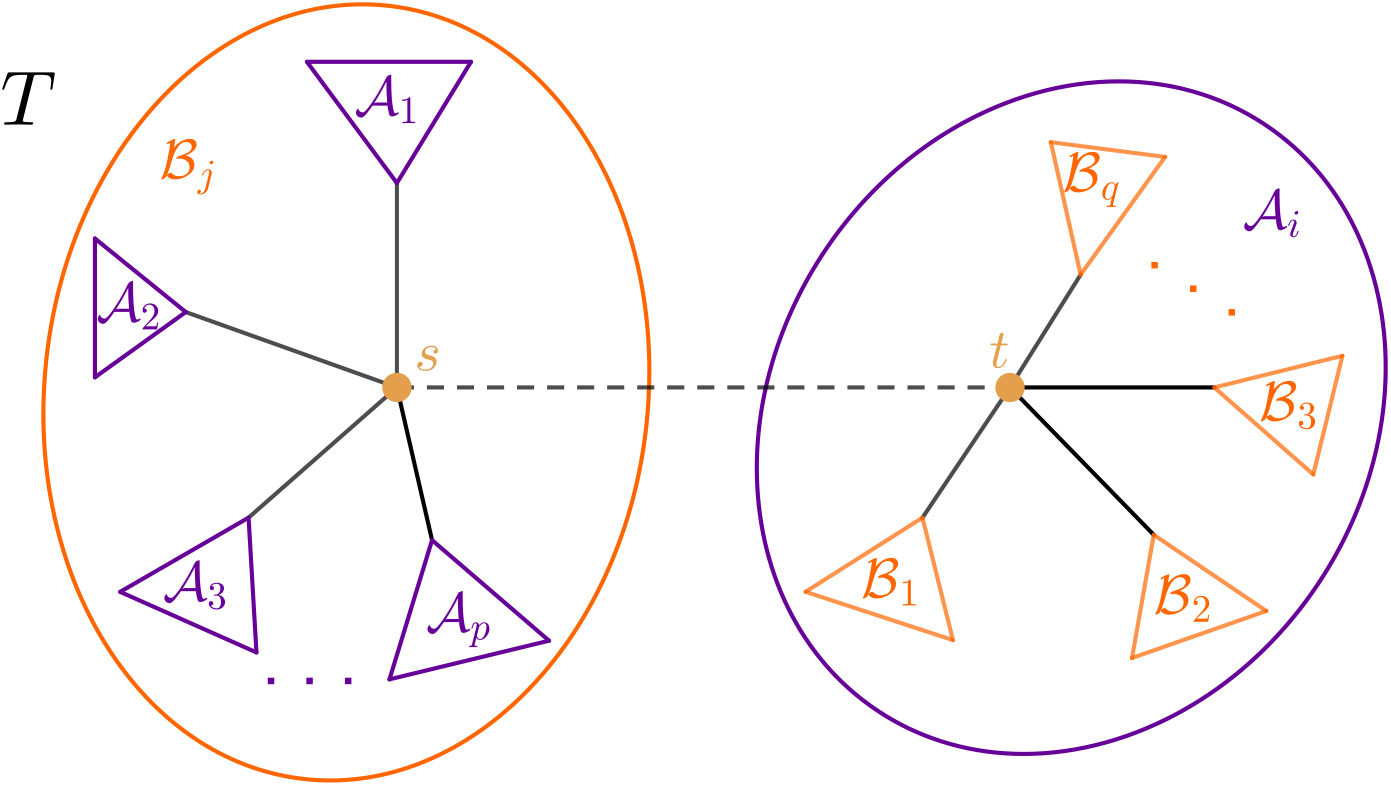
An unrooted tree *T* on leaf set *X* that realizes the two compatible blob partitions *ψ*_1_ = *A*_1_ | *A*_2_ |… | *A*_*p*_ and *ψ*_2_ = *B*_1_ | *B*_2_ | … | *B*_*q*_, as described in the first part of the proof of Lemma 2.4.

Now, suppose *ψ*_1_ = *A*_1_ | *A*_2_ | … | *A*_*p*_ and *ψ*_2_ = *B*_1_ | *B*_2_ | … | *B*_*q*_ are two distinct partitions of *X*, and assume there exist indices *i* ∈ {1, …, *p*} and *j* ∈ {1, …, *q*} such that

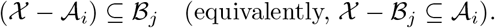

This condition implies that the two blocks *A*_*i*_ and *B*_*j*_ together cover all of *X*, i.e.,

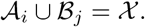

Equivalently, each element of *X* either lies in *A*_*i*_, in *B*_*j*_, or in their intersection *A*_*i*_ ∩ *B*_*j*_.

We now construct an unrooted tree *T* on leaf set *X* with two distinct inner vertices *s* and *t* such that *s* induces *ψ*_1_ and *t* induces *ψ*_2_. Let *C* := *A*_*i*_ ∩ *B*_*j*_.

- *Case 1: C* = ∅. In this case, *A*_*i*_ and *B*_*j*_ form an *X*-split, since *A*_*i*_ ∪ *B*_*j*_ = *X* and *A*_*i*_ ∩ *B*_*j*_ = ∅. In particular, *X* − *A*_*i*_ = *B*_*j*_ and *X* − *B*_*j*_ = *A*_*i*_. We construct *T* as follows (see Fig. S4(i)):
  1. Start with two vertices *s* and *t* joined by an edge.
  2. For each block *A*_*k*_ of *ψ*_1_ with *k* ≠ *i*, attach to *s* a subtree whose leaf set is exactly *A*_*k*_.
  3. For each block *B*_*l*_ of *ψ*_2_ with *l* ≠ *j*, attach to *t* a subtree whose leaf set is exactly *B*_*l*_. By construction, removing *s* yields connected components with leaf sets *A*_1_, …, *A*_*p*_, and removing *t* yields connected components with leaf sets *B*_1_, …, *B*_*q*_. Hence *s* induces *ψ*_1_ and *t* induces *ψ*_2_.
- *Case 2: C* ≠ ∅. In this case, the blocks *A*_*i*_ and *B*_*j*_ overlap. We construct *T* as follows (see Fig. S4(ii)):
  1. Start with three vertices *s, w, t* arranged in a path, introducing the edges {*s, w*} and {*w, t*}.
  2. For each block *A*_*k*_ of *ψ*_1_ with *k* ≠ *i*, attach to *s* a subtree whose leaf set is exactly *A*_*k*_.
  3. For each block *B*_*l*_ of *ψ*_2_ with *l* ≠ *j*, attach to *t* a subtree whose leaf set is exactly *B*_*l*_.
  4. Attach to *w* a subtree whose leaf set is exactly *C* = *A*_*i*_ ∩ *B*_*j*_.

Removing *s* yields connected components with leaf sets *A*_*k*_ for *k* ≠ *i*, together with a single component containing *w, t* and all leaves in *A*_*i*_; thus *s* induces *ψ*_1_. Similarly, removing *t* yields connected components corresponding exactly to the blocks of *ψ*_2_, so *t* induces *ψ*_2_.

**Figure S4:**
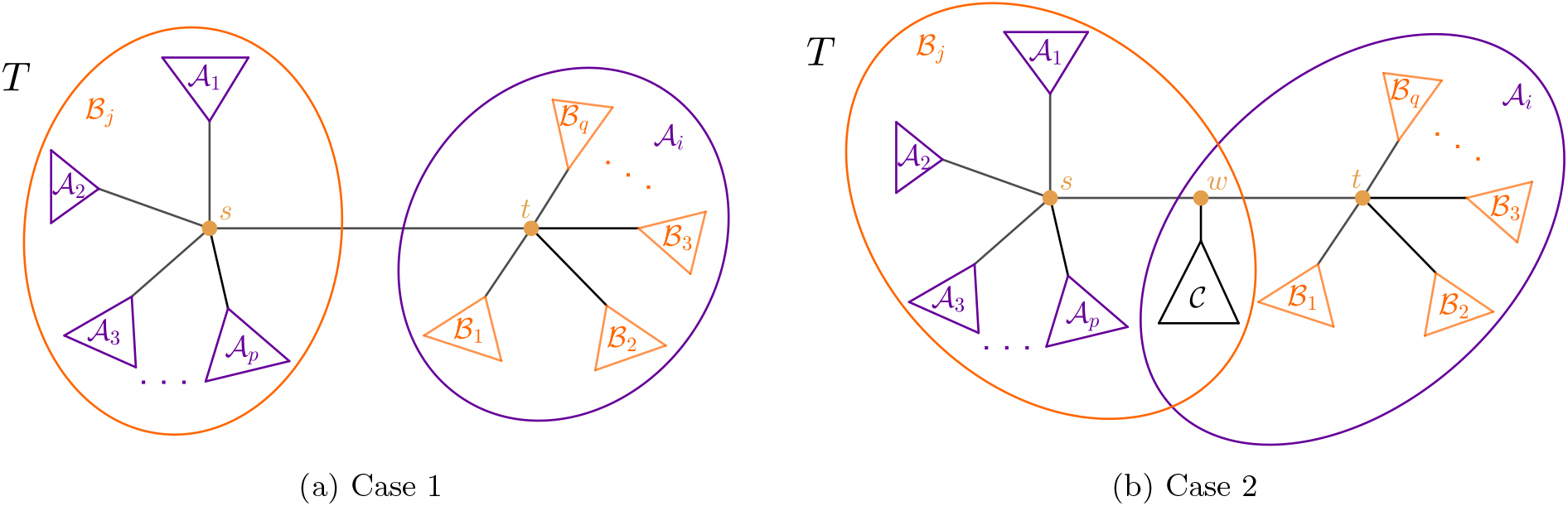
Unrooted trees *T* on leaf set *X*, constructed in the second part of the proof of Lemma 2.4. Panel (i) illustrates Case 1, in which the blocks *A*_*i*_ and *B*_*j*_ are disjoint, while Panel (ii) illustrates Case 2, in which they overlap.

### 2.2 Pairwise compatibility implies global compatibility

In the previous section, we derived simple conditions for two blob partitions, or for a blob partition and a non-redundant split, to be compatible, and we recall that for two splits, blob compatibility coincides with the usual notion of tree-compatibility for *X*-splits. Clearly, if a collection of blob partitions and non-redundant splits is globally compatible, then every pair of its elements is realized by the same tree of blobs and hence satisfies the corresponding pairwise compatibility conditions. In the following, we show that global compatibility is completely characterized by these pairwise compatibility conditions.

#### Theorem 2.5.

*Let* Ψ* = {*ψ*_1_, …, *ψ*_*m*_} *be a collection of blob partitions of X, each with at least three blocks, and let* Σ* *be a collection of X-splits that are non-redundant with respect to* Ψ*. *Then the pair* (Ψ*, Σ*) *is compatible if and only if the following conditions hold:*

1. *every pair ψ*_*i*_, *ψ*_*j*_ ∈ Ψ* *is compatible;*
2. *every pair σ, σ*′ ∈ Σ* *is tree-compatible in the usual sense of X-splits (and hence blob-compatible);*
3. *every pair ψ* ∈ Ψ* *and σ* ∈ Σ* *is compatible*.

The main idea of the proof of Theorem 2.5 is to reduce global (blob-)compatibility to a question about tree-compatibility of splits. Each blob partition can be viewed as inducing a collection of splits corresponding to its blocks. Using the assumed pairwise compatibility, we show that these induced splits are all pairwise tree-compatible (Lemma 2.6), and that they are also tree-compatible with the non-redundant splits under consideration (Lemma 2.7). As a consequence, all these splits together determine a unique unrooted tree on leaf set *X*. We then show that this tree realizes the given blob partitions and non-redundant splits, thereby establishing global compatibility.

We first relate the blob partitions to splits.

#### Lemma 2.6.

*Let* Ψ* *be a collection of blob partitions of X. For each partition ψ* = *A*_1_ | · · · | *A*_*p*_ ∈ Ψ* *(with p* ≥ 3*), define*

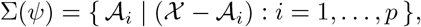

*and let*

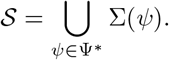

*If every pair of partitions in* Ψ* *is compatible, then every pair of splits in S is compatible in the usual sense of X-splits*.

*Proof*. Take two splits in *S*.

1. *Case 1: Both splits come from the same partition ψ*. Let *ψ* = *A*_1_ | · · · | *A*_*p*_ ∈ Ψ*, and suppose the two splits are *A*_*i*_ | (*X* − *A*_*i*_) and *A*_*j*_ | (*X* − *A*_*j*_) with *i* ≠ *j*. Since the subsets of a partition are disjoint, we have *A*_*i*_ ∩ *A*_*j*_ = ∅. Hence the two splits are tree-compatible.
2. *Case 2: The splits come from distinct partitions ψ*_*s*_ *and ψ*_*t*_. Let

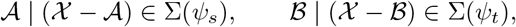

where *A* is a block of *ψ*_*s*_ and *B* a block of *ψ*_*t*_.

Since *ψ*_*s*_ and *ψ*_*t*_ are compatible, by Lemma 2.4, there exist blocks *A*_*i*_ ∈ *ψ*_*s*_ and *B*_*j*_ ∈ *ψ*_*t*_ such that (*X* − *A*_*i*_) ⊆ *B*_*j*_; hence the splits *A*_*i*_ | (*X* − *A*_*i*_) and *B*_*j*_ | (*X* − *B*_*j*_) are tree-compatible.

Thus, if *A* = *A*_*i*_ and *B* = *B*_*j*_, then *A* | (*X* − *A*) and *B* | (*X* − *B*) are tree-compatible. We now show that all other choices for *A* and *B* also yield tree-compatible splits.

- If *A* ⊊ (*X* − *A*_*i*_) ⊆ *B*_*j*_ and *B* = *B*_*j*_, then *A* ⊊ *B*, and the splits *A* | (*X* − *A*) and *B* | (*X* − *B*) are tree-compatible.
- Similarly, if *A* = *A*_*i*_ and *B* ⊊ (*X* − *B*_*j*_) ⊆ *A*_*i*_, then *B* ⊊ *A*, and the splits *A* | (*X* − *A*) and *B* | (*X* − *B*) are tree-compatible.
- Finally, if *A* ⊊ (*X* −*A*_*i*_) ⊆ *B*_*j*_ and *B* ⊊ (*X* −*B*_*j*_) ⊆ *A*_*i*_, then *A*∩*B* = ∅, and the splits *A* | (*X* −*A*) and *B* | (*X* − *B*) are tree-compatible.

Since the choice of splits was arbitrary, every pair of splits in *S* is tree-compatible.

We next show that if a blob partition *ψ* and a non-redundant split *σ* are compatible, then all splits induced by *ψ* (as described in the previous lemma) and *σ* are compatible in the usual sense of *X*-splits.

#### Lemma 2.7.

*Let ψ* = *A*_1_ | · · · |*A*_*p*_ *be a blob partition of X with p* ≥ 3, *and let σ* = *B* | *C be an X-split. If ψ and σ are compatible, then* every split induced by *ψ* is tree-compatible with *σ. That is*,

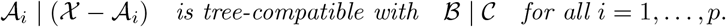

*Proof*. Assume that *ψ* = *A*_1_ | · · · | *A*_*p*_ and *σ* = *B* | *C* are compatible. By Lemma 2.3, there exists a block *A*_*i*_ of *ψ* such that either *B* ⊆ *A*_*i*_ or *C* ⊆ *A*_*i*_. Without loss of generality, assume *B* ⊆ *A*_*i*_.

Consider first the split *A*_*i*_ | (*X* −*A*_*i*_). Since *B* ⊆ *A*_*i*_, the splits *A*_*i*_ | (*X* −*A*_*i*_) and *B* | *C* are tree-compatible. Now, let *A*_*j*_ | (*X* − *A*_*j*_) be any other split of *ψ* with *j* ≠ *i*. Since blocks of a partition are disjoint, *A*_*j*_ ∩ *A*_*i*_ = ∅, hence *A*_*j*_ ∩ *B* = ∅. Therefore, *A*_*j*_ | (*X* − *A*_*j*_) is also tree-compatible with *B* | *C*. Since *j* was arbitrary, all splits induced by *ψ* are tree-compatible with *σ*.

#### Remark 2.8.

A non-redundant split is tree-compatible with all blob-induced splits, yet it is distinct from each of them. Furthermore, compatibility between two blob partitions, or between a blob partition and a non-redundant split, implies tree-compatibility of the corresponding blob induced splits (as in Lemma 2.6) in the usual sense of *X*-splits. However, these two notions are not equivalent: for example, the split {*A, B*} | {*C, D*} is compatible (in the usual sense) with all splits induced by the partition {*A*} | {*B*} | {*C*} | {*D*}, i.e., with the splits {*A*} | *X* − {*A*}, {*B*} | *X* − {*B*}, {*C*} | *X* − {*C*}, and {*D*} | *X* − {*D*}, but the split and partition are not compatible as the conditions of Lemma 2.3 are violated.

*Proof of Theorem 2*.*5*. The forward implication has already been noted above, so it suffices to prove the converse.

For the converse, assume that every pair of partitions in Ψ* is compatible, every pair of splits in Σ* is tree-compatible (and hence (blob-)compatible), and every partition-split pair is compatible.

Consider the collection of blob-induced splits (as in Lemma 2.6) and non-redundant splits, i.e., let

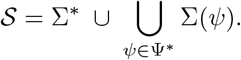

By Lemma 2.6, all splits induced by the partitions in Ψ* are pairwise tree-compatible. Moreover, by Lemma 2.7, each split in Σ(*ψ*) is tree-compatible with every split in Σ*, because each partition-split pair is compatible. Hence, every pair of splits in *S* is compatible in the usual sense of *X*-splits.

By the Splits-Equivalence Theorem (Theorem 1.5), there exists a unique unrooted phylogenetic *X* –tree *T* whose set of splits is exactly *S* (together with all trivial splits, if these are not already contained in *S*).

Fix a partition *ψ* ∈ Ψ* with blocks *A*_1_, …, *A*_*p*_. For each *i*, the split *A*_*i*_ | (*X* − *A*_*i*_) belongs to *S*. Since the sets *A*_1_, …, *A*_*p*_ form a partition of *X*, in the tree *T*, the edges corresponding to the splits in Σ(*ψ*) are all incident to a single interior vertex *v*. The vertex *v* has *p* incident subtrees whose leaf sets are exactly *A*_1_, …, *A*_*p*_, and hence *v* realizes the blob partition *ψ*.

Moreover, distinct partitions in Ψ* are realized by distinct interior vertices of *T*, and every split in Σ* is realized by an edge of *T*. Therefore, the tree *T* simultaneously realizes all blob partitions in Ψ* and all non-redundant splits in Σ*. It follows that the pair (Ψ*, Σ*) is compatible. By the uniqueness guaranteed by the Splits-Equivalence Theorem, this realization of (Ψ*, Σ*) by *T* is itself unique.

### 2.3 An algorithm to determine pairwise compatibility

We end this section by introducing an algorithm to determine whether two partitions are compatible. This algorithm uses a vectorial encoding of splits and partitions. We also use this encoding to check whether two partitions are equivalent in *O*(*n*). To do so, we encode splits and partitions as vectors with one entry per taxon, and one number per block. For instance, *A, B, C*, and *D* correspond to entries in positions 1, 2, 3, and 4, respectively; and the vector 0012333 … 3 corresponds to the partition {*A, B*} | {*C*} |{*D*} | (*X* − {*A, B, C, D*}). Ordering the numbers representing the blocks (as in the previous example) allows us to check the equivalence of two partitions in linear time.

Recall from Section 2.1 that two partitions *ψ*_1_ = *A*_1_ | *A*_2_ | · · · | *A*_*p*_ and *ψ*_2_ = *B*_1_ | *B*_2_ | · · · | *B*_*q*_, with *p, q ≥* 2, are compatible if all but one block *A*_*i*_ are contained in a single block *B*_*j*_. Algorithm 1 exploits this characterization to determine pairwise compatibility between two partitions, whether induced by two splits, two blob vertices, or by a split and a blob vertex. An illustration of the algorithm is provided in Figure S5.

#### Algorithm 1

Algorithm to determine compatibility between two partitions set of size *n*.

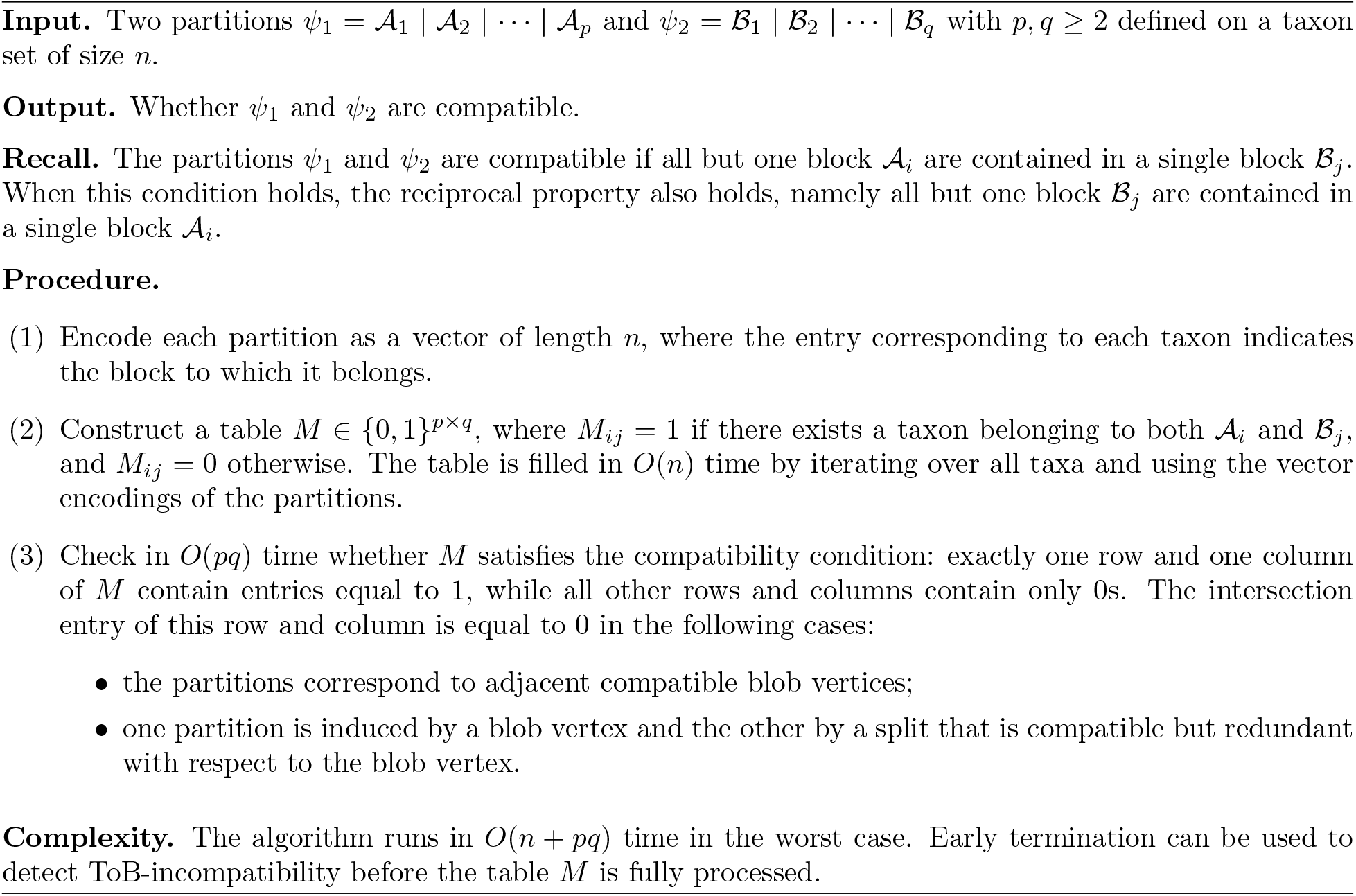

**Figure S5:**
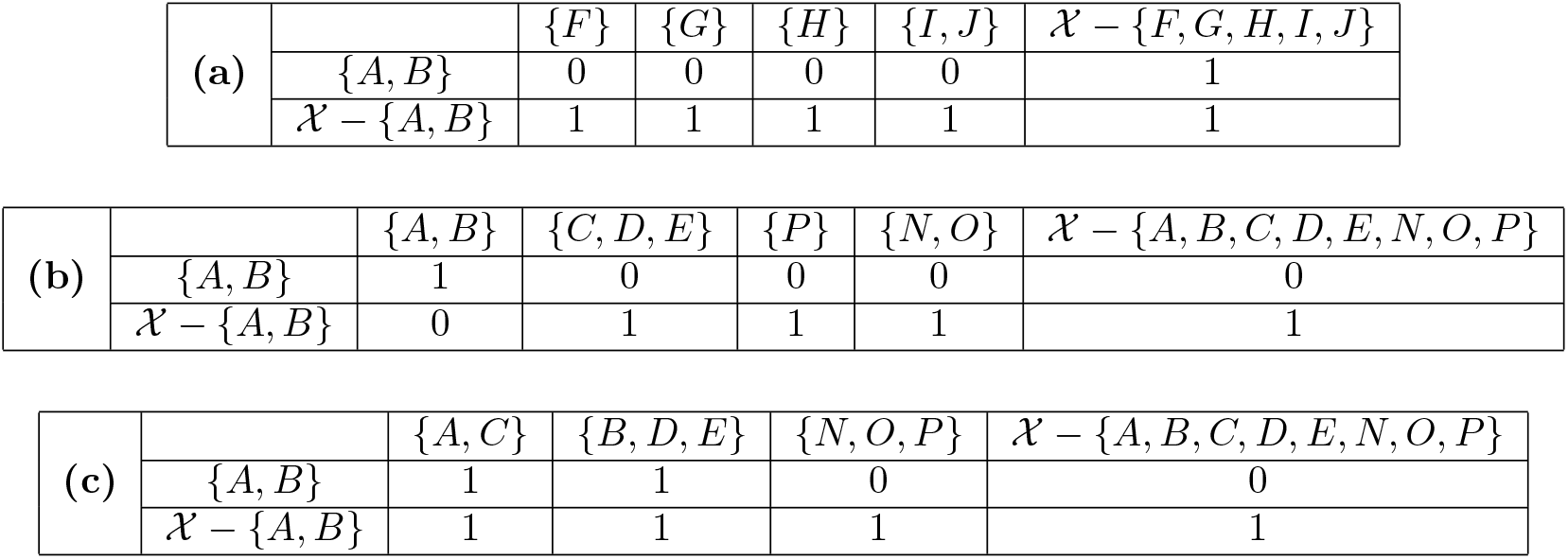
Computing the compatibility condition for *X* = {*A, B, C, D, E, F, G, H, I, J, K, L, M, N, O, P*}: (a) One split and one blob partition, which are compatible. (b) One split and one blob partition, which are compatible, but the split is redundant with the blob. (c) One split and one blob partition that are not compatible.

## 3 Consensus methods for trees of blobs

### 3.1 Methods and algorithms

We now formally introduce the consensus methods for trees of blobs as described in the main part of our manuscript. Given a collection of trees of blobs on the same taxon set, we extract from each tree its blob partitions and its non-redundant splits. We then record how frequently each such partition component appears across the input trees. These partition components are subsequently filtered and combined according to their frequency and mutual compatibility, yielding different types of consensus trees of blobs.

We now formalize this procedure and define three consensus methods for trees of blobs, analogous to the classical strict, majority-rule, and greedy (or extended majority-rule) consensus trees.

#### Definition 3.1

(Consensus component sets). Let *T* = {*T*_1_, …, *T*_*k*_} be a collection of trees of blobs on a common taxon set *X*. For each *T*_*i*_, let Ψ(*T*_*i*_) denote its set of blob partitions and Σ(*T*_*i*_) its set of non-redundant splits. Let

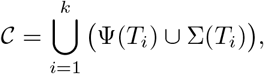

be the multiset consisting of all elements of Ψ(*T*_*i*_) ∪ Σ(*T*_*i*_) for *i* = 1, …, *k*, where each element (either a blob partition or a non-redundant split) is counted according to its frequency in the input trees.

We define the strict, majority-rule, and greedy consensus *component sets* as collections of blob partitions and non-redundant splits selected according to their frequency and compatibility across *T*.

1. **Strict consensus:** the set of blob partitions and non-redundant splits that occur in every tree in *T*.
2. **Majority-rule consensus:** the set of blob partitions and non-redundant splits that occur in strictly more than half of the trees in *T*.
3. **Greedy consensus:** the set obtained by iteratively (greedily) adding blob partitions and non-redundant splits from *C* in decreasing order of frequency, provided each added element is compatible with all previously selected elements. In the case of a tie between a blob partition and a non-redundant split, we select the blob partition, since it is a more complex structure.

In the greedy consensus construction, it suffices to test whether a candidate partition or split is pairwise compatible with all previously accepted elements. If so, it is guaranteed to be compatible with the entire set and can be safely added.

#### Proposition 3.2.

*Let* Ψ* *be a set of blob partitions of X and* Σ* *a set of non-redundant X-splits. Assume that* (Ψ*, Σ*) *is globally compatible. Let C be a blob partition or split such that C is compatible with every element of* Ψ* ∪ Σ*. *Then* (Ψ* ∪ {*C*}, Σ*) *or* (Ψ*, Σ* ∪ {*C*}) *is globally compatible*.

*Proof*. By Theorem 2.5, global compatibility is equivalent to pairwise compatibility. Since *C* is pairwise compatible with all existing elements, the enlarged collection remains pairwise compatible, and hence mutually compatible.

We now show that each consensus method produces a biologically meaningful summary that can be interpreted as a tree of blobs, rather than merely as a collection of abstract partitions and splits.

#### Proposition 3.3.

*Each consensus set contains pairwise compatible components, and therefore can be realized by an unrooted tree of blobs*.

*Proof*. We verify the three compatibility conditions of Proposition 2.5.

- **Strict and majority-rule consensus**. By construction, every pair of partition components (whether partition–partition, split–split, or partition–split) contained in these consensus sets co-occurs in at least one input tree of blobs. Consequently,
  - any two blob partitions are compatible;
  - any two non-redundant splits are compatible;
  - any blob partition and split are compatible. Notice that a split is included in the majority-rule (resp. strict) consensus component set only if it is non-redundant in a strict majority of (resp. all) input trees of blobs. Therefore, it is non-redundant with respect to every blob partition included in the corresponding component set, even though (in the case of majority-rule) it may be redundant in other input trees of blobs.
  Thus, the conditions of Proposition 2.5 are satisfied.
- **Greedy consensus**. At each step of the greedy procedure, a partition or split is added only if it is compatible with all previously selected elements. Moreover, a split is added only if it carries information not already represented, that is, only if it is non-redundant with respect to all existing blob partitions. Consequently, all pairs of elements in the final set are pairwise compatible (with splits being non-redundant), and Proposition 2.5 applies.

In all three cases, the resulting set of blob partitions and non-redundant splits is compatible and therefore realizable by an unrooted tree of blobs.

#### Remark 3.4.

Notice that a consensus tree of blobs may contain vertices of degree four or higher that do not correspond to any blob vertex. For instance, in Figure S6, the majority-rule consensus tree of blobs includes an interior vertex of degree 6 that is not a blob vertex, as it does not appear as a blob vertex in a strict majority of the input trees (in fact, it is absent from all of them). We therefore explicitly label inner vertices that correspond to blob vertices in the input trees as blob vertices in the consensus tree of blobs.

Next, we establish uniqueness of the consensus tree of blobs.

#### Corollary 3.5.

*Let* (Ψ*, Σ*) *be any of the consensus component sets obtained by the strict or majority-rule procedure. Then there exists a unique unrooted tree of blobs that realizes the blob partitions in* Ψ* *and the non-redundant splits in* Σ* *(as well as all trivial splits)*.

*For the greedy consensus method, every possible greedy ordering of adding partition components yields a compatible set, and each such set is realized by a unique tree of blobs; however, different orderings (e*.*g*., *due to equally frequent partition components) may result in distinct consensus trees of blobs*.

Non-uniqueness of the greedy consensus tree arises only when multiple blob partitions or splits occur with identical frequencies. In this case, alternative consensus trees of blobs may exist. In practical implementations, certain ties can be resolved, for example by prioritizing blob partitions over splits as implemented in our software. However, certain ties are inevitable and the final solution of any greedy approach can be suboptimal in terms of global support.

*Proof*. By Theorem 3.3, the pair (Ψ*, Σ*) is compatible. Let

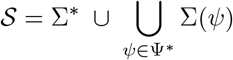

be the associated set of splits.

By Theorem 2.5, all splits in *S* are pairwise tree-compatible. By the Splits-Equivalence Theorem, there exists a unique unrooted phylogenetic *X*-tree *T* whose edge-induced splits are exactly *S* (together with all trivial splits, if these are not already contained in).

For each *ψ* ∈ Ψ*, the splits in Σ(*ψ*) are realized by edges incident to a unique interior vertex of *T*, which therefore induces *ψ*. Hence *T* realizes exactly the blob partitions in Ψ* and the non-redundant splits in Σ* (again, together with all trivial splits).

Uniqueness of *T* follows from uniqueness of the split realization.

For the greedy consensus method, different orderings of equally frequent partition components may result in different split sets *S* and hence in distinct consensus trees of blobs.

#### Algorithm: Early stopping of the greedy selection of components

We describe here a criterion to speed up the iteration through the lists of blob partitions and non-redundant splits, when adding components to build a consensus tree of blobs.

Once the current tree of blobs has no divergence node of degree greater than 3, then no new component can further resolve it. Only blob partitions with 3 blocks may be added to the list of retained components, which would add a blob label to an existing node in the current tree of blobs. If input networks do not have degree-3 blobs, then the process can terminate. Otherwise, the iteration process should continue, but non-redundant splits and blob partition with 4 or more blocks can be ignored.

To determine if the current tree of blobs has no divergence node of degree 4 or higher, one may simply compare its number of internal edges with the maximum possible. Each blob node retained in the tree of blobs, from a blob partition *ψ* with *p*(*ψ*) blocks, decreases the maximum number of internal edges by *p*(*ψ*) − 3. Therefore, the maximum number of edges in the consensus tree of blobs, given the blob partitions already retained, is *n* − 3 − ∑_*ψ*∈Ψ retained_(*p*(*ψ*) − 3).

### 3.2 Properties of the consensus tree of blobs

We now establish several properties for the tree of blobs consensus methods described above. We first analyze how the consensus behaves when there is strong agreement among the input trees of blobs, beginning with the case in which one tree of blobs occurs with strict majority frequency.

#### Proposition 3.6.

*Let T* = {*T*_1_, …, *T*_*k*_}*be a collection of trees of blobs on a common taxon set X. If a strict majority of the input trees are identical to some tree of blobs T, then the majority-rule consensus tree of blobs equals T*.

*The same holds for the greedy consensus tree of blobs when all vertices of degree at least four in the input trees of blobs are blob vertices (and not divergence vertices)*.

*Proof*. First, consider the majority-rule consensus tree of blobs. Every blob partition and every non-redundant split of *T* appears in strictly more than *k/*2 input trees, hence is contained in the majority-rule component set. Any component not present in *T* appears in at most *k/*2 trees and is therefore excluded. Thus the selected component set is exactly Ψ(*T*) ∪ Σ(*T*), which uniquely determines *T*.

Now, for the greedy consensus tree of blobs, notice that when all vertices of degree at least four in the input trees of blobs are blob vertices, the component set Ψ(*T*) ∪ Σ(*T*) is already maximal with respect to compatibility. Hence, no additional minority component can be added during the greedy phase, and the greedy consensus tree of blobs also coincides with *T*.

Complete agreement between the input trees of blobs is obtained as a special case of the situation above, yielding a property known as *idempotence* or *unanimity* in the literature [2].

#### Corollary 3.7.

*Let T* = {*T*_1_, …, *T*_*k*_} *be a collection of trees of blobs on a common taxon set X*. *If all input trees of blobs are identical, i*.*e. T*_1_ = · · · = *T*_*k*_ = *T*, *then the strict, majority-rule, and greedy consensus trees of blobs all equal T*.

The next two properties relate to the majority-rule consensus tree of blobs. First, recall that an unrooted phylogenetic *X*-tree *T* ′ *refines* an unrooted phylogenetic *X*-tree *T* if every split of *T* is a split of *T* ′. As indicated in the main part of the paper, the standard majority-rule consensus tree (with respect to splits) refines the majority-rule consensus tree of blobs. For an example, see Figure S6.

#### Proposition 3.8.

*Let T* = {*T*_1_, …, *T*_*k*_} *be a collection of trees of blobs on a common taxon set X. Then every split induced by the majority-rule consensus tree of blobs is also induced by the standard majority-rule consensus tree. In particular, the standard majority-rule consensus tree refines the majority-rule consensus tree of blobs*.

*Proof*. Let *T* = {*T*_1_, …, *T*_*k*_} be a collection of trees of blobs on a common taxon set *X*. Let *T*_blob_ denote the majority-rule consensus tree of blobs (built from blob partitions and non-redundant splits that occur in strictly more than *k/*2 trees) and let *T*_split_ be the standard majority-rule consensus tree (built from splits that occur in strictly more than *k/*2 trees).

**Figure S6:**
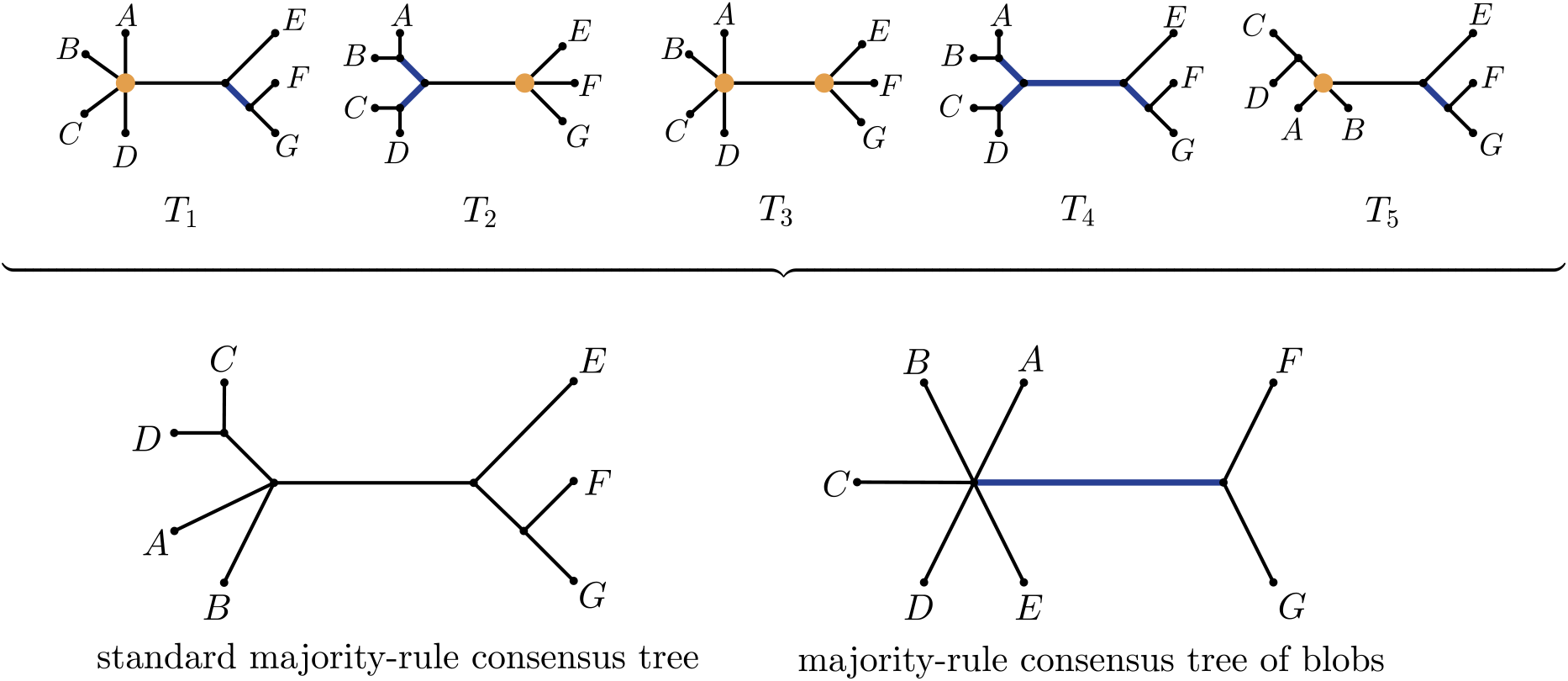
Five trees of blobs *T*_1_, …, *T*_5_ on common taxon set *X* = {*A*, …, *G*} (where all vertices of degree at least four are interpreted as blob vertices and blue edges correspond to non-trivial non-redundant splits) together with the standard majority-rule consensus tree (based solely on frequency of splits) and our majority-rule consensus tree of blobs (based on the frequency of blob partitions and non-redundant splits). Notice that the standard majority-rule consensus tree (with respect to splits) refines the majority-rule consensus tree of blobs. Furthermore, notice that vertex of degree 6 in the majority-rule consensus tree of blobs is not interpreted as a blob vertex (as none of the input trees contain this blob vertex), but is a divergence vertex.

We claim that every split displayed by *T*_blob_ is also displayed by *T*_split_.

Notice that any split *σ* = *A* | *B* displayed by *T*_blob_ arises in one of following two ways:

1. The split *σ* is induced by a blob partition *ψ* = *A*_1_ | … | *A*_*p*_ that occur in strictly more than *k/*2 trees.
2. The split *σ* is a non-redundant split that occurs in strictly more than *k/*2 trees. Thus, it suffices to show that both types are majority splits in the usual sense.

First, let *ψ* = *A*_1_ | … | *A*_*p*_ be a blob partition appearing in strictly more than *k/*2 trees. Fix any induced split *σ* = *A*_*i*_ | (*X* −*A*_*i*_). Now, notice that every tree that induces *ψ* necessarily displays all splits induced by *ψ*. Hence,

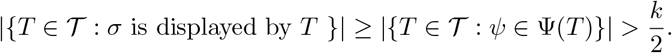

So, *σ* is a split that occurs in strictly more than *k/*2 trees, and therefore is displayed by *T*_split_.

Similarly, if *σ* is a non-redundant split that occurs in strictly more than *k/*2 trees, it is clearly also displayed by *T*_split_.

In summary, all splits displayed by *T*_blob_ are displayed by *T*_split_, which shows that the standard majority-rule consensus tree refines the majority-rule consensus tree of blobs.

#### Remark 3.9.

Proposition 3.8 reflects an important difference in how support is measured in the two consensus constructions. In the standard majority-rule consensus, every split is counted independently across the input trees. In contrast, the majority-rule consensus tree of blobs focuses on blob partitions and only then considers *non-redundant* splits relative to those partitions.

As a consequence, many splits that are counted separately in the standard majority analysis are treated as redundant with respect to a blob partition, and thus do not accumulate frequency in the tree of blobs setting. Their effective frequency is therefore lower in the majority-rule consensus tree of blobs construction than in the standard one. This leads to fewer retained splits overall, so the majority-rule consensus tree of blobs is typically less resolved.

This loss of resolution is intentional: the majority-rule consensus tree of blobs preserves blob vertices as important features rather than over resolving relationships within them. Since blob vertices represent regions of reticulate evolution, the tree of blobs consensus preserves these structures while avoiding unsupported refinement.

Finally, we establish a median property of the majority rule consensus tree, analogous to the well-known median property of the standard majority-rule consensus tree [1]. Recall that each tree of blobs *T* is represented by the component set

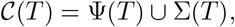

where Ψ(*T*) and Σ(*T*) are the blob partitions and non-redundant splits of *T*, respectively.

Given two trees of blobs *T*_1_, *T*_2_ on a common taxon set *X*, we define a distance

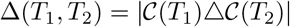

between *T*_1_ and *T*_2_ based on the symmetric difference of their respective component sets.

Now, given a collection *T* = {*T*_1_, …, *T*_*k*_} of trees of blobs on a common taxon set *X*, a *median tree T* * minimizes

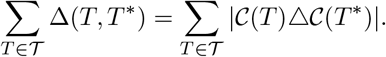

#### Proposition 3.10.

*Let T* = {*T*_1_, …, *T*_*k*_} *be a collection of trees of blobs on a common taxon set*. *Under the symmetric-difference distance on blob partitions and non-redundant splits, the majority-rule consensus tree of blobs is a median of T*.

*Proof*. Let *T* be any tree of blobs on *X*. The distance Δ(*T, T*_*i*_) counts the number of blob partitions and non-redundant splits which occur in precisely one of *T* and *T*_*i*_ (but not in both).

Fix a blob partition *ψ*, and let *f* (*ψ*) denote the number of trees in *T* that contain *ψ*. If *ψ* is contained in *T*, then it contributes *k* − *f* (*ψ*) to the total distance 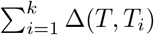 if *ψ* is not contained in *T*, then it contributes *f* (*ψ*). Hence, the contribution of *ψ* is minimized by including *ψ* when 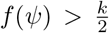, and by excluding *ψ* when 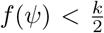. If 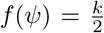, both choices give the same contribution, so either inclusion or exclusion is distance-minimizing.

Since each blob partition and each non-redundant split contributes independently to the total distance, the sum 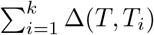 is minimized by any tree *T* that contains every blob partition and non-redundant split occurring in strictly more than half of the trees, excludes those occurring in strictly fewer than half, and makes an arbitrary choice for those occurring in exactly half of the trees.

By definition, the majority-rule consensus tree of blobs is therefore one such median tree of *T*.

#### Remark 3.11.

In the argument above, if 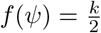 (which can occur only when *k* is even), then including or excluding *ψ* yields the same contribution 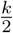 to the total distance. Hence, such blob partitions or non-redundant splits do not affect the value of and either choice is distance-minimizing.

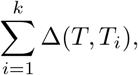

and either choice is distance-minimizing.

It follows that the median need not be unique: any tree obtained by freely adding or omitting elements that occur in exactly half of the input trees is also a median (provided that these elements are mutually compatible). The majority-rule consensus tree of blobs corresponds to the conventional choice of retaining only those elements whose frequency is strictly greater than ^*k*^, thereby selecting a canonical representative among the (possibly many) medians.

#### Non-uniqueness of greedy consensus methods and other observations

We illustrate now that a greedy consensus ToB need not be unique, and that it can differ from a traditional majority-rule consensus tree and also from the input trees. To this end, we modify Example 2.1 in the main text taking as input the trees of blobs shown in Figure S7.

**Figure S7:**
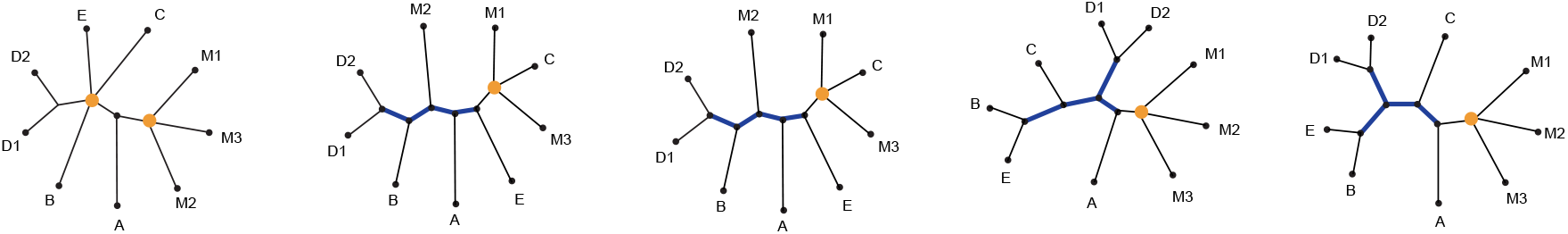
This input has 3 greedy consensus ToBs and 3 traditional greedy consensus trees.

Components and their frequencies are shown in the next table, ordered by frequency.

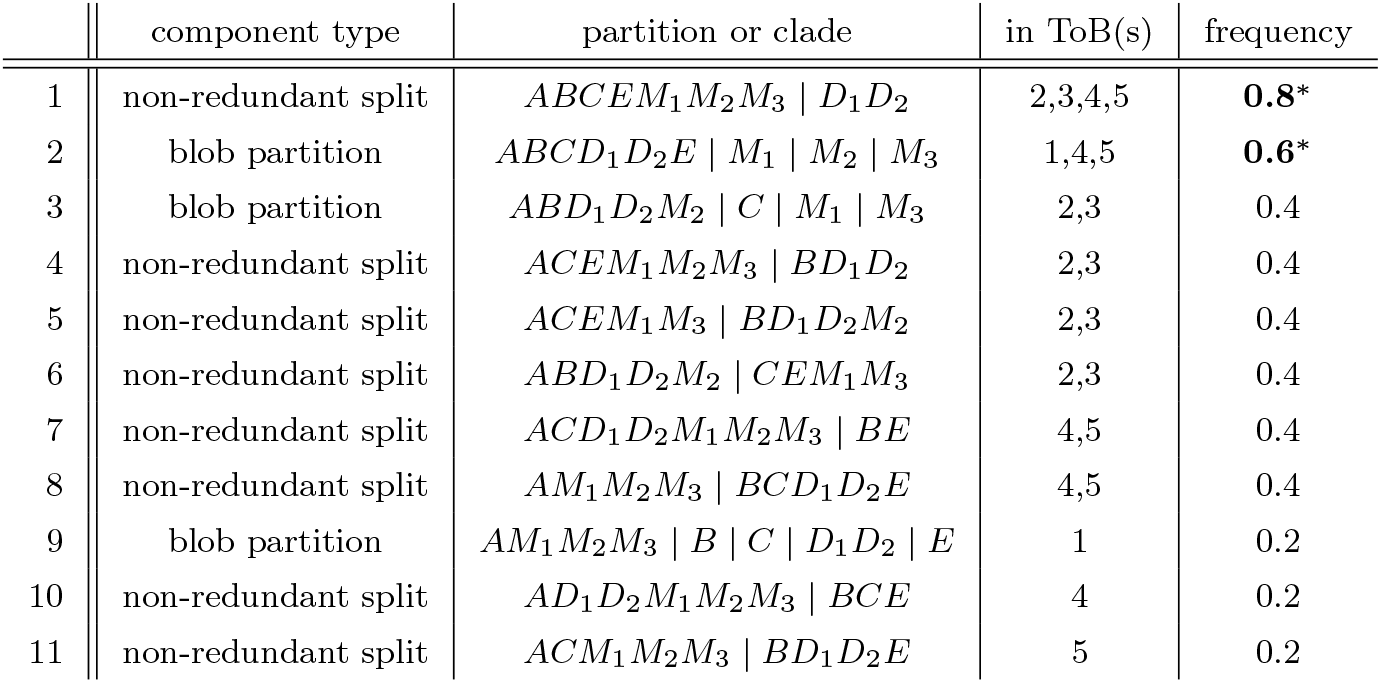

|  | component type | partition or clade | in ToB(s) | frequency |
|---|---|---|---|---|
| 1 | non-redundant split | ABCEM<sub>1</sub>M<sub>2</sub>M<sub>3</sub> \| D<sub>1</sub>D<sub>2</sub> | 2,3,4,5 | **0.8**<sup>\*</sup> |
| 2 | blob partition | ABCD<sub>1</sub>D<sub>2</sub>E \| M<sub>1</sub> \| M<sub>2</sub> \| M<sub>3</sub> | 1,4,5 | **0.6**<sup>\*</sup> |
| 3 | blob partition | ABD<sub>1</sub>D<sub>2</sub>M<sub>2</sub> \| C \| M<sub>1</sub> \| M<sub>3</sub> | 2,3 | 0.4 |
| 4 | non-redundant split | ACEM<sub>1</sub>M<sub>2</sub>M<sub>3</sub> \| BD<sub>1</sub>D<sub>2</sub> | 2,3 | 0.4 |
| 5 | non-redundant split | ACEM<sub>1</sub>M<sub>3</sub> \| BD<sub>1</sub>D<sub>2</sub>M<sub>2</sub> | 2,3 | 0.4 |
| 6 | non-redundant split | ABD<sub>1</sub>D<sub>2</sub>M<sub>2</sub> \| CEM<sub>1</sub>M<sub>3</sub> | 2,3 | 0.4 |
| 7 | non-redundant split | ACD<sub>1</sub>D<sub>2</sub>M<sub>1</sub>M<sub>2</sub>M<sub>3</sub> \| BE | 4,5 | 0.4 |
| 8 | non-redundant split | AM<sub>1</sub>M<sub>2</sub>M<sub>3</sub> \| BCD<sub>1</sub>D<sub>2</sub>E | 4,5 | 0.4 |
| 9 | blob partition | AM<sub>1</sub>M<sub>2</sub>M<sub>3</sub> \| B \| C \| D<sub>1</sub>D<sub>2</sub> \| E | 1 | 0.2 |
| 10 | non-redundant split | AD<sub>1</sub>D<sub>2</sub>M<sub>1</sub>M<sub>2</sub>M<sub>3</sub> \| BCE | 4 | 0.2 |
| 11 | non-redundant split | ACM<sub>1</sub>M<sub>2</sub>M<sub>3</sub> \| BD<sub>1</sub>D<sub>2</sub>E | 5 | 0.2 |

Since components 1 and 2 occur in a majority of input trees they occur in all consensus trees, whereas components 3, 5, and 6 are in none since they are incompatible with those components. Below are pictured the majority-rule consensus ToB (left) with components 1 and 2, and the three greedy consensus tree of blobs (right) obtained by greedily adding compatible components in the orders (4,8,11), (8,7,10), and (8,7,11) respectively to the majority-rule consensus ToB. All four trees are topologically distinct, and only the greedy consensus tree of blobs on the far right matches an input tree.

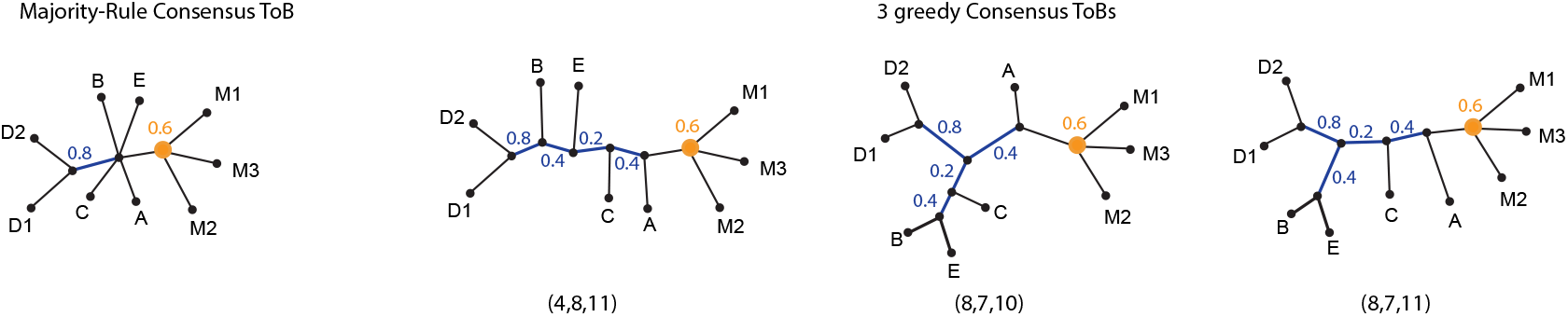

That traditional greedy consensus tree methods can produce multiple optima is well-known. Indeed, in this example the three greedy consensus trees are identical in topology to those greedy consensus ToBs depicted, though the support for splits might be higher since redundant splits also contribute to this value. For instance, because the split *D*_1_*D*_2_ |*ABCEM*_1_*M*_2_*M*_3_ occurs in all five input trees (sometimes as a redundant split, sometimes non-redundant), that split has support 1.0 in all three traditional consensus trees. The traditional majority-rule tree (not shown) is more resolved than the majority-rule consensus ToB, since the split *AM*_1_*M*_2_*M*_3_|*BCD*_1_*D*_2_*E* occurs in 3 of the trees, but only twice as a non-redundant split.

## 4 Consensus methods for binary semi-directed level-1 networks

The aim of this section is to provide additional context on the consensus methods for binary semi-directed level-1 networks introduced in the main text. However, we first show that such networks are encoded by their circular partitions, hybrid clades, and non-redundant splits. For simplicity of exposition, we assume that the input networks are binary, but this assumption can be relaxed. We only need that nodes within a cycle (blob) be binary, of degree 3. Internal nodes outside a cycle may have degree greater than 3, as explained after Definition 1.8.

### Proposition 4.1.

*Let N be a binary semi-directed level-*1 *network on taxon set X. Then N is uniquely determined by:*

i. *the circular partitions C*_*b*_ *induced by the blobs of N, together with the hybrid clade for each blob, and*
ii. *the non-redundant splits of N, i*.*e*., *edges not incident to any blob*.

*Proof*. Each blob *b* of *N* is associated with a circular partition *C*_*b*_ = (*A*_1_, …, *A*_*p*_) (the circular ordering of its blocks) and a hybrid clade, say *A*_*b*_.

Now, the set of blob partitions (the underlying blocks of the circular partitions) together with the non-redundant splits uniquely determines the tree of blobs *T* (*N*) of *N*.

Constructively, we can replace each blob vertex in *T* (*N*) by its corresponding cycle, attaching subnetworks according to the circular order, where the hybrid clade *A*_*b*_ determines the reticulation inside blob *b*.

Hence, the combination of circular partitions, hybrid clades, and non-redundant splits uniquely determines *N*.

The above proposition shows that binary semi-directed level-1 networks are fully encoded by circular partitions, hybrid clades, and non-redundant splits. We now use this decomposition to define a consensus method for such networks. The main text provides a detailed description of the method; here, we summarize its key steps and underlying construction. We say that a collection *C* of hybrid clades on a leaf set *X* is *level-*1*-compatible* (or *l1-compatible* for short) if there exists a level-1 network *N* on *X* such that every *C*_*i*_ ∈ *C* is a hybrid clade of *N*.

Let *N* = {*N*_1_, …, *N*_*k*_} be a collection of binary semi-directed level-1 networks on a common taxon set *X*. The *greedy consensus network* of *N* is obtained as follows:

- **Step 1: Greedy consensus tree of blobs**. For each *N*_*i*_, let *T*_*i*_ denote its tree of blobs. A greedy consensus tree of blobs *T* * is constructed from {*T*_1_, …, *T*_*k*_} using the greedy component set of Definition 3.1.
- **Step 2: Circular partition selection**. Each blob vertex in the consensus tree of blobs *T* * corresponds to a set of circular partitions observed in the input networks. We select the most frequent circular partition for each blob vertex. This step is straightforward and cannot generate conflicts, since the circular partition merely encodes the circular ordering of blocks in a blob partition.
- **Step 3: Selection of hybrid clades**. For each blob vertex *b* of the consensus tree of blobs *T* *, let *H* (*b*) denote the set of all blocks induced by the blob partition *ψ*_*b*_ associated with *b*. We seek to select one hybrid clade per blob vertex so as to maximize the total frequency, defined as the sum, over all blob vertices, of the frequencies of the selected hybrid clades across the input networks.

Under the assumptions that all blob vertices have degree at least 3 and arise from level-1 networks, a set of hybrid clades (one per blob vertex) is l1-compatible if and only if there exists a taxon *X* ∈ *X* that is excluded from all selected hybrid clades. This characterization yields an exact algorithm. When an outgroup is available, it plays the role of such a taxon *X*, and all candidate hybrid clades are mutually l1-compatible; thus, the optimal solution is obtained by selecting, independently for each blob vertex, the most frequent hybrid clade in *H*(*B*).

In the absence of an outgroup, we consider each taxon *X* ∈ *X* in turn. For a fixed *X*, we restrict *H*(*B*) to blocks that do not contain *X*, and select the most frequent remaining block for each blob vertex. This yields a candidate solution, which is scored by the sum of hybrid clade frequencies. The final solution is the one with maximum score over all choices of *X*. In case of ties, we break them by comparing the sum of frequencies restricted to blob vertices of degree greater than 4.

The correctness of Step 3 relies on the following characterization of l1-compatibility.

### Lemma 4.2.

*Let T* * *be the consensus tree of blobs obtained from a collection of binary semi-directed level-*1 *networks, and assume that all blob vertices have degree at least* 3. *Let S be a set containing one block per blob partition of T* *. *Then S is level-1-compatible if and only if there exists a taxon X* ∈ *X that is excluded from all blocks in S*.

*Proof*. We prove both directions.

- Suppose that there exists a taxon *X* ∈ *X* that is not contained in any block of *S*. We treat *X* as an outgroup, which induces a rooting of the tree of blobs *T* * (i.e., we root *T* * at the edge incident to *X*). Consider any blob vertex *b* of *T* *. Since all blob vertices have degree at least 3, removing *b* partitions the taxa into at least three parts corresponding to the incident edges. In the rooted tree, exactly one of these parts contains the outgroup *X*, while each of the other parts correspond to a clade (cluster) in the rooted tree. Because *X* is not contained in any block in *S*, the hybrid clade selected for each blob vertex must correspond to one of the parts not containing *X*. Hence every element of *S* is a cluster of the rooted tree *T* *. It is a standard property of rooted trees that any two clusters are either disjoint or nested. It remains to show that *S* is induced by some level-1 network. Starting from the rooted tree of blobs *T* *, we construct a rooted level-1 network *N* by expanding each blob vertex *b* as follows. For each blob vertex *b*, the chosen block in *S* corresponds to one of the parts of the partition induced by *b* that does not contain the outgroup *X*. We replace *b* by a simple level-1 structure (i.e., a single cycle) whose reticulation vertex is the parent of the chosen hybrid clade. Since the blocks in *S* are pairwise disjoint or nested, these local constructions are consistent across blob vertices and do not interfere with one another. Thus, performing this expansion independently for each blob vertex yields a well-defined level-1 network *N* that induces every block in *S* as a hybrid clade. Therefore, *S* is level-1-compatible.
- Assume that *S* is level-1-compatible. By definition, there exist some semi-directed level-1 network *N*, and hence a rooted level-1 network *N* ^+^, that induces every block in *S* as a hybrid clade. Since every rooted level-1 network is tree-child (i.e., every non-leaf vertex has a child that is not a reticulation; see [4]), every vertex *v* in *N* ^+^ admits a directed path to a leaf *l* that contains no reticulation vertices except possibly *v* itself (a tree path). In particular, there exists a root-to-leaf tree path, and let *X* be its endpoint. We claim that *X* is not contained in any hybrid clade. Suppose for contradiction that *X* lies in a hybrid clade. Then *X* is a descendant of some hybrid vertex *h*, meaning that every directed path from the root to *X* must pass through a reticulation vertex. This contradicts the existence of a root-to-*X* tree path, which contains no reticulation vertices. Hence, *X* is not contained in a hybrid clade.

This result justifies the rooting-based enumeration strategy used in Step 3, and shows that the algorithm explores exactly the space of feasible (i.e., l1-compatible) solutions.

### Remark 4.3.

While we focus here on a greedy consensus method for binary semi-directed level-1 networks, strict and majority-rule variants can be defined in a natural way by retaining components (circular partitions, hybrid clades, and non-redundant splits) that occur in all input networks or in a strict majority of them, respectively. For any given component type, the set of components occurring in a strict majority of the input networks minimizes the total symmetric-difference distance (see also Section 5) to the input component sets (the proof is similar to that of Proposition 3.10).

As in classical consensus settings, components that each occur in a strict majority of the input networks must co-occur in at least one input network and are therefore compatible. However, applying majority rule independently to different component types does not necessarily yield a valid level-1 network. For example, a hybrid clade may occur in a strict majority of the input network even though its associated blob partition or circular order does not. Consequently, the majority-supported components may not jointly define a fully specified level-1 network.

Rather than imposing additional resolution rules to enforce a valid network structure, we adopt a more transparent approach in our implementation by reporting the consensus network together with component-wise frequencies, allowing the user to assess support for each feature directly. We used this approach in the simulation study to define L1N-majority, by taking a greedy level-1 network consensus, and classifying each component (blob partition, circular order, hybrid clade, or split) as recovered if its frequency exceeds 50%. For splits, we use the standard support values, considering both redundant and non-redundant splits. In particular, the components retained by L1N-majority minimize the total symmetric-difference distance to the input component sets, although they need not define a valid level-1 network.

We next show that the greedy consensus network is well-defined.

### Lemma 4.4.

*Let N* = {*N*_1_, …, *N*_*k*_} *be a collection of semi-directed binary level-*1 *networks on a common taxon set X. Then the greedy consensus procedure produces a valid semi-directed level-*1 *network, possibly with some high-degree vertices*.

*Proof*. Each network *N*_*i*_ has an associated tree of blobs *T*_*i*_. Constructing a greedy consensus tree of blobs as defined in Section 3 produces an unrooted tree *T* * on leaf set *X*, even if ties occur and are resolved arbitrarily.

The tree *T* * may contain divergence vertices; these vertices are left unresolved (i.e., are not resolved into cycles).

Each blob vertex of *T* * is then resolved into a simple cycle by choosing a circular partition and a consistent hybrid clade. A consistent hybrid clade can always be selected: by rooting *T* * arbitrarily and directing edges from past to present, each blob partition has at least one block that is l1-compatible with all previously assigned hybrids. In practice, this block may have zero frequency as a hybrid clade in the input networks, but this does not affect validity. Assigning such a hybrid clade ensures that the resulting network is a well-defined semi-directed level-1 network.

Furthermore, notice that the greedy consensus network construction satisfies the same “majority-preserving” property as the consensus methods for trees of blobs discussed earlier.

### Remark 4.5.

If a binary semi-directed level-1 network *N* appears in more than half of the input networks, the greedy consensus procedure will reproduce *N* exactly. Indeed, all components of *N* (circular partitions, hybrid clades, and non-redundant splits) occur in a strict majority of inputs, so the consensus tree of blobs matches *T* (*N*), and the selection of circular partitions and hybrid clades reconstructs each cycle according to *N*. This property generalizes the trivial case where all input networks are identical.

## 5 Component-based distances between level-1 networks

Let *N*_1_ and *N*_2_ be semi-directed binary level-1 networks on the same taxon set *X*, without 2-cycles. As in the main text, we consider four types of components: blob partitions, circular orders, hybrid clades, and splits. For each component type *X* ∈ {blob partition, circular order, hybrid clade, split}, let 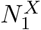 and 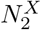 denote the corresponding sets of components in *N*_1_ and *N*_2_.

For a component *X*, we define the *normalized* symmetric-difference distance and *Jaccard distance* between *N*_1_ and *N*_2_ as

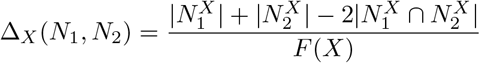

and

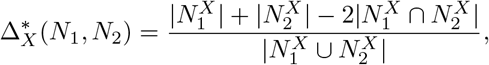

where *F* (*X*) *>* 0 is a normalization factor for component type *X*, chosen to scale the corresponding distance to approximately the same range as the other component-wise distances.

We define the overall symmetric-difference and Jaccard distances by averaging over the four component types:

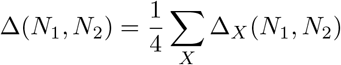

and

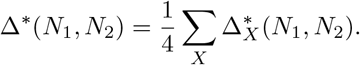

### Metric properties

For any component type *X*, both Δ_*X*_ and 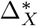 are metrics on finite sets of components. In particular, they are non-negative and symmetric, vanish if and only if the two component sets are equal, and satisfy the triangle inequality (for Δ_*X*_, this follows from the fact that the ordinary symmetric-difference distance

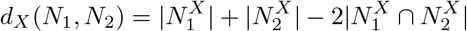

is a metric and that 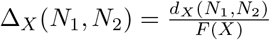 with *F* (*X*) *>* 0).

To establish that the overall distances are metrics on networks, we use the fact that semi-directed binary level-1 networks are uniquely determined by their component sets (Proposition 4.1). Consequently, if Δ(*N*_1_, *N*_2_) = 0, then the component sets agree for all four types and hence *N*_1_ = *N*_2_. The same argument applies to Δ*(*N*_1_, *N*_2_). Non-negativity, symmetry, and the triangle inequality follow from the corresponding properties of the component-wise metrics and the use of positive weights in the averages. Thus, both Δ and Δ* are metrics on the specified class of networks.

These distances are also straightforward to compute. Once the component sets have been extracted, their values can be obtained using standard set operations to determine the cardinalities of the sets, their intersections, and their unions. No optimization over network rearrangements is required.

### Application to the simulation study

In the simulation study, these pairwise distances are used to compare a consensus network with the input bootstrap networks. The symmetric-difference distance is normalized by choosing *F* (*X*) as the average number of components of type *X* across the model networks, thereby facilitating comparisons across component types. The Jaccard distance provides an alternative normalization based on the union of the component sets. For each model network and replicate, the corresponding distances are averaged over the 100 bootstrap input networks. The resulting summaries are then averaged over the 20 model networks and 10 replicates for each simulation setting. We denote these simulation-level averages by Δ(*C, X*) and Δ*(*C, X*), respectively, for consensus method *C* and component type *X*. Finally, the overall dissimilarity summaries are obtained by taking the unweighted mean over the four component types, as described in the main text.

For the empirical comparisons, we include both redundant and non-redundant splits, consider blob partitions and circular orders of size at least four, and include only hybrid clades occurring in blob partitions of size greater than four. These restrictions define the component sets used in the reported simulation summaries.

## 6 Support values in PhyloSummaries and PhyloNetworks

As indicated in the main text, our implementation of the consensus tree of blobs and consensus level-1 network further returns tables listing the support for various components in the consensus tree of blobs and level-1 network, with support measured by the proportion of input networks that have the component.

Additionally, traditional support for splits can be obtained by first extracting the bipartition splits from a network, which are those in the network’s tree of blobs using `treeofblobs` from PhyloNetworks, applied to each input network. These trees of blobs can then be summarized with `consensustree` from `PhyloSummaries`, to obtain the standard support of splits (regardless of whether they are redundant or not with a blob).

Other support values can be mapped onto a reference network *N*_0_, using existing tools in the PhyloNetworks package. These functions work more generally, in that they can take networks of any level, but require that input networks have inheritance probabilities on their hybrid edges. Given inheritance probabilities, we can define the major parent of each hybrid: the population (edge) from which most of its genome was inherited. We can then define a network’s major tree: the tree obtained by removing all minor relationships. The function `treeedges_support` calculates, for each edge in *N*_0_’s major tree, the percentage of input networks that also have this edge in their major tree. The function `hybridclades_support` calculates various measures of support for each hybrid node in *N*_0_, such as the support for each clade to be its (minor or major) sister.

## 7 A brief review of the HMS consensus of Huber et al. [8]

We refer the reader to Huber et al. [8] for a complete and rigorous description of the HMS consensus method and summarize the main ideas here.

The HMS consensus is based on an encoding of so-called 1-nested networks [10] (which coincide with level-1 networks in the binary case) as collections of ordered pairs (*S, H*) of subsets of the taxon set *X*, where *S* ≠ ∅ and *S* ∩ *H* = ∅. Specifically, each vertex *v* in a rooted phylogenetic network *N* ^+^ on *X* induces such a pair as follows: let *C* (*v*) denote the set of taxa descended from *v* (the set *C* (*v*) is also called the *hardwired cluster* of *v* [9]). Then *S* is defined as the subset of *C* (*v*) consisting of those taxa that can be reached from the root only via directed paths that pass through *v*, and *H* = *C* (*v*)*\S*.

These set pairs encode *compressed* 1-nested networks [5, 8], where “compressed” means that no vertex has out-degree 1; in particular, edges leaving reticulation vertices are contracted.

Huber et al. [8] further characterize which set pair systems can be represented by compressed 1-nested networks by deriving a set of technical conditions. This characterization forms the basis of their consensus construction.

Given a collection of *k* compressed 1-nested networks, the method proceeds as follows:

- Compute the set pair system for each network in the collection.
- For each pair (*S, H*), count its frequency across the collection, denoted by *c*(*S, H*).
- Construct a set pair system *P* consisting of all pairs (*S, H*) satisfying
  - 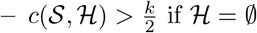, and
  - 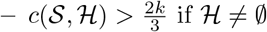.

The resulting set pair system *P* is 1-nested compatible, and hence can be represented by a unique compressed 1-nested network, referred to as the HMS consensus network.

This construction can be viewed as a variant of a majority-rule consensus: set pairs corresponding to tree-like evolution (*H* ≠ ∅) are retained under a strict majority threshold, while those corresponding to reticulation structure (*H* = ∅) must satisfy a stricter two-thirds threshold. As a consequence, the resulting consensus network may contain soft polytomies whenever no set pair exceeds the required frequency threshold. In particular, the more stringent 2*/*3 criterion for reticulate structures seems to suppress weakly supported hybrid signals, often leading to a higher degree of unresolved structure compared to our L1N-majority approach.

### Implementation

As Huber et al. [8] did not provide a practical implementation of the HMS consensus method, we developed a basic Perl implementation, available on the GitHub repository associated with this manuscript (https://github.com/HectorBanos/NetworkConsensus). Since our input networks are not compressed, we modify the extraction of set pairs as follows: for each network, we count each pair only once, even though a reticulation vertex and its child may induce the same set pair.

In addition, our implementation outputs non-compressed networks by ensuring that reticulation vertices have out-degree one. In this case, a reticulation vertex and its child induce the same set pair, but no additional set pairs are introduced. This adjustment facilitates a fair comparison with our own consensus method. However, the original HMS consensus network can be recovered from our output by contracting all edges leaving reticulation vertices.

Finally, a list of network weights can also be provided. By default, all input networks are weighted equally.

## 8 A brief review of the HWC consensus of Nielsen et al. [16]

The HWC (hardwired cluster) consensus, implemented in AdmixtureBayes [16], provides a summary of a collection of rooted phylogenetic networks based on their hardwired clusters. We briefly outline the main idea here and refer the reader to the original sources for further details.

Given a rooted phylogenetic network on a taxon set *X*, each vertex *v* induces a *hardwired cluster*, defined as the set *C* (*v*) of taxa descended from *v* [9]. Thus, each network can be represented by its collection of hardwired clusters. For a collection of *k* input networks, the frequency of each cluster is computed across the sample, counting each cluster at most once per network (even if it is induced multiple times; for instance, a reticulation vertex and its child define the same hardwired cluster). Given a threshold *α*, the consensus retains all clusters that occur in strictly more than an *α* proportion of the input networks. The resulting set of clusters is then organized into a network by constructing its Hasse diagram under set inclusion, yielding the HWC consensus network.

This construction can be viewed as a cluster-based majority consensus, analogous to classical consensus methods for trees. However, because the output is necessarily a *cluster network* [9], the method implicitly assumes that the input networks lie in this class. Since not all rooted phylogenetic networks are cluster networks, the HWC consensus does not, in general, satisfy idempotence. In particular, there exist networks *N* such that applying the HWC consensus to a collection of identical copies {*N, N*, …, *N*} yields a network that is not isomorphic to *N*. In contrast, the L1N consensus guarantees recovery of *N* when it has a strict majority among the inputs (see above). Furthermore, HWC does not guarantee a level-1 output, even when the inputs are level-1. In our experiments with an *α* value of 0.5, this occurred rarely. However, it occurs much more frequently with higher *α* values.

### Implementation

We tested the implementation of the HWC consensus provided in AdmixtureBayes [16]. However, this implementation outputs only graphical representations of the consensus network (plots in pdf format) and does not provide additional metadata, such as an extended Newick representation, which limits its direct use in downstream analyses. Moreover, the required input format for networks is not based on extended Newick, necessitating an additional reformatting step for users. To facilitate comparison with our method and enable reproducibility, we therefore also implemented a basic version of HWC in Perl (https://github.com/HectorBanos/NetworkConsensus), which accepts input networks and outputs the resulting consensus network in extended Newick format. As with the HMS consensus, we output non-compressed networks by ensuring that reticulation vertices have out-degree one. Finally, a list of network weights can be provided, but by default, all input networks are weighted equally.

## 9 Treatment of non-binary or non-level-1 consensus networks

As described in the main text, for each consensus method considered in this manuscript, we recorded the induced blob partitions, circular orders, hybrid clades, and splits. All L1N methods produce binary level-1 networks, whereas HMS and HWC may produce non-binary networks. Moreover, HWC may also produce networks that are not level-1.

If an HMS or HWC consensus network induced a blob partition associated with a cycle containing high-degree vertices, such that the circular order of taxon blocks was not uniquely determined, we always scored the circular order as incorrect. The blob partition and hybrid clade were evaluated independently and scored as correct or incorrect depending on whether each matched the corresponding blob partition or hybrid clade, respectively, in the reference bootstrap network or model network.

By contrast, if an HMS or HWC consensus network contained a high-degree vertex not belonging to any cycle (i.e., a classic multifurcation), we scored the associated blob partition, circular order, and hybrid clades as incorrect.

For non-level-1 networks produced by HWC, we randomly deleted reticulation edges from high-level blobs to obtain a level-1 network. In the few cases where this was necessary, removing a single reticulation edge was sufficient to obtain a level-1 network. We then used the blob partitions, circular orders, hybrid clades, and splits induced by the resulting network for evaluation.

## 10 Simulation settings

In this section, we present the simulation settings used in the work.

### 10.1 Networks

For this simulation, 20 rooted metric networks on 20 taxa were generated using a Birth-Death process (BD) that accounts for hybridization, as implemented in the R package SiPhyNetwork [11]. We generated networks using this process until we obtained 20 networks that are level-1 and either had at least three 4-cycles or one 6-cycle. Any network not satisfying these conditions was discarded. Note that such networks can contain 2- and 3-cycles; nonetheless, these are not identifiable by any of the inference software used in this work. Additionally, under the simulation settings considered here, the hybrid node in 4-cycles is also not identifiable by any of the software evaluated. As a result, all analyses exclude the inference of 2- and 3-cycles, as well as the identification of hybrid nodes in 4-cycles. Yet data was simulated from networks with 2- and 3-cycles.

To reduce the level of incomplete lineage sorting, each edge (hybrid or tree) in the species network with a length of less than 0.4 coalescent units was repeatedly resampled from an exponential distribution with mean 1 until its length was at least 0.4 coalescent units. Similarly, to avoid a very weak hybridization signal, any hybridization parameter below 0.3 (and, consequently, above 0.7 for the complementary inheritance probability) was repeatedly resampled from a uniform distribution on the interval [0.3, 0.7].

All 20 network topologies are depicted in Figures S8 and S9.

**Figure S8:**
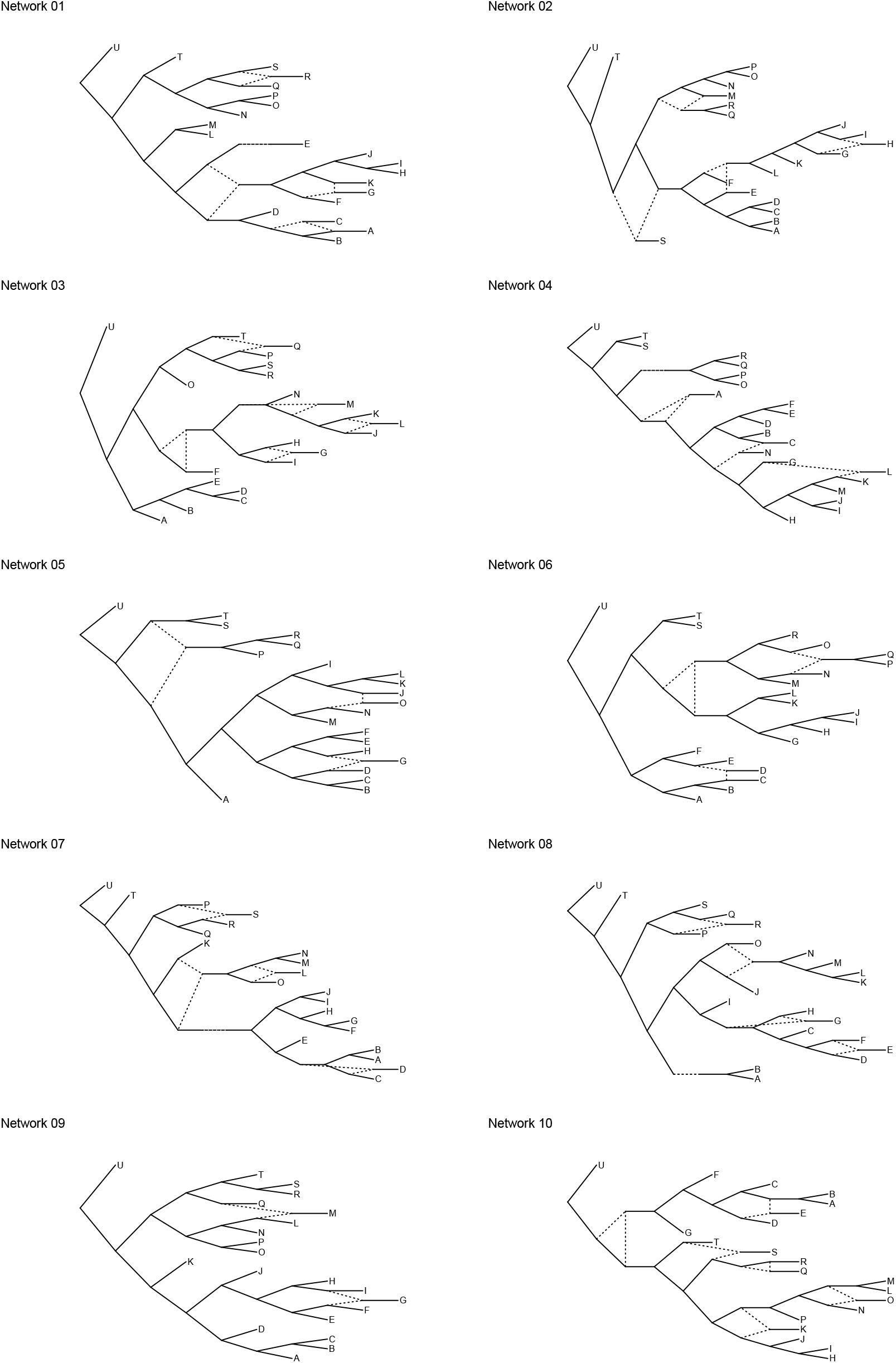
The first ten of the twenty level-1 network topologies obtained from `SiPhyNetwork`.

**Figure S9:**
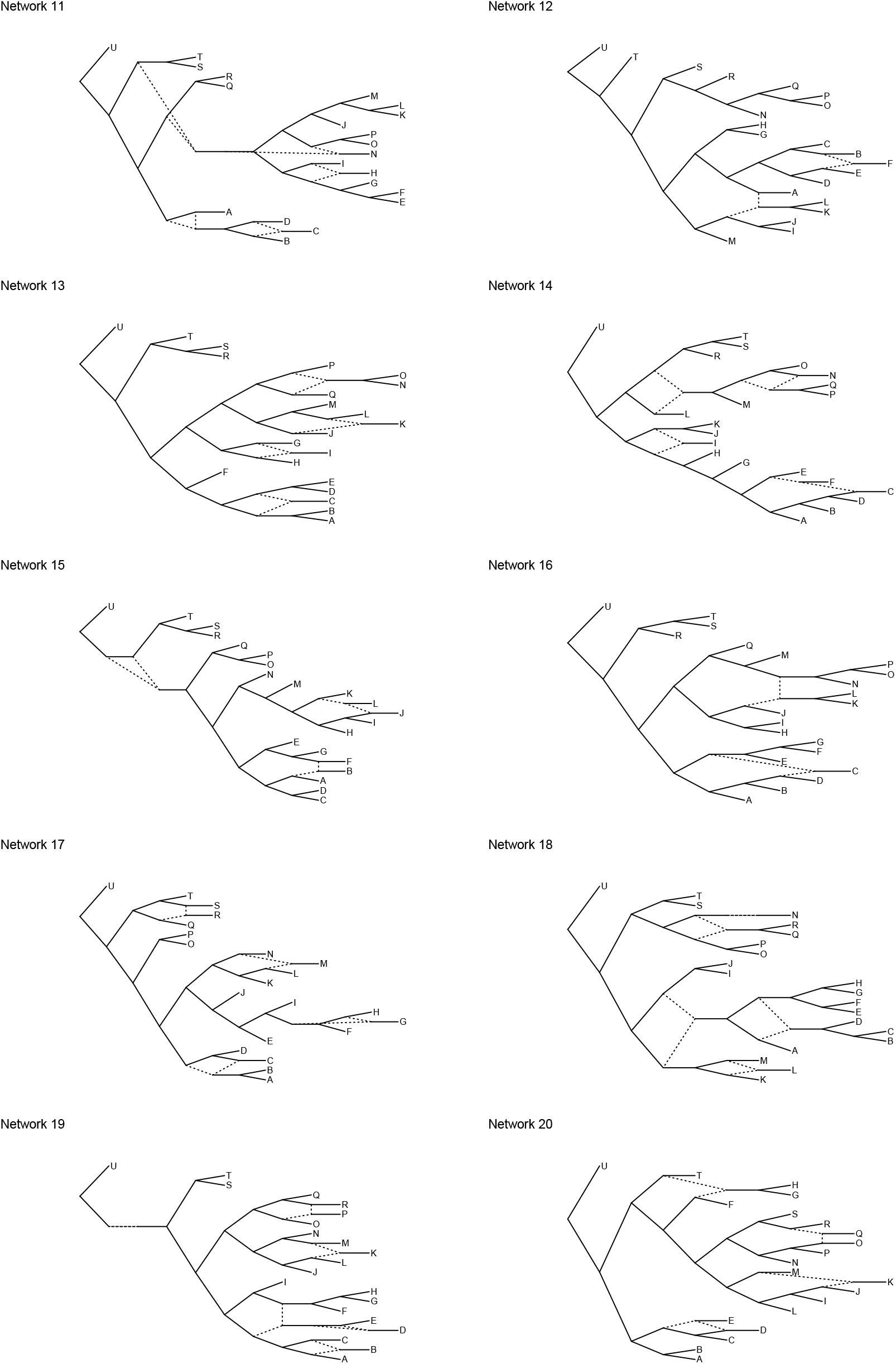
The second ten of the twenty level-1 network topologies obtained from SiPhyNetwork.

**Figure S10:**
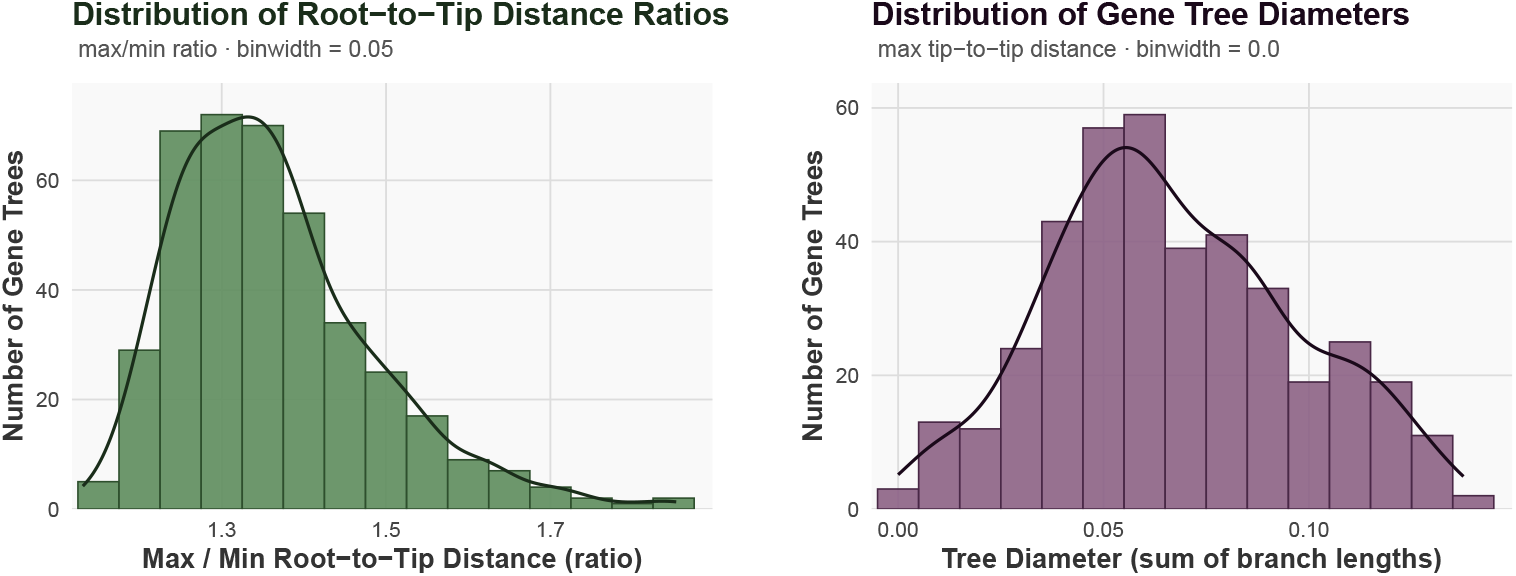
Left: Histogram of the max/min root-to-taxon distance ratio after rescaling the edge lengths of 400 gene trees. Right: Histogram of the resulting tree diameters for the same set of 400 rescaled gene trees.

### 10.2 Gene trees

For each network, we simulated 400 gene trees under the Network Multispecies Coalescent using the PhyloCoalSimulations package in Julia [6], using the default parameters.

The edge lengths of the gene trees were first rescaled so that they match key summary statistics observed in the 7 Papionini species analyzed in [12] and whose data were produced in [17], as described below.

To break ultrametricity in the gene trees, all edge lengths were multiplied by values independent drawn from a log-normal distribution with mean 1 and standard deviation 0.127. This standard deviation was chosen so that, on average, the ratio between the maximum distance from the root to a taxon and the minimum distance from the root to a taxon is approximately 1.2. This value was the median ratio from the Papionini dataset, where the trees were rooted using the midpoint method. Figure S10 (left) shows the distribution of this ratio for one set of rescaled sampled gene trees for a single network.

Next, each tree was rescaled so that its diameter followed the same distribution as the diameters observed in the Papionini dataset, where the distribution was truncated at diameter values of 0.15. Figure S10 (right) shows the distribution of diameters from one set of rescaled sampled gene trees for a single network (with median ~0.06).

Then, an outgroup was appended to each gene tree. The branch length from the root to the outgroup was set equal to the diameter of the original tree, while the branch length from the root to the root of the ingroup gene tree was set to half of the tree diameter. Therefore, there was no ILS between the original 20 taxa and the appended outgroup, as this was attached after gene tree generation under the network multispecies coalescent (NMSC).

### 10.3 Sequences

For each gene, we simulated multiple sequence alignments using `AliSim` (`--alisim` tool in `IQ-TREE`) [15].

For each gene, the sequence length was sampled from a Gamma distribution shifted to have a minimum approaching 300bp, a mean of 600bp, and a moderate right skew. Figure S11 shows the distribution of sequence lengths.

Sequences were generated under the GTR substitution model with relative rate parameters

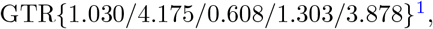

base frequencies

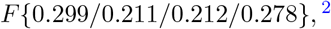

and a gamma shape parameter {*G* 1.518}.

**Figure S11:**
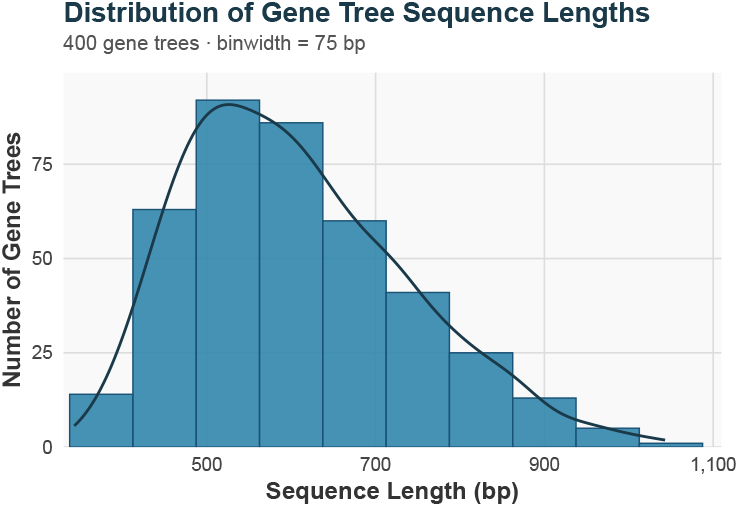
Histogram showing the sequence length distribution from all 400 simulated sequence alignments.

These parameter values were estimated from 10 primate genes obtained from www.ensembl.org: *RAG1, RAG2, IRBP, BRCA1, APOB, VWF, GHR, MC1R, TYR*, and *MITF*. Specifically, for each gene, we used `IQ-TREE3` [18] to estimate the substitution rate matrix, base frequencies, and the *α* shape parameter of the Γ(4) rates model. The final parameter values were obtained by taking the mean of the estimates across the 10 genes.

Figure S12 shows the estimated substitution rates, base frequencies, and *α* parameters for all 10 genes, respectively.

Gaps were incorporated using the mimic option in `AliSim`, with the Papionini dataset [17] provided as a template.

**Figure S12:**
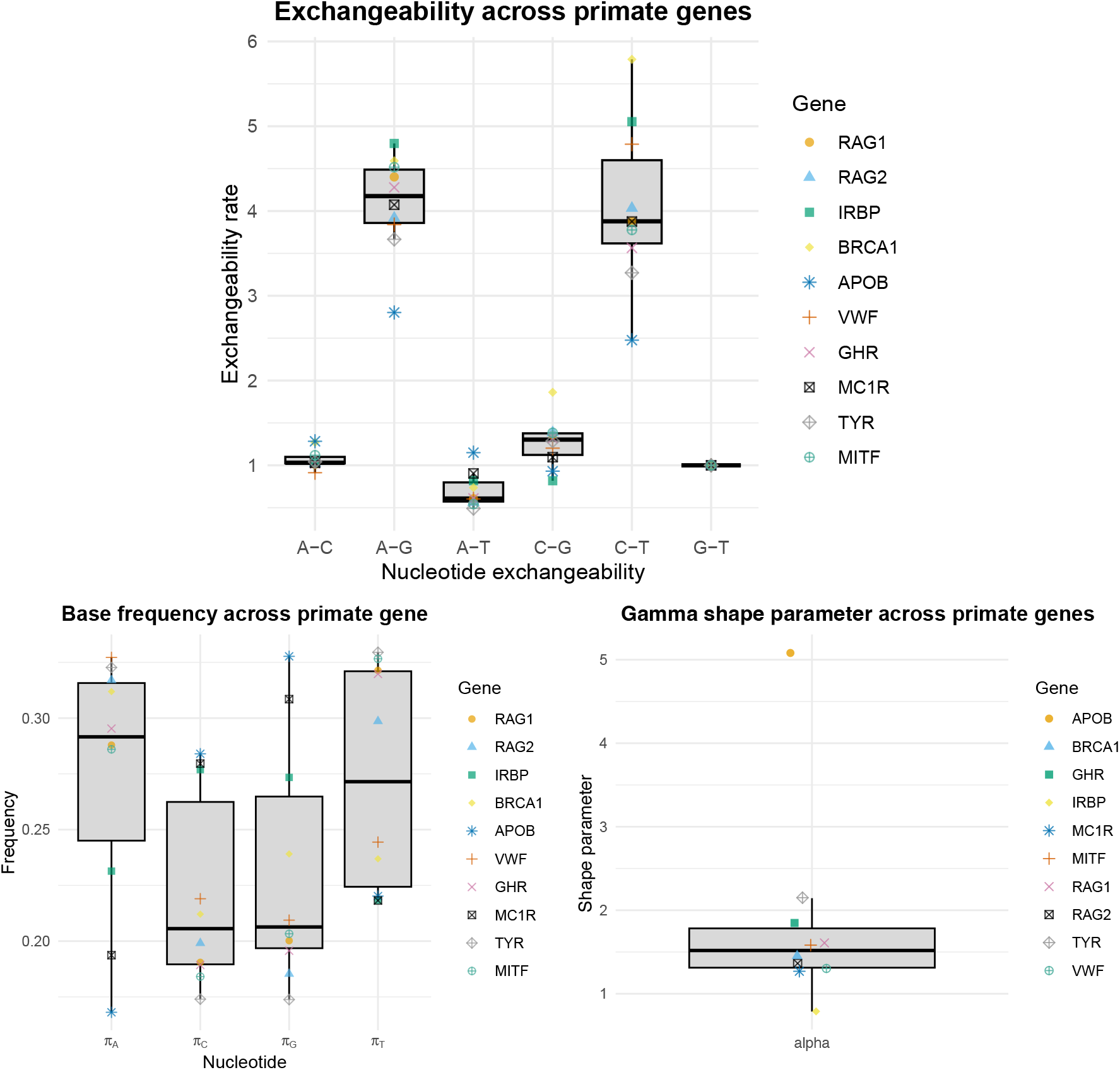
Estimates of substitution model parameters across the 10 primate genes. Distribution of the estimated substitution rates (top), base frequencies (bottom left), and *α* parameters (bottom right). The parameter values used in the simulations correspond to the median estimates across genes.

## 11 Supplementary results and figures for the simulation study

In this section, we report additional results from the simulation study. Table S1 summarizes the average number of blob partitions and the average number of blob partitions of size at least 5 inferred by each method in both the ‘Full’ and ‘Third’ gene settings (see main text for details).

**Table S1:** Average number of blob partitions of size at least four and blob partitions of size greater than four per method and setting, over all replicates.

|  | Setting | TRUE | Squirrel | HMS | HWC | L1N-majority | L1N-unrooted | L1N |
|---|---|---|---|---|---|---|---|---|
| Blob partitions (≥ 4) | Full | 3.35 | 2.94 | 1.78 | 1.43 | 2.56 | 3.10 | 3.02 |
| Blob partitions (≥ 4) | Third | 3.35 | 3.19 | 1.27 | 1.44 | 2.15 | 3.34 | 3.10 |
| Blob partitions (> 4) | Full | 1.7 | 1.14 | 0.90 | 0.95 | 1.10 | 1.29 | 1.25 |
| Blob partitions (> 4) | Third | 1.7 | 1.21 | 0.70 | 0.79 | 0.89 | 1.10 | 1.14 |

Figures S13 and S14 complement the figures shown in the main text. They show the average distance of the consensus networks to the input networks (Fig. S13) as well as the accuracy of the consensus networks with respect to the model networks (Fig. S14).

**Figure S13:**
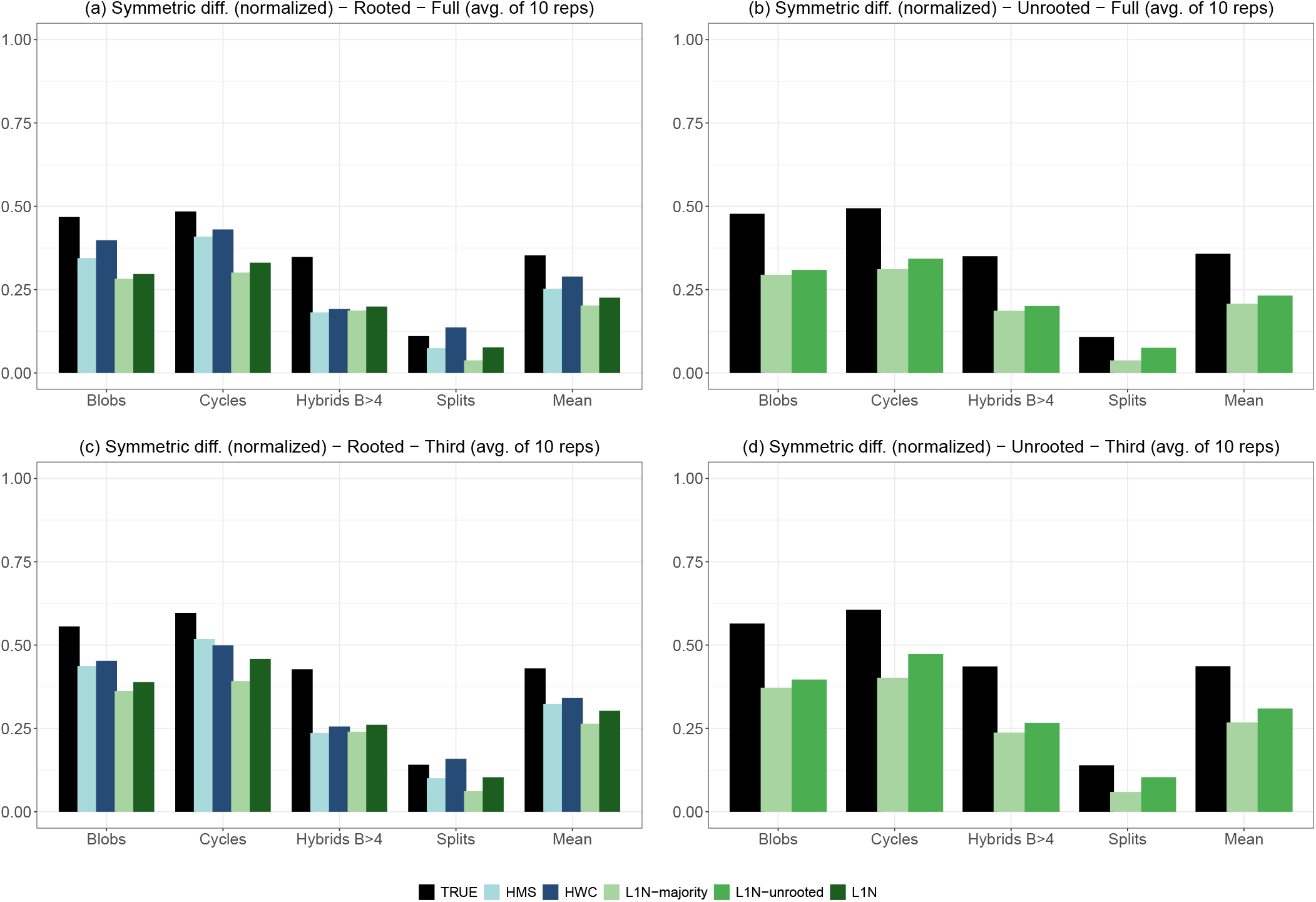
Average symmetric distance to the input networks. For each method and setting, we provide the average symmetric distance to the input networks with respect to blob partitions (Blobs), circular orders (Cycles), hybrid clades in blobs of size greater than 4 (Hybrids B*>*4), and all splits (both redundant and non-redundant), as well as the mean of these four measurements. (a) ‘Full’ genes and ‘Rooted’ setting; (b) ‘Full’ genes and ‘Unrooted’ setting; (c) ‘Third’ of each gene and ‘Rooted’ setting; (d) ‘Third’ of each gene and ‘Unrooted’ setting. For the ‘Rooted’ setting we compare the HMS, HWC, and L1N consensuses. Moreover, we provide the results of L1N-majority, in which only components (e.g. blob partitions) with over 50% support are retained (standard support is applied to splits). In the ‘Unrooted’ setting, HMS and HWC consensuses are not applicable; we provide the results of L1N-unrooted, in which neither Squirrel nor L1N is given knowledge of the outgroup, and L1N-majority, which is also non-informed about the outgroup in that case. For all four settings, we provide the average symmetric distance between the input networks and the ‘TRUE’ model network that was used to generate the data (black bars). This provides a basis for comparison, particularly in the ‘Unrooted’ setting, since we would expect any consensus method to be closer to the input networks than the ‘TRUE’ model is. See main text for further details and explanations.

**Figure S14:**
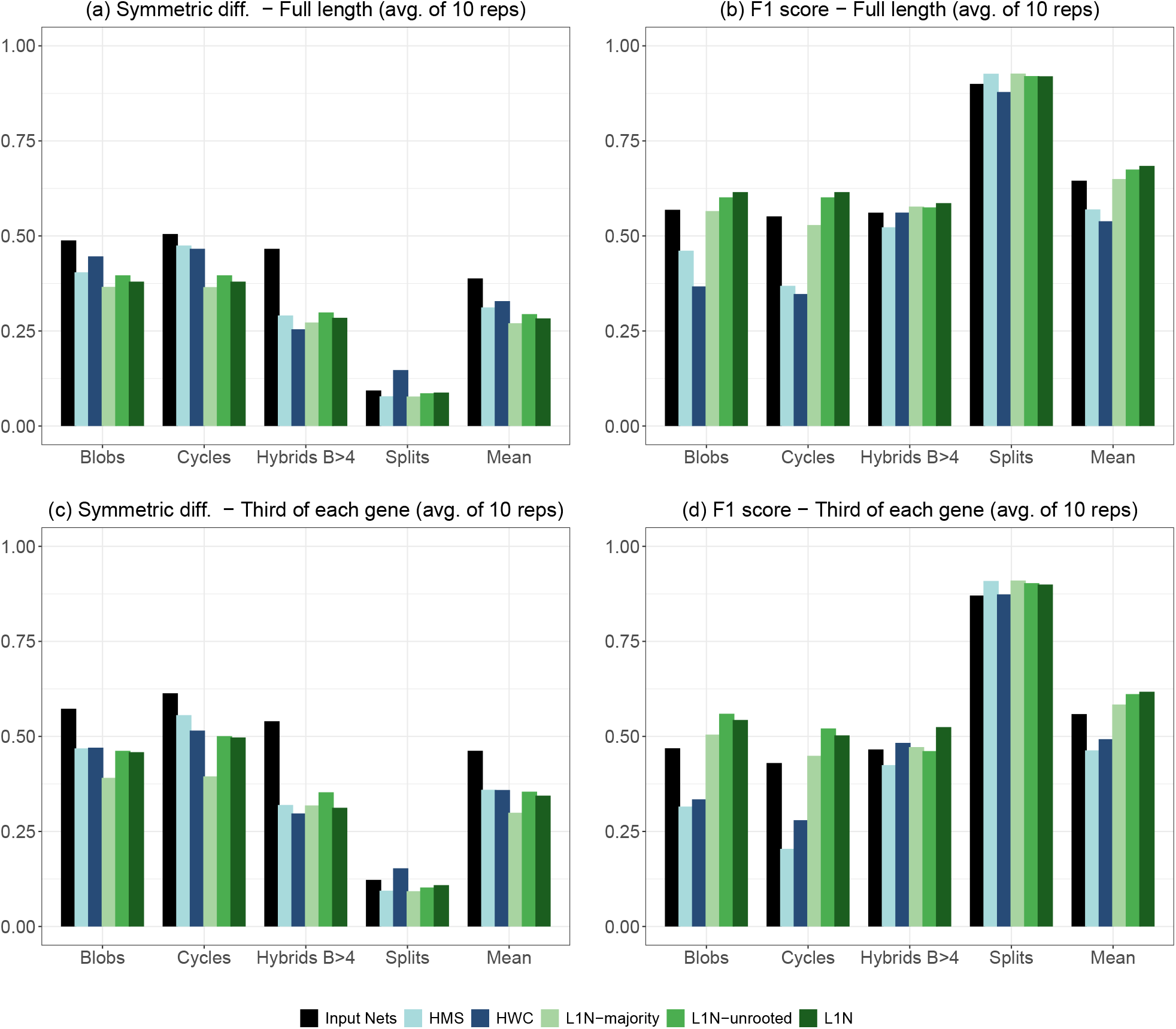
Accuracy of the consensus networks with respect to the true model network. For each method, we provide the symmetric difference (lower is better) and the F1 score (higher is better) between the consensus network and the true model network used to generate the data, with respect to blob partitions (Blobs), circular orders (Cycles), hybrid clades in blobs of size greater than 4 (Hybrids B*>*4), and all splits (both redundant and non-redundant), as well as the mean of these four measurements. All values are averaged over 10 replicates. (a) Symmetric difference, ‘Full’ genes; (b) F1 score, ‘Full’ genes; (c) Symmetric difference, ‘Third’ of each gene; (d) F1 score, ‘Third’ of each gene. We compare the HMS, HWC, and L1N consensuses, which are informed about the outgroup. Moreover, we provide the results of L1N-majority, in which only components (e.g. blob partitions) with over 50% support are retained (standard support is applied to splits), and L1N-unrooted, in which neither Squirrel nor L1N is given knowledge of the outgroup. For comparison, we also provide the average symmetric difference and F1 score between the input networks and the true model network (‘Input Nets’, black bars), which indicates how much a consensus method improves on the individual input networks. See main text for further details and explanations.

## 12 *Leopardus* information

Code for and information about reproducing the bootstrap replicates and subsequent analysis is available at https://github.com/HectorBanos/NetworkConsensus/blob/main/Leopardus/.

## 13 Supplementary figures for the HIV analysis

This section reports the two additional HIV analyses referred to in the main text: the consensus of the semi-directed networks returned by Squirrel in its unrooted setting (Figure S15), and the full pipeline repeated with subtypes J and H excluded (Figure S16). Both use the same 1,000-replicate resampling scheme and the same support measures as the main analysis.

**Figure S15:**
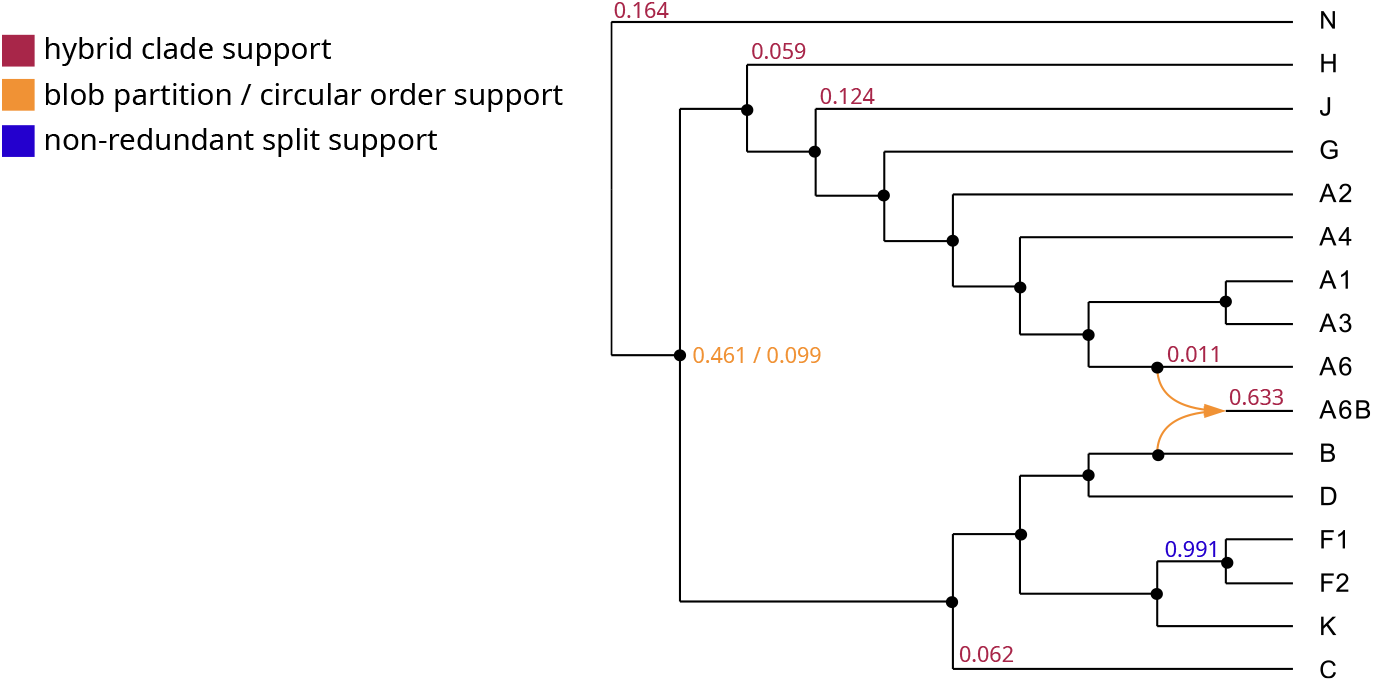
L1N consensus network for the HIV-1 group M subtypes on 1,000 semi-directed networks inferred by Squirrel [7] in its unrooted setting, i.e. without designating N as outgroup. The topology is the same as that of the rooted L1N consensus in the main text, except that N is a pendant clade of the blob rather than the outgroup, and is inferred as the hybrid in 16.4% of the input networks. Support values follow Figure 10 in the main text, except that no cluster support is reported, as hardwired clusters are defined only for rooted networks.

**Figure S16:**
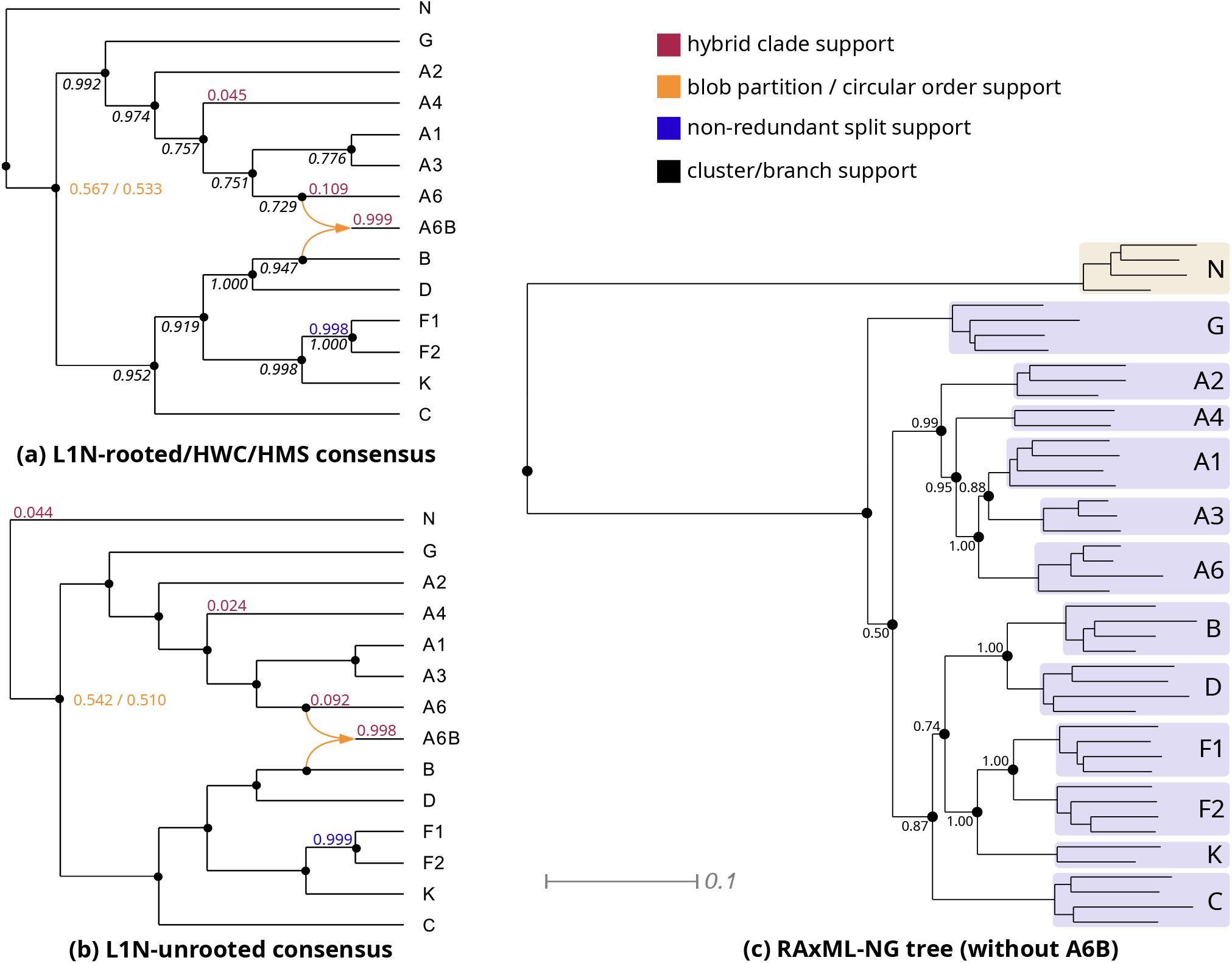
Results of the full analysis of the HIV-1 group M subtypes excluding J and H: the consensus network from the 1,000 rooted Squirrel [7] networks, which L1N, HWC and HMS all return (a), the L1N consensus network on the 1,000 networks inferred in the unrooted Squirrel setting (b), and the RAxML-NG [13] tree inferred from the reference alignment with the A6/B recombinant removed (c). Support values follow Figure 10 in the main text, except that no cluster support is reported in (b), as hardwired clusters are defined only for rooted networks.

## Footnotes

1 Corresponding exchangeabilities of A-C, A-G, A-T, C-G, C-T, and G-T, respectively.

2 Corresponding frequencies of A, C, G, and T, respectively. We used the normalized means from the Primate data.

